# Evolutionary dynamics of pigmentary grey and non-iridescent structural blue colouration in Tanagers (family: Thraupidae)

**DOI:** 10.1101/2023.09.07.556662

**Authors:** Frane Babarovic, Christopher R Cooney, Thomas Guillerme, Nicola J Nadeau, Gavin Huw Thomas

## Abstract

Birds are one of the most colourful animal groups in the world and there are multiple ways by which they achieve this feature. Mechanisms of colour production range from pigmentary (pigment deposition) to structural (nanostructural arrangements), or the combination of both. Despite the huge *breadth* of colour gamut, the basic components of feathers are shared across all of them (keratin, air and presence of pigments in accordance with the colour produced). It has been shown that in some instances, colour evolution between pigmentary and structural colours can proceed by rearrangement of the nano-structural elements of feathers. Here, we investigated evolutionary transitions between pigmentary grey and non-iridescent structural blue. We focus on the Thraupidae (tanagers and allies) that display a variety of blues and greys including a potential transition state that we refer to as slate. We used digitally calibrated images of birds to quantify colour and determine the distinctiveness of slate colour in colourspace. Following, we identify the most likely pathway for the evolution of the colour blue: from grey via slate colour. Our research reveals a new pathway in the evolution of blue colour.

## 3.2 Introduction

The array of colours in a bird’s plumage are produced either by pigment deposition in their plumage (pigmentary colours), precisely arranged structural elements of feathers (structural colours) or as a combination of both (Hill & McGraw, 2006; Shawkey & D’Alba, 2017). In melanin-based pigmentary colour, the pigment melanin is produced, transported, and stored in organelles called melanosomes (D’Alba & Shawkey, 2019). Melanin based colouration includes black, brown, and grey hues with each one having a characteristic melanosome shape (Babarović et al., 2019; Li et al., 2010). In structural colours, precisely arranged elements of feather nanostructure are responsible for colour production (Hill & McGraw, 2006). Non-iridescent structural colours in birds are produced by a keratin and air matrix (spongy layer) placed within the feather barb (Hill & McGraw, 2006; Prum et al., 1998; Shawkey & Hill, 2006). A melanosomes layer beneath the colour producing nanostructure is essential in colour production by absorption of all backscattered light (Shawkey & Hill, 2006). Non-iridescent structural colours encompass blue, violet and ultraviolet (UV) hues (Saranathan et al., 2012; Shawkey & D’Alba, 2017).

Evolutionary transitions between pigmentary and structural colours in birds’ plumage have been described previously (Doucet et al., 2004; Driskell et al., 2010; Shawkey et al., 2006). Due to the similarity of elements involved in the production of each colour, i.e. keratin, melanin and air gaps, it has been proposed that evolutionary transitions happen through rearrangements of feather nanostructure (Prum, 2006). This hypothesis has been demonstrated in the case of the transition between matte black and iridescent colouration in grackles and allies (Icteridae), where light-scattering melanosomes are organized in an ordered layer on the edges of the barbules (Shawkey et al., 2006). Other structural similarities between certain pigmentary and structural colours exist that indicate their potential transitions. For example, the black plumage of some fairy wrens has been found to have a spongy layer indicative of non-iridescent structural colours, but with additional melanosomes incorporated into the feather barb. These additional melanosomes prevent colour production by the spongy layer, leading to a black colour (Doucet et al., 2004; Driskell et al. 2010).

Recently, it has been suggested that non-iridescent structural blue colours evolve via a transition from grey, based on the similarity in the structural components of the feather (Babarović et al., 2019). The investigation of melanosome shape showed an overlap between those involved in grey colour production and melanosomes placed underneath the colour producing nanostructures in non-iridescent structural colour. The mechanistic shift needed for this colour evolution would involve the rearrangement of melanosomes from grey colouration around the central feather shaft and subsequent development of the spongy layer. The potential mechanism for this transition is also supported by evidence from research into the colour producing spongy layer in feather barbs (Saranathan et al., 2012). A broad analysis of non-iridescent structural colours across different bird clades indicated that a colour category broadly defined as blue-grey (or slate) possesses “rudimentary or weakly nanostructured” feathers (Saranathan et al., 2012). This indicates that this colour state has a poorly developed spongy layer and could potentially be a transition between grey and non-iridescent structural blue colour (which has the spongy layer fully developed) in bird’s plumage.

To explore the evolutionary dynamics and potential transition between pigmentary grey colour and the non-iridescent structural blue colour, we focused on the passerine family Thraupidae, the tanagers. Thraupidae have diverse plumage including pigmentary grey, non-iridescent structural blue as well as a wide range of slate (blue-grey) colours. This diversity of colour distribution across the phylogeny makes them an ideal study system for addressing the origins and potential dynamics that lead to the proliferation of blue colours among birds. We collected data from handbook descriptions of bird colouration as well as using quantitative measure of colour (cone catch values) from digital images of birds from museum specimens. We tested: 1) whether slate is an intermediate colour state between pigmentary grey and non-iridescent structural blue using visual modelling in avian colourspace, and 2) whether an evolutionary pathway to non-iridescent structural blue in Tanagers started with grey and proceeded through slate.

## 3.3 Materials and methods

### 3.3.1 Data collection

We collected plumage colouration data from written descriptions of the plumage colour from Birds of the World (Billerman, et al. 2022) and from digitally calibrated images of study skins from the bird collection from Natural History Museum at Tring.

### 3.3.2 Plumage Colour descriptions

We classified colour in ten patches (crown, nape, mantle, rump, throat, breast, belly, tail, wing coverts and wing primaries/secondaries) for both males and females for 174 species based on verbal descriptions from the Identification paragraph in the Birds of the World (Supplement data: Table S1). Using these descriptions, we coded each plumage patch as blue, slate and grey (Supplement data: Table S1). Due to human induced bias in recognising each colour, we verified the verbal description of colour from both the available images and video recordings from Birds of the World (Billerman, et al. 2022). Full colour descriptions and the subsequent assigned colour categories are reported in the Additional table 2. (Males scoring) and Additional table 3. (Females scoring) (https://figshare.com/s/1110fce894e65a69c329). In the Supplement data (Table S1) is the abbreviated version of these tables with only plumage patches that have either grey, slate or blue reported.

### 3.3.3 Plumage Colour measurements

To quantify chromatic variation of colour (hue and saturation), calibrated digital images of study skins from Cooney et al. (2022) were used and colour was quantified from them. Briefly, each bird species was photographed six times: from dorsal, lateral, and ventral angel and with two filters (one permitting UV wavelengths and human visible wavelengths). An average of three male and three female study skins were photographed per species. For all imaging, a Nikon D7000 digital single-lens reflex camera was used (for details of all technical specificity, see Cooney et al. (2019)).

Next, all digital images were linearized and converted to .TIFF files (Coffin, D., 2016). The images were normalized by reference to grey standards with known reflectance (Troscianko & Stevens, 2015). Plumage colouration was measured for seven selected body regions: crown, nape, mantle, rump, throat, breast, and belly. On these regions a predominance of a single colour is more likely than in other plumage patches. On every photo, polygons used to mark plumage patches with a custom IMAGE J script and RGB values were extracted for both the human-visible and UV range (Rueden et al., 2017).

Finally, by using already available tools in the IMAGE J Calibration and Analysis Toolbox (version 1.22), mapping functions were applied using methods described in Troscianko and Stevens (2015) to convert all RGB values to raw cone-catch values (u, s, m, l) adjusted to avian colour vision. We used u, s, m and l values for further analysis because they account for avian spectral sensitivities to ultraviolet (u), shortwave (s), mediumwave (m), and longwave (l) light (Stoddard & Prum, 2011). For each sex, average patch values were calculated across all specimens as a species-level measurements.

## 3.4 Analysis

### 3.4.1. Distinctiveness analysis

The extracted data from digital images were filtered for only those plumage patches that scored either blue, slate or grey in our written description dataset and for seven plumage patches (crown, nape, mantle, rump, throat, breast and belly) (Supplement data: Table S1). We focused on these seven patches because the marked regions on the digital image data for other patches typically included only a small proportion of the focal colour resulting in patch colour measurements that would provide inconsistent and unreliable measures of the presence of grey, slate, and blue. These values were then visualised in an avian tetrahedral colourspace, using methods from Stoddard & Prum (2008) implemented in the R package pavo (Maia et al., 2019; R Core Team, 2021) (Fig. 3.1, a, b, c). In tetrahedral colourspace every data point is represented by four values (ultraviolet cone – u, short-wavelength cone – s, medium-wavelength cone – m, long-wavelength cone -l) that are equivalent to how much each colour stimulates each cone in the bird’s retina.

**Figure 3.1.**
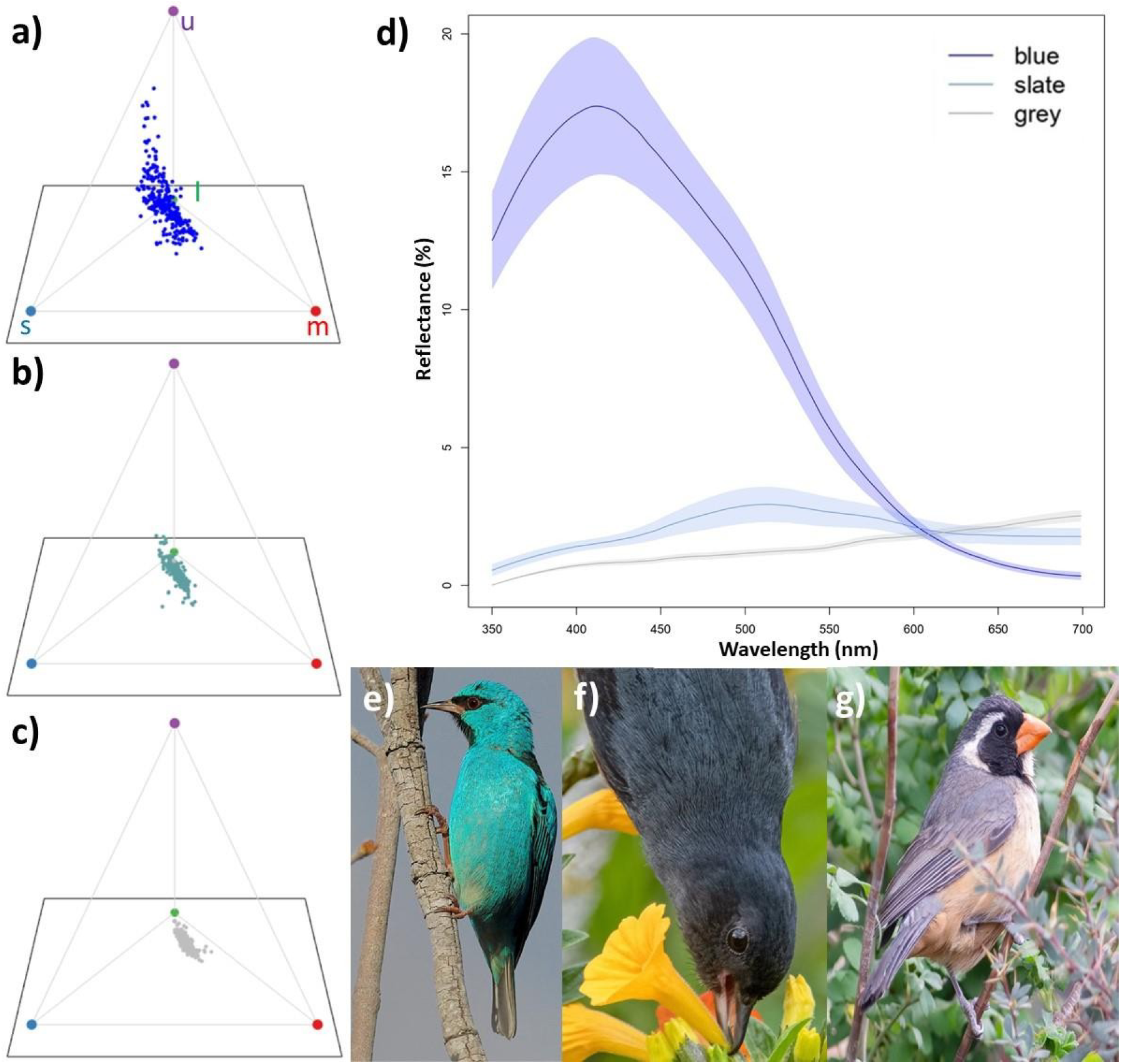
Overview of plumage colouration across three colour categories (blue, slate and grey) in tetrahedral colourspace. Panel a) is a tetrahedral colourspace with examples of blue data points, with panel e) (Blue dacnis; *Dacnis cayana*) an example of blue plumage colouration in living birds. Panel b) is tetrahedral colourspace with examples of slate data points, with panel f) (Slaty flowerpiercer; *Diglossa plumbea*) being an example of slate plumage colouration in living birds. Panel g) is a tetrahedral colourspace with examples of grey data points, with panel i) (Black-and-rufous warbling finch; *Poospiza nigrorufus*) an example of grey plumage colouration in living birds. Panel d) represents the average reflectance data of blue, slate and grey plumage patches (as indicated on the legend within the panel) with an average value shown with a line and standard error as the shaded area around the average for each colour. The data for panel d) are taken from Chapter 4 (section 4.3.2. Reflectance data with species listed in details in Supplement data: Table S1.). All photos © Daniel J. Field, University of Cambridge. Used with permission.

First, we tested for how different the blue, slate and grey colour categories are and whether the proposed intermediate colour category, i.e. slate, could be considered distinct from either blue or grey or both in avian colour space. For this purpose, we calculated the orthogonal projection of every data point onto the vector constructed in the colourspace. We constructed a vector through the colourspace that goes from one side of the tetrahedral colourspace (cone catch values: u = 0, s = 1/3, m = 1/3, l = 1/3) to its opposite side, i.e., u cone (cone catch values: s = 0, u = 1, m = 0, l = 0). The vector ̅ is defined with two sets of coordinates:

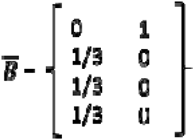

Since most of the data points have strong reflectance in the u part of the spectrum, the vector is constructed to capture the orientation of our data within the colourspace. Every data point in the colourspace, defined by the four coordinates that correspond to four cone catch values (u, s, m, l), can be represented as the vector itself:

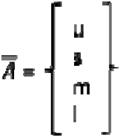

Projection is a linear transformation of any data point in the morphospace (in our case, tetrahedral colourspace), defined by four coordinates (in our case: u, s, m and l values) by orthogonal projection of that data point to a vector defined by two sets of coordinates. In the following step, an orthogonal projection of the data point ̅ to a vector̅ is calculated by:

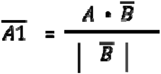

With ̅being the norm (lengths)of the ̅. The norm was calculated for every data point in our dataset, and we named them projection values and used them for further analysis. (Fig. 3.2, a; Supplement data: Table S1, projection values). After the data point in the colourspace is orthogonally projected on the predefined vector, the distance from the vector’s origin to the new point is calculated and called projection value.

**Figure 3.2.**
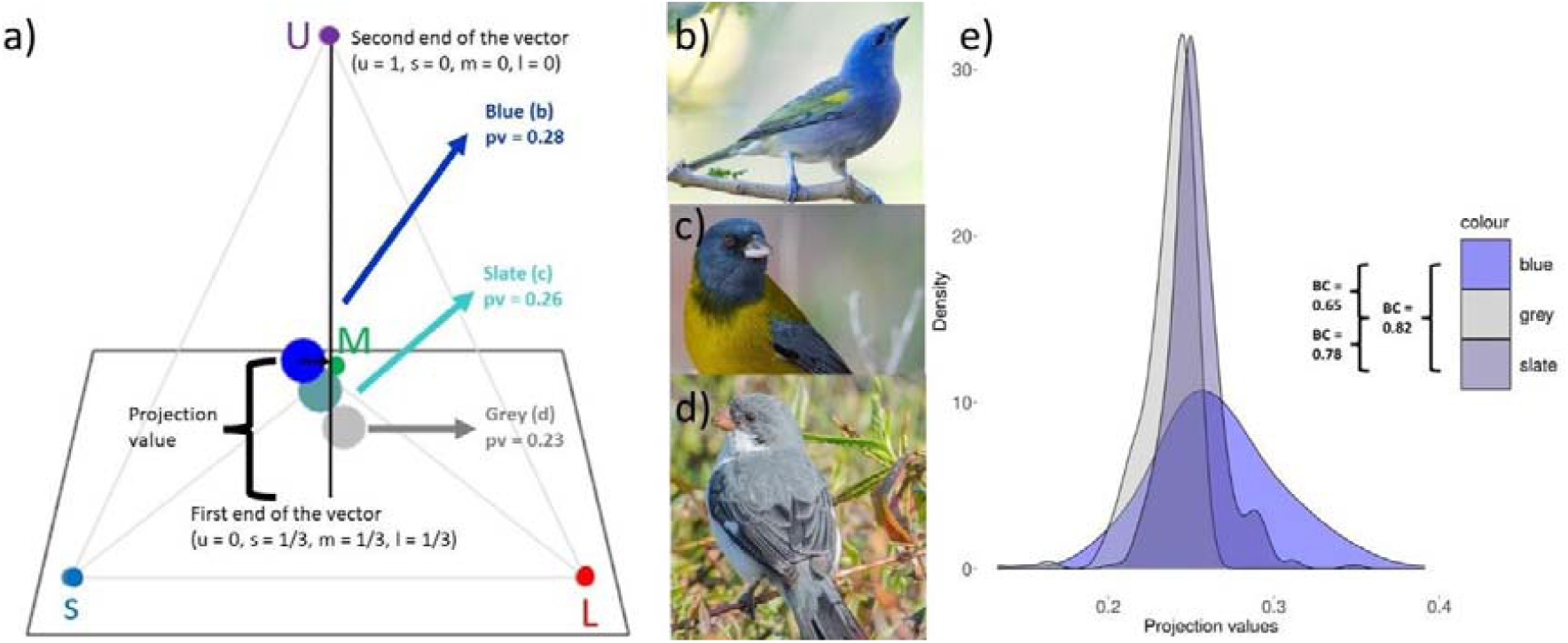
Distinctiveness analysis. Panel a) shows tetrahedral colourspace with the examples of grey, slate and blue data points. On an example of blue data point, a calculation of the projection value is shown. Blue, slate and grey data points are from the crown patch: blue is from Golden chevroned tanager (*Thraupis ornata*) (panel b), slate is from the Grey-hooded siera finch (*Phygilus gayi*) (panel c) and in grey is for White-bellied Seedeater (*Sporophila leucoptera*) (panel d). A vector is constructed throughout the colourspace that goes from one side of the tetrahedral colourspace (cone catch values: u = 0, s = 1/3, m = 1/3, l = 1/3, “first end of the vector” on the figure) to its opposite side, i.e., u cone (cone catch values: s = 0, u = 1, m = 0, l = 0; “second end of the vector” on the figure). On the example of the blue data point, we showed how is the projection value calculated, i.e. once data point is projected on the vector how far away that new data point is from the vector’s 0 end (“first end of the vector” on the firgure). Near every data point, a projection value for that data point is indicated. Panel e) shows distribution of the grey, slate and blue data points with the Battacharayya coefficient (BC) indicated for each colour par on the right-hand side. All photos © Daniel J. Field, University of Cambridge. Used with permission.

We used the projection values to calculate the Bhattacharyya coefficient which quantified the overlap between the distributions of projection values between two colour categories, i.e. blue – slate, grey – slate and blue – grey. Higher value of Bhattacharyya coefficient will indicate greater overlap between two distributions and therefore lesser distinctiveness. The calculation of both the projection values and the Battacharyya coefficient were performed using the R package dispRity (Guillerme, 2018).

### 3.4.2 Multistate analysis

To test for the evolutionary sequence from grey through slate to blue (grey -> slate -> blue) we used Reverse Jump Multistate model in BayesTraits (Currie & Meade, 2014; Pagel et al., 2004). This analysis allows testing a range of models with all possible combinations of transitions among character states (all possible pathways are illustrated in an Supplement data, Fig. S1.) The model that is best fit to our data is visited during the analysis most frequently. For our evolutionary hypothesis, this would translate to pathway that leads from any colour to grey, grey to slate, and slate to blue being more visited in comparison to any other model or have higher transition rates than the rest of parameters in our model. We used molecular phylogenies for Tanagers available from birdtree.org (Jetz et al., 2012) with 1000 random trees downloaded. From a previously collected dataset of presence and absence of blue, slate and grey colour in Tanagers, we developed a new database usedfor the RJ MCMC Multistate analysis (Supplement data: Table S2; Table S3). We coded each species in our dataset for presence/absence of each colour category such that the colour is treated as present if the bird has it on at least one body patch (crown, nape, mantle, rump, throat, breast, belly, tail, wing coverts and wing primaries/secondaries). Therefore, 0 is coded absence of the colour of interest, 1 for presence of grey colour, 2 for presence of slate colour (also if slate co-occurs with grey colour in the same bird species: two species in both males and females) and 3 for presence of blue colour (also if blue co-occurs with grey in the same species: four cases in males and nine cases in females; or slate in the same species: five species in males and four species in females). This coding scheme implicitly assumes that if a species has the ability to produce the subsequent colour in the proposed evolutionary pathway, it also can produce the preceding colour in the evolutionary pathway. A separate dataset was made for males and females (Supplement data: Table S2; Table S3) which were analysed separately due to notable differences in the plumage colouration. We repeated the same analysis for males and females, but without species that exhibit co-occurrence of two colours of our interest (Supplement data: Table S2; Table S3, in both dataset, species marked with red have double scoring and were not included in the second analysis). By excluding these species we avoid the assumption of a specific evolutionary pathway. We report the results of the latter analysis, that are qualitatively similar to the main analysis, in the Supplement data (Table S6; Table S7). For each analysis we applied Reverse Jump (RJ) MCMC which considers all possible models of evolution with proportionally visiting the best fitting one. Models were run for 220000000 iterations with Burnin 2000000 and exponential prior of value 10. Each analysis was run 3 times to confirm the consistency in our results.

## 3.5 Results

### 3.5.1 Distinctiveness analysis

First, we plotted grey, slate and blue colour categories in tetrahedral colourspace (Fig. 3.1, a, d, g). Our measure of the overlap between probability distributions (Battacharayya coefficient, BC) of the projection values for grey, blue and slate colour indicated that the categories are neither distinct nor completely overlapping (BC for slate and blue = 0.82, BC for slate and grey = 0.78, and BC for grey and blue = 0.65) (Fig. 3.2, e). A Battacharayya coefficient greater than 0.95 would indicate overlap and less than 0.05 would indicate distinctiveness (Guillerme & Cooper, 2016). These results can be interpreted as evidence that grey and blue are opposite ends of continuum in which slate is an intermediate state.

### 3.5.1 Multistate

Our RJ MCMC Multistate analyses revealed that a pathway from any other colour to grey through slate leading to blue is present. While other evolutionary transitions among the coded characters are possible in both males and females, no other pathways led towards blue colour.

In females, the 95% credible set consists of only one model (Fig. 3.3, b). The model suggests that transition from any other colour to slate (q02), any other colour to blue (q03), grey to blue (q13), blue to grey (q31) and blue to slate (q32) are very unlikely to happen. This is indicated by low percentage of models that estimate these transition rates to not be zero (0.01% -3.32%) (Fig. 3.3, d). On the other hand, transition from any other colour to grey and vice versa (q01, q10), grey to slate and vice versa (q12 and q21), slate to any other colour (q20), slate to blue (q23) and blue to any other colour (q30) are likely to happen. These parameters are estimated as non-zero in a very high percentage of all models (99.4% -100%) (Fig. 3.3, d). All the transitions that do happen have the same mean values of 0.06 indicating that when the transitions do happen, their average rate is equal(Fig. 3.3, c). Full results are reported in the Supplement data: Tale S5. Results without double scoring species are reported in the Supplement data: Table S7.

**Figure 3.3.**
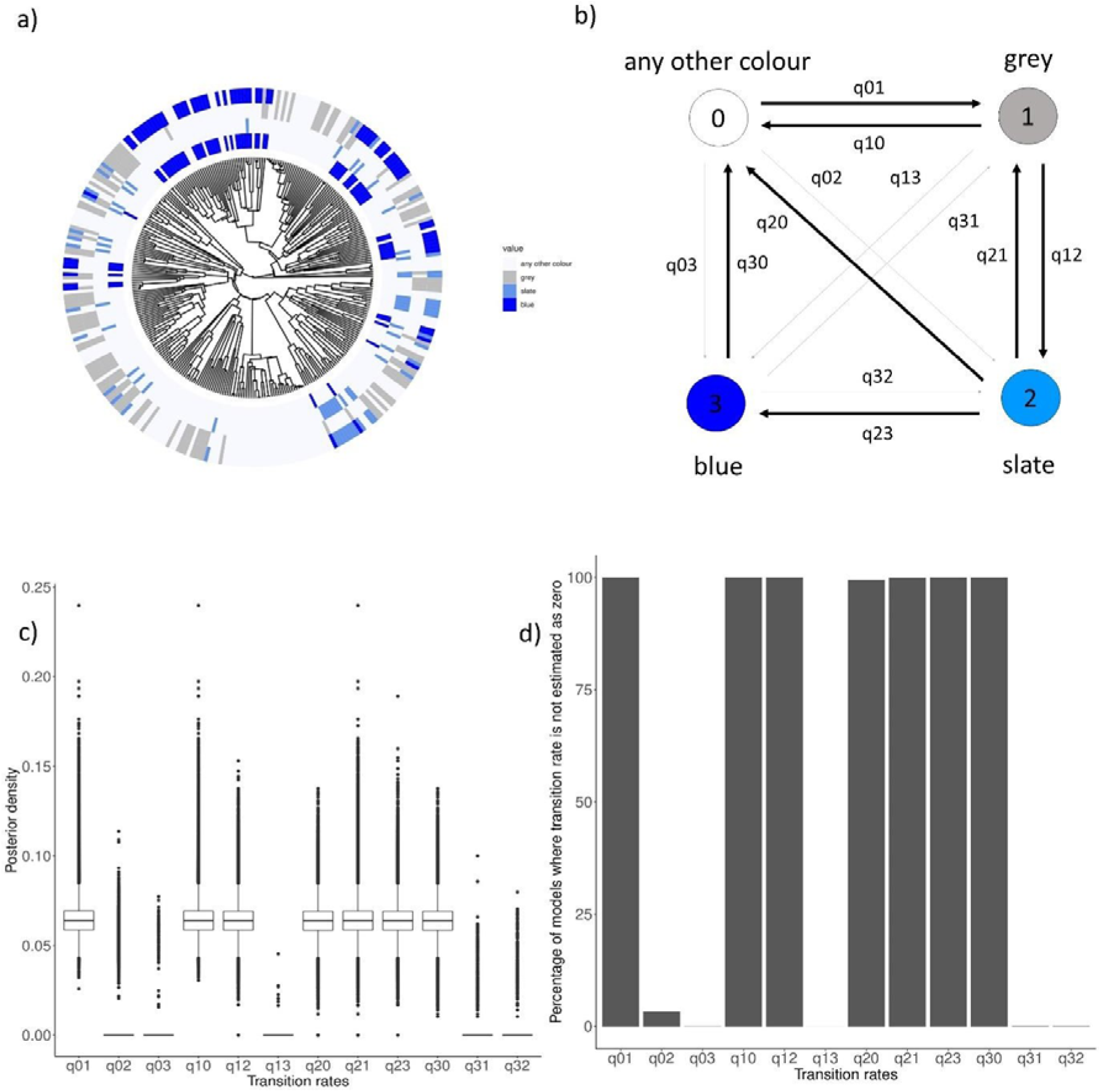
The evolution of blue, slate and grey colour in the plumage of female Tanagers. Panel a shows the phylogeny of 319 Tanagers species with the plotting of blue, slate and grey on tips of the phylogeny. The first circle indicates the presence of blue colour, the second circle indicates the presence of slate colour, the third circle indicates the presence of grey colour and the fourth circle represents the coding of colours used for the analysis. The presence of colour in the plumage of species is indicated with the presence of the bar, while the lack of it is represented with the lack of it for each circle. Panel b shows schematics of the evolutionary pathway between coded colour states. Transition rates that are present are represented with thicker lines, while those that are not detected are represented with thinner lines. Panel c shows posterior densities of the transition rates from the RJ Multistate model. Panel d shows the percentage of models from 220 000 000 iterations where transition rates are not estimated as zero. A high percentage indicates that the transition is happening (q01, q10, q12, q20, q21, q23, q30). A low percentage indicates that the transitions are not happening (q02, q03, q13, q31, q32). Transition rates on panels c and d indicate the following transition rates: q01 is any other colour to grey, q02 is any other colour to slate, q03 is any other colour to blue, q10 is grey to any other colour, q12 is grey to slate, q13 is grey to blue, q20 is slate to any other colour, q21 is slate to grey, q23 is slate to blue, q30 is blue to any other colour, q31 is blue to grey, q32 is blue to slate.

In males, the 95% credible set of all models visited during the analysis consists of three possible models. Below we focus on the most frequently visited model (144,398 times out of 218,000) (Fig. 3.4, b). In this model, transitions from any other colour to blue (q03), grey to blue (q13), blue to grey (q31) and blue to slate (q32) are very unlikely to happen. This is indicated by low percentage of models that include these transitions at a non-zero rate (0.1% -1.9%) (Fig. 3.4, d). Contrary to that, transitions from any other colour to grey (and reverse) (q01, q10), grey to slate (and reverse) (q12, q21), slate to blue (q23) and blue to any other colour (q30), are frequently estimated to be non-zero (99.999 – 100%) indicating that they are likely to happen (Fig. 3.4, d). In between these two extremes, both the gain of slate from any other colour and vice versa (q02, q20) occur in an intermediate percentage of models at a non-zero rate (93.45% and 70.45% respectfully) (Fig. 3.4, d). This would indicate that these transitions occur, but are not as likely as those with a very low percentage of models that estimate them to be zero. The most favoured model sets all non-zero transition rates to be equal with a rate of 0.4 transitions per lineage per million years (Fig. 3.4, c). The other two models in the 96% credible set are visited less frequently (57,610 and 8,357 times respectively). They differ from the most visited model only in two parameters: in the second most visited model the parameter q20 = 0 whereas in the third most visited model q02 = 0. These parameters relate to transitions between any other colour and slate. The presence or absence of these specific transitions does not qualitatively alter our main conclusions on the most likely pathways in grey-slate-blue colourspace. Full results are reported in the Supplement data: Table S4. Results without double scoring species are reported in the Supplement data: Table S6.

**Figure 3.4.**
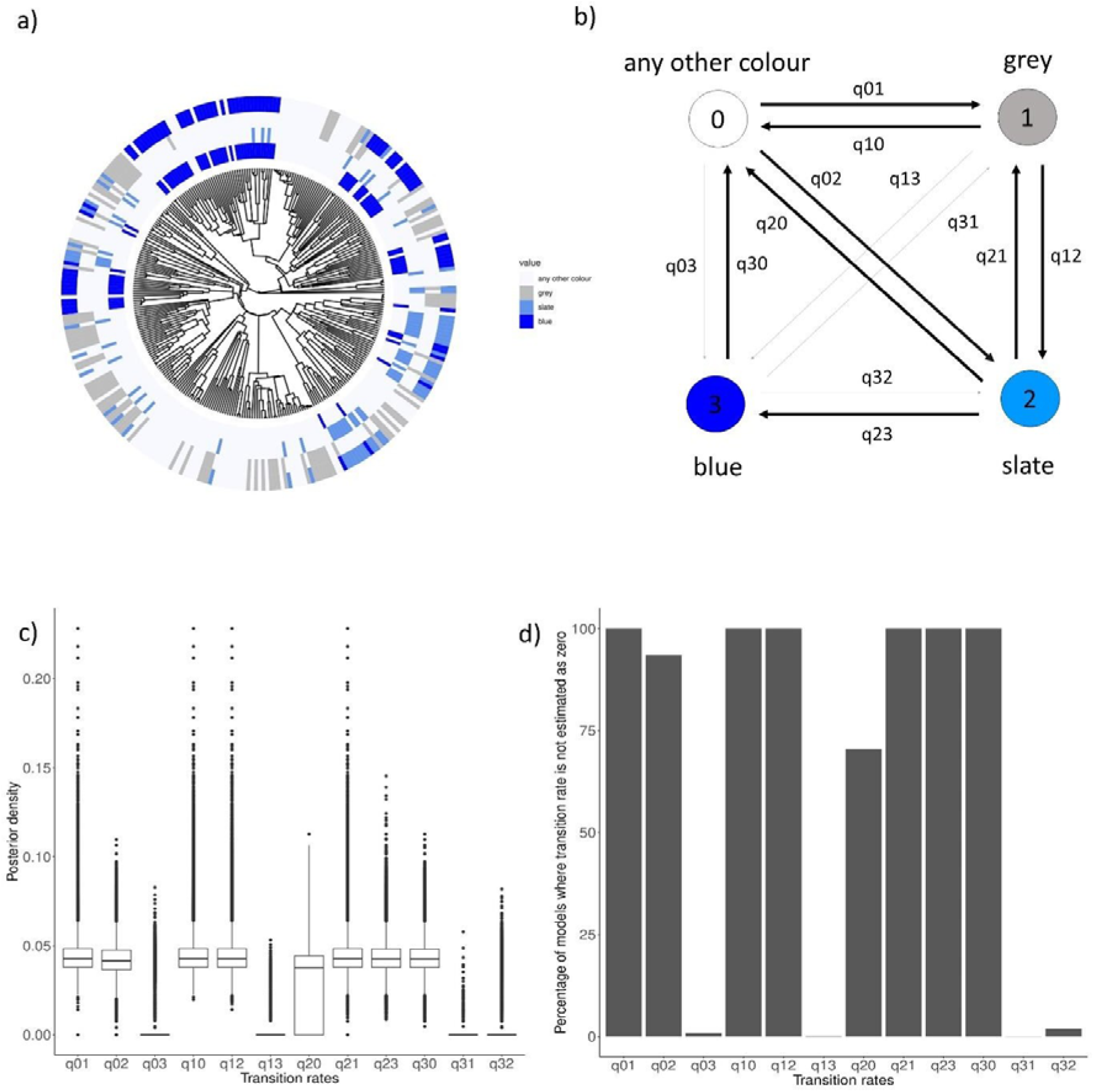
The evolution of blue, slate and grey colour in the plumage of male Tanagers. Panel a shows the phylogeny of 319 Tanagers species with the plotting of blue, slate and grey on tips of the phylogeny. The first circle represent the blue colour, the second circle represents slate colour, the third circle represents the grey colour and the fourth circle represents the coding of colours used for the analysis. The presence of colour in the plumage of species is indicated with the presence of the bar, while the lack of it is represented with the lack of it for each circle. Panel b shows schematics of the evolutionary pathway between coded colour states. Transition rates that are present are represented with ticker lines, while those that are not detected are represented with tinner lines. Panel c shows posterior densities of the transition rates from the RJ Multistate model. Panel d shows the percentage of models from 220 000 000 iterations where transition rates are not estimated as zero. A high percentage indicates that the transition is happening (q01, q10, q12, q20, q21, q23, q30). A low percentage indicates that the transitions are not happening (q02, q03, q13, q31, q32). Transition rates on panels c and d indicate the following transition rates: q01 is any other colour to grey, q02 is any other colour to slate, q03 is any other colour to blue, q10 is grey to any other colour, q12 is grey to slate, q13 is grey to blue, q20 is slate to any other colour, q21 is slate to grey, q23 is slate to blue, q30 is blue to any other colour, q31 is blue to grey, q32 is blue to slate.

Multistate analysis also allows us to estimate ancestral state in the basal node of the entire clade for the categorical variables that we use for the analysis (i.e. any other colour, grey, slate, and blue). The model output provides a probability estimate of each state in the basal node of the Tanagers. In males, the highest probability of the ancestral estimate for the basal node is for the slate colour (P(slate) = 0.587), the lower and equivalent probability is for the grey colour, and any other category colour (P(grey) = 0.175, P(any other colour)=0.188) and the lowest probability is for the blue colour (P(blue) = 0.05). In females, the highest probability of the ancestral estimate for the basal node is slate colour (P(slate) = 0.49), followed by grey colour (P(grey) = 0.37), while blue and any other colour category has the lowest and similar probabilities (P(blue) = 0.07 and P(any other colour) = 0.06). The full results are reported in Additional table 5.; (https://figshare.com/s/1110fce894e65a69c329).

## 3.6 Discussion

Our results showed that slate colour is an intermediate colour with blue colour on one and grey on the other side of the area occupied by these three colours in the tetrahedral colourspace of Tanager plumage colouration. Furthermore, analysis of all possible evolutionary pathways between grey, slate and blue indicated that the most likely pathway for the evolution of the blue colour is the route from grey colour and through the slate colour. Blue colour can equally likely evolve to other parts of the colourspace and we propose white and black colour as the most likely candidates of this transition. Interestingly, slate colour could have an independent evolutionary origin in males suggesting that discovered evolutionary pathway towards blue colour might be rare and hard to achieve.

Due to the fourth cone in their retina, birds’ colour vision is extended into the UV part of the light spectrum, making analysis of plumage colouration within the tetrahedral colourspace a crucial partof the framework for understanding colour evolution (Endler & Mielke Jr, 2005; Stoddard & Prum, 2008). We confirm that, as expected, slate is part of a continuum in tetrahedral colourspace between grey and blue. While grey and blue are distinct from one another, the slate colour category overlaps with both grey and blue colour equally (Fig. 3.2, e) suggesting that slate colour shares properties both of grey colour and blue colour. To what extent the observed overlap in tetrahedral colourspace translates to overlap in the nanostructures of these colours is yet to be seen, but some general assumptions based on what we know about these nanostructures can be made. Slate colour was previously described to be weakly nanostructured, implying that the feather barbs have at least a partially developed medullary spongy layer that would produce what seems to be blue wavelengths in this colour (Saranathan et al., 2012). Since thickness of the medullary layer is important for hue variance in non-iridescent structural colours (Fan et al., 2019), it might be that observed the hue of slate is due to the lack of the sufficient thickness of rudimentary medullary spongy layer. Alternatively, we suggest that melanin deposition masks the blue colour produced by the medullary layer nanostructure as this effect has previously been observed in the case of blue colour (Doucet et al., 2004). In the case of slate, melanosomes characteristic of grey colour could be involved in this process (Babarović et al., 2019).

We tested the macroevolutionary origins and dynamics of non-iridescent structural colours in Tanagers. Our results revealed that the most common pathway for the evolution of blue colour in Tanagers is from slate colour (Fig. 3.3, b; Fig. 3.4, b). Any other evolutionary pathway to non-iridescent structural blue is highly unlikely (Fig. 3.3, b; Fig. 3.4, b). Evolution from pigmentary to structural colour has been recorded previously and explained as a process of evolutionary tinkering (Shawkey et al., 2006). This process would involve the evolution of a new phenotype by rearrangements of elements of an already existing phenotype (Bockaert & Pin, 1999, 1999). In the case of the evolution from pigmentary grey to non-iridescent structural colour blue this would involve a two-stage process: 1) arrangement of melanosomes in a layer next to the central shaft, and 2) development of a keratin and air nanostructure in medullary cells of the feather barbs responsible to produce the colour. Due to the lack of information of the internal anatomy of slate feather barbs, the precise sequence and extent of these processes remains to be tested. The existence of the intermediary stage is vital for the emergence of the blue colouration since direct pathways from “any other colour” to blue have not been detected in our analysis (Fig. 3.3, b; Fig. 3.4, b). Nevertheless, the requirement for an intermediary stage indicates that the process of evolution of blue colour via this route is likely hard to achieve and rare.

Interestingly, slate colour is the most likely ancestral state estimate for the plumage colour out of the colours we have investigated. This could further indicate the dynamics of the evolutionary processes we observe in this analysis. Namely, while slate is the most likely ancestral state, and we know that transitions between slate and grey are bi-directional and common, we suggest that these transitions happened early in the evolutionary history of this clade. This also suggests that the evolutionary sequence we observe is not hierarchical for grey and slate colours, while blue can evolve only from slate colour. The observed outcome could depend on the ecological and behavioural characteristics of the species having these colours in the modern clade and the hypothesized ancestral species to modern Tanagers.

The transition from blue to “any other colour” has also been detected within our plumage colour evolution model (Fig. 3.3, b; Fig. 3.4, b). Transitions from blue colour to other colours have been detected previously in birds’ plumage. For example, in swallow tanager (*Tarsina viridis*) white belly feathers have a slight blue-green wash on the tips of the distant barbs (Bazzano et al., 2021). Production of that colour was assigned to the keratin and air matrix which was like that of blue feathers. The rest of the plumage of swallow tanager is green-blue produced by the keratin and air nanostructure in medullary cells of feather barbs with an underlying melanosome layer. This common nanostructural component might indicate a potential evolutionary transition between white colour and non-iridescent structural colours. Furthermore, the black plumage of two island subspecies of the White-winged fairywren *Malurus leucopterus* (*M. l. leucopterus* and *M. l. edouardi*) has been confirmed by genetic and nanostructural analysis to have evolved from blue plumage colouration of mainland subspecies *M. l. leuconotus* (Doucet et al., 2004). The black plumage of the two island subspecies has a rudimentary spongy layer embedded with melanosomes that cloud blue colour production resulting in black. These examples are consistent with our finding that blue can evolve into multiple other colours.

The existence of the intermediary stage could also indicate that slate colour could have a separate ecological or signalling purpose within a bird’s plumage colouration which made it a stable phenotype. Due to the lack of information on the nanostructural basis of slate colour, it is impossible to predict from which colour this transition might happen. Both blue and grey colour have predictions of their signalling capacities in their adjacent light environments. While blue colour achieves increased conspicuousness in woodland light environment, grey colour does the same in the open light environment. It would be interesting to test experimentally the signalling capacity of slate colour in the different light environments and whether slate colour contains trade-off between signalling properties of blue and grey colour.

