## Supplement Tables S1-S7 and Supplement Figure S1 for "Evolutionary dynamics of pigmentary grey and non-iridescent structural blue colouration in Tanagers (family: Thraupidae)": Evolutionary dynamics of pigmentary grey and non- iridescent structural blue colouration in Tanagers (family- Thraupidae) _ supplement.docx

**Table S1.** Table containing values of colour measurements and colour descriptions with sex, latin and english name and region of the plumage patch where the measurement was taken. Column u, s, m and l are the results of the quanitification of the colour from digitally calibrated images. Column colour_HBW is the plumage colour description from the Birds of the World and colour coding column is the result of classification of the colour descriptions from HBW into the colour categories,

i.e. blue, slate and grey. Finally, projection values is the column containing calculations of the projection values.

| Latin name | English Name | sex | regi  on | colour HBW | colour  coding | u | s | m | l | projectio  n values |
| --- | --- | --- | --- | --- | --- | --- | --- | --- | --- | --- |
| Anisognathus  lacrymosus | Lacrimose  Mountain-Tanager | M | cro | slaty blue | blue | 0.20  8 | 0.27  1 | 0.270 | 0.251 | 0.271 |
| Anisognathus  melanogenys | Black-cheeked  Mountain-Tanager | M | cro | shining blue | blue | 0.20  6 | 0.28  4 | 0.266 | 0.244 | 0.284 |
| Buthraupis  eximia | Black-chested  Mountain-Tanager | M | cro | blue | blue | 0.27  0 | 0.30  3 | 0.235 | 0.192 | 0.303 |
| Conirostrum  bicolor | Bicolored Conebill | M | cro | pale bluish-grey | blue | 0.18  4 | 0.25  1 | 0.288 | 0.277 | 0.251 |
| Cyanerpes  caeruleus | Purple  Honeycreeper | M | cro | lustrous violet-  blue | blue | 0.54  0 | 0.21  8 | 0.132 | 0.110 | 0.218 |
| Cyanerpes  cyaneus | Red-legged  Honeycreeper | M | cro | azure / blue | blue | 0.14  7 | 0.21  8 | 0.272 | 0.363 | 0.218 |
| Cyanerpes  lucidus | Shining  Honeycreeper | M | cro | ultramarine blue | blue | 0.41  1 | 0.30  0 | 0.169 | 0.120 | 0.300 |
| Cyanerpes  nitidus | Short-billed  Honeycreeper | M | cro | purplish-blue | blue | 0.35  0 | 0.37  3 | 0.163 | 0.114 | 0.373 |
| Cyanicterus  cyanicterus | Blue-backed  Tanager | M | cro | bright cadet-blue | blue | 0.40  5 | 0.30  8 | 0.168 | 0.120 | 0.308 |
| Dacnis cayana | Blue Dacnis | M | cro | bright turquoise-  blue | blue | 0.26  2 | 0.29  5 | 0.304 | 0.139 | 0.295 |
| Dacnis hartlaubi | Turquoise Dacnis | M | cro | turquoise | blue | 0.28  0 | 0.28  7 | 0.293 | 0.140 | 0.287 |
| Dacnis lineata | Black-faced Dacnis | M | cro | turquoise-blue | blue | 0.28  0 | 0.29  9 | 0.280 | 0.141 | 0.299 |
| Dacnis nigripes | Black-legged | M | cro | greenish blue / | blue | 0.23 | 0.28 | 0.318 | 0.164 | 0.284 |

|  | Dacnis |  |  | turquoise |  | 4 | 4 |  |  |  |
| --- | --- | --- | --- | --- | --- | --- | --- | --- | --- | --- |
| Dacnis venusta | Scarlet-thighed  Dacnis | M | cro | black / turquoise | blue | 0.22  7 | 0.30  2 | 0.323 | 0.148 | 0.302 |
| Dacnis viguieri | Viridian Dacnis | M | cro | green | blue | 0.29  7 | 0.16  6 | 0.379 | 0.159 | 0.166 |
| Diglossa  albilatera | White-sided  Flowerpiercer | M | cro | dark slate-grey | blue | 0.21  2 | 0.25  2 | 0.279 | 0.258 | 0.252 |
| Diglossa cyanea | Masked  Flowerpiercer | M | cro | dark blue | blue | 0.27  8 | 0.31  5 | 0.232 | 0.176 | 0.315 |
| Diglossa glauca | Deep-blue  Flowerpiercer | M | cro | deep blue | blue | 0.35  5 | 0.25  9 | 0.209 | 0.177 | 0.259 |
| Diglossa  indigotica | Indigo  Flowerpiercer | M | cro | indigo-blue | blue | 0.30  5 | 0.35  5 | 0.199 | 0.141 | 0.355 |
| Iridosornis analis | Yellow-throated  Tanager | M | cro | deep purplish blue | blue | 0.20  0 | 0.25  9 | 0.281 | 0.259 | 0.259 |
| Iridosornis porphyrocephalu  s | Purplish-mantled Tanager | M | cro | dark blue | blue | 0.34  0 | 0.24  1 | 0.221 | 0.198 | 0.241 |
| Pipraeidea  melanonota | Fawn-breasted  Tanager | M | cro | medium blue | blue | 0.29  8 | 0.34  9 | 0.205 | 0.147 | 0.349 |
| Porphyrospiza  caerulescens | Blue Finch | M | cro | bright indigo-blue | blue | 0.26  7 | 0.28  7 | 0.243 | 0.203 | 0.287 |
| Tangara  cyanicollis | Blue-necked  Tanager | M | cro | blue | blue | 0.22  1 | 0.30  3 | 0.325 | 0.152 | 0.303 |
| Tangara  cyanocephala | Red-necked  Tanager | M | cro | blue-violet | blue | 0.34  3 | 0.36  8 | 0.170 | 0.119 | 0.368 |
| Tangara  nigrocincta | Masked Tanager | M | cro | pale lavender | blue | 0.23  8 | 0.31  6 | 0.249 | 0.197 | 0.316 |
| Tangara seledon | Green-headed  Tanager | M | cro | turquoise-green | blue | 0.07  8 | 0.27  5 | 0.446 | 0.201 | 0.275 |
| Tangara vassorii | Blue-and-black  Tanager | M | cro | dark cobalt blue | blue | 0.39  9 | 0.30  9 | 0.169 | 0.122 | 0.309 |
| Tersina viridis | Swallow Tanager | M | cro | bright turquoise-  blue | blue | 0.22  5 | 0.25  5 | 0.344 | 0.176 | 0.255 |
| Thraupis abbas | Yellow-winged  Tanager | M | cro | campanula-blue | blue | 0.42  6 | 0.27  3 | 0.164 | 0.137 | 0.273 |
| Thraupis  bonariensis | Blue-and-yellow  Tanager | M | cro | dull blue | blue | 0.30  3 | 0.28  3 | 0.231 | 0.183 | 0.283 |
| Thraupis  cyanocephala | Blue-capped  Tanager | M | cro | shining  cornflower-blue | blue | 0.42  9 | 0.27  3 | 0.169 | 0.129 | 0.273 |
| Thraupis ornata | Golden-chevroned  Tanager | M | cro | shining cadet-blue | blue | 0.32  4 | 0.28  3 | 0.209 | 0.183 | 0.283 |
| Xenodacnis  parina | Tit-like Dacnis | M | cro | dark blue | blue | 0.33  7 | 0.30  1 | 0.202 | 0.160 | 0.301 |
| Anisognathus  melanogenys | Black-cheeked  Mountain-Tanager | M | nap | shining blue | blue | 0.29  3 | 0.26  9 | 0.231 | 0.207 | 0.269 |
| Buthraupis  eximia | Black-chested  Mountain-Tanager | M | nap | blue | blue | 0.46  2 | 0.25  3 | 0.162 | 0.124 | 0.253 |
| Buthraupis  montana | Hooded  Mountain-Tanager | M | nap | shining pale milky  blue | blue | 0.21  1 | 0.24  9 | 0.286 | 0.254 | 0.249 |
| Chlorophanes  spiza | Green  Honeycreeper | M | nap | glossy viridian  green | blue | 0.05  1 | 0.26  7 | 0.460 | 0.221 | 0.267 |
| Conirostrum  bicolor | Bicolored Conebill | M | nap | pale bluish-grey | blue | 0.20  9 | 0.26  0 | 0.279 | 0.252 | 0.260 |
| Cyanerpes  caeruleus | Purple  Honeycreeper | M | nap | lustrous violet-  blue | blue | 0.64  0 | 0.19  6 | 0.154 | 0.011 | 0.196 |
| Cyanerpes  lucidus | Shining  Honeycreeper | M | nap | ultramarine blue | blue | 0.41  5 | 0.28  4 | 0.178 | 0.122 | 0.284 |
| Cyanerpes  nitidus | Short-billed  Honeycreeper | M | nap | purplish-blue | blue | 0.34  5 | 0.34  8 | 0.174 | 0.134 | 0.348 |
| Cyanicterus  cyanicterus | Blue-backed  Tanager | M | nap | bright cadet-blue | blue | 0.36  6 | 0.29  6 | 0.192 | 0.146 | 0.296 |
| Dacnis cayana | Blue Dacnis | M | nap | bright turquoise-  blue | blue | 0.27  8 | 0.30  5 | 0.291 | 0.127 | 0.305 |
| Dacnis nigripes | Black-legged  Dacnis | M | nap | greenish blue /  turquoise | blue | 0.24  8 | 0.28  0 | 0.304 | 0.168 | 0.280 |
| Dacnis venusta | Scarlet-thighed  Dacnis | M | nap | turquoise | blue | 0.23  7 | 0.31  1 | 0.300 | 0.152 | 0.311 |

| Dacnis viguieri | Viridian Dacnis | M | nap | greenish blue | blue | 0.27  5 | 0.19  6 | 0.378 | 0.151 | 0.196 |
| --- | --- | --- | --- | --- | --- | --- | --- | --- | --- | --- |
| Diglossa  albilatera | White-sided  Flowerpiercer | M | nap | dark slate-grey | blue | 0.22  9 | 0.24  5 | 0.268 | 0.258 | 0.245 |
| Diglossa cyanea | Masked  Flowerpiercer | M | nap | dark blue | blue | 0.30  6 | 0.31  7 | 0.218 | 0.159 | 0.317 |
| Diglossa glauca | Deep-blue  Flowerpiercer | M | nap | deep blue | blue | 0.36  2 | 0.25  1 | 0.208 | 0.180 | 0.251 |
| Diglossa  indigotica | Indigo  Flowerpiercer | M | nap | indigo-blue | blue | 0.30  1 | 0.33  0 | 0.206 | 0.164 | 0.330 |
| Iridosornis analis | Yellow-throated  Tanager | M | nap | greenish turquoise | blue | 0.21  9 | 0.26  8 | 0.267 | 0.247 | 0.268 |
| Iridosornis porphyrocephalu  s | Purplish-mantled Tanager | M | nap | dark blue | blue | 0.37  9 | 0.27  0 | 0.194 | 0.157 | 0.270 |
| Pipraeidea  melanonota | Fawn-breasted  Tanager | M | nap | medium blue | blue | 0.35  7 | 0.34  5 | 0.166 | 0.131 | 0.345 |
| Porphyrospiza  caerulescens | Blue Finch | M | nap | bright indigo-blue | blue | 0.24  0 | 0.26  7 | 0.257 | 0.236 | 0.267 |
| Tangara  cyanicollis | Blue-necked  Tanager | M | nap | blue | blue | 0.22  7 | 0.31  5 | 0.304 | 0.155 | 0.315 |
| Tangara  cyanocephala | Red-necked  Tanager | M | nap | blue-violet | blue | 0.21  9 | 0.22  5 | 0.215 | 0.341 | 0.225 |
| Tangara  nigrocincta | Masked Tanager | M | nap | pale lavender | blue | 0.26  1 | 0.31  7 | 0.235 | 0.187 | 0.317 |
| Tangara seledon | Green-headed  Tanager | M | nap | turquoise-green | blue | 0.09  4 | 0.27  3 | 0.435 | 0.198 | 0.273 |
| Tangara vassorii | Blue-and-black  Tanager | M | nap | dark cobalt blue | blue | 0.41  4 | 0.31  8 | 0.152 | 0.116 | 0.318 |
| Tersina viridis | Swallow Tanager | M | nap | bright turquoise-  blue | blue | 0.23  2 | 0.26  2 | 0.349 | 0.156 | 0.262 |
| Thraupis abbas | Yellow-winged  Tanager | M | nap | campanula-blue | blue | 0.39  2 | 0.24  2 | 0.194 | 0.172 | 0.242 |
| Thraupis  bonariensis | Blue-and-yellow  Tanager | M | nap | dull blue | blue | 0.31  9 | 0.28  5 | 0.222 | 0.174 | 0.285 |
| Thraupis  cyanocephala | Blue-capped  Tanager | M | nap | shining  cornflower-blue | blue | 0.45  0 | 0.25  6 | 0.164 | 0.129 | 0.256 |
| Thraupis ornata | Golden-chevroned  Tanager | M | nap | shining cadet-blue | blue | 0.38  3 | 0.27  9 | 0.181 | 0.156 | 0.279 |
| Xenodacnis  parina | Tit-like Dacnis | M | nap | dark blue | blue | 0.31  7 | 0.31  2 | 0.213 | 0.158 | 0.312 |
| Anisognathus  lacrymosus | Lacrimose  Mountain-Tanager | M | man | slaty blue | blue | 0.21  5 | 0.25  8 | 0.268 | 0.258 | 0.258 |
| Anisognathus  melanogenys | Black-cheeked  Mountain-Tanager | M | man | dark grey-blue | blue | 0.21  8 | 0.26  7 | 0.273 | 0.242 | 0.267 |
| Buthraupis  montana | Hooded  Mountain-Tanager | M | man | shining dark blue | blue | 0.60  2 | 0.23  3 | 0.131 | 0.035 | 0.233 |
| Chlorophanes  spiza | Green  Honeycreeper | M | man | glossy viridian  green | blue | 0.05  1 | 0.32  2 | 0.431 | 0.197 | 0.322 |
| Conirostrum  bicolor | Bicolored Conebill | M | man | pale bluish-grey | blue | 0.21  1 | 0.25  8 | 0.277 | 0.254 | 0.258 |
| Cyanerpes  caeruleus | Purple  Honeycreeper | M | man | lustrous violet-  blue | blue | 0.57  5 | 0.25  1 | 0.133 | 0.042 | 0.251 |
| Cyanerpes  lucidus | Shining  Honeycreeper | M | man | ultramarine blue | blue | 0.40  4 | 0.26  6 | 0.186 | 0.144 | 0.266 |
| Cyanerpes  nitidus | Short-billed  Honeycreeper | M | man | purplish-blue | blue | 0.37  7 | 0.34  7 | 0.158 | 0.118 | 0.347 |
| Cyanicterus  cyanicterus | Blue-backed  Tanager | M | man | bright cadet-blue | blue | 0.31  4 | 0.26  6 | 0.240 | 0.180 | 0.266 |
| Diglossa  albilatera | White-sided  Flowerpiercer | M | man | dark slate-grey | blue | 0.23  5 | 0.24  7 | 0.265 | 0.253 | 0.247 |
| Diglossa cyanea | Masked  Flowerpiercer | M | man | dark blue | blue | 0.29  5 | 0.30  5 | 0.227 | 0.172 | 0.305 |
| Diglossa glauca | Deep-blue  Flowerpiercer | M | man | deep blue | blue | 0.40  2 | 0.26  4 | 0.188 | 0.145 | 0.264 |
| Diglossa  indigotica | Indigo  Flowerpiercer | M | man | indigo-blue | blue | 0.31  2 | 0.32  5 | 0.203 | 0.161 | 0.325 |
| Dubusia taeniata | Buff-breasted  Mountain-Tanager | M | man | black / dark blue | blue | 0.23  3 | 0.26  5 | 0.261 | 0.240 | 0.265 |

| Iridosornis analis | Yellow-throated  Tanager | M | man | greenish turquoise | blue | 0.15  8 | 0.27  0 | 0.310 | 0.262 | 0.270 |
| --- | --- | --- | --- | --- | --- | --- | --- | --- | --- | --- |
| Iridosornis porphyrocephalu  s | Purplish-mantled Tanager | M | man | dark blue | blue | 0.26  6 | 0.30  6 | 0.242 | 0.186 | 0.306 |
| Iridosornis  rufivertex | Golden-crowned  Tanager | M | man | purplish-blue | blue | 0.24  4 | 0.25  7 | 0.254 | 0.245 | 0.257 |
| Pipraeidea  melanonota | Fawn-breasted  Tanager | M | man | blackish blue | blue | 0.37  9 | 0.29  0 | 0.179 | 0.152 | 0.290 |
| Porphyrospiza  caerulescens | Blue Finch | M | man | bright indigo-blue | blue | 0.25  6 | 0.27  9 | 0.250 | 0.215 | 0.279 |
| Stephanophorus  diadematus | Diademed Tanager | M | man | shining dark blue | blue | 0.48  1 | 0.24  5 | 0.155 | 0.119 | 0.245 |
| Tangara vassorii | Blue-and-black  Tanager | M | man | dark cobalt blue | blue | 0.42  3 | 0.32  1 | 0.148 | 0.108 | 0.321 |
| Tangara  viridicollis | Silvery Tanager | M | man | pale turquoise | blue | 0.04  3 | 0.28  4 | 0.365 | 0.307 | 0.284 |
| Tangara  xanthocephala | Saffron-crowned  Tanager | M | man | black / turquoise  green | blue | 0.08  0 | 0.24  7 | 0.414 | 0.260 | 0.247 |
| Tersina viridis | Swallow Tanager | M | man | bright turquoise-  blue | blue | 0.24  1 | 0.26  4 | 0.337 | 0.159 | 0.264 |
| Thraupis  glaucocolpa | Glaucous Tanager | M | man | smoky grey /  greenish | blue | 0.18  1 | 0.24  2 | 0.305 | 0.272 | 0.242 |
| Xenodacnis  parina | Tit-like Dacnis | M | man | dark blue | blue | 0.29  7 | 0.32  4 | 0.221 | 0.158 | 0.324 |
| Anisognathus  igniventris | Scarlet-bellied  Mountain-Tanager | M | rum | blue | blue | 0.28  5 | 0.29  2 | 0.229 | 0.194 | 0.292 |
| Anisognathus  lacrymosus | Lacrimose  Mountain-Tanager | M | rum | slaty blue | blue | 0.25  3 | 0.24  7 | 0.249 | 0.251 | 0.247 |
| Anisognathus  melanogenys | Black-cheeked  Mountain-Tanager | M | rum | dark grey-blue | blue | 0.21  9 | 0.24  7 | 0.277 | 0.257 | 0.247 |
| Buthraupis  montana | Hooded  Mountain-Tanager | M | rum | shining dark blue | blue | 0.29  3 | 0.23  6 | 0.240 | 0.232 | 0.236 |
| Chlorophanes  spiza | Green  Honeycreeper | M | rum | glossy viridian  green | blue | 0.06  9 | 0.33  0 | 0.396 | 0.204 | 0.330 |
| Conirostrum  bicolor | Bicolored Conebill | M | rum | pale bluish-grey | blue | 0.21  1 | 0.24  8 | 0.274 | 0.266 | 0.248 |
| Cyanerpes  caeruleus | Purple  Honeycreeper | M | rum | lustrous violet-  blue | blue | 0.60  2 | 0.22  4 | 0.135 | 0.039 | 0.224 |
| Cyanerpes  lucidus | Shining  Honeycreeper | M | rum | ultramarine blue | blue | 0.47  8 | 0.24  0 | 0.186 | 0.095 | 0.240 |
| Cyanerpes  nitidus | Short-billed  Honeycreeper | M | rum | purplish-blue | blue | 0.40  1 | 0.33  7 | 0.135 | 0.127 | 0.337 |
| Cyanicterus  cyanicterus | Blue-backed  Tanager | M | rum | bright cadet-blue | blue | 0.27  6 | 0.26  0 | 0.253 | 0.211 | 0.260 |
| Dacnis cayana | Blue Dacnis | M | rum | turquoise blue | blue | 0.21  2 | 0.27  2 | 0.257 | 0.259 | 0.272 |
| Dacnis hartlaubi | Turquoise Dacnis | M | rum | turquoise | blue | 0.23  7 | 0.29  1 | 0.254 | 0.218 | 0.291 |
| Dacnis nigripes | Black-legged Dacnis | M | rum | light turquoise / greenish blue /  black | blue | 0.23  5 | 0.27  9 | 0.281 | 0.205 | 0.279 |
| Dacnis viguieri | Viridian Dacnis | M | rum | greenish blue | blue | 0.17  2 | 0.22  3 | 0.330 | 0.275 | 0.223 |
| Diglossa  albilatera | White-sided  Flowerpiercer | M | rum | dark slate-grey | blue | 0.22  3 | 0.23  9 | 0.265 | 0.273 | 0.239 |
| Diglossa cyanea | Masked  Flowerpiercer | M | rum | dark blue | blue | 0.26  5 | 0.28  4 | 0.245 | 0.206 | 0.284 |
| Diglossa glauca | Deep-blue  Flowerpiercer | M | rum | deep blue | blue | 0.42  4 | 0.26  6 | 0.178 | 0.132 | 0.266 |
| Diglossa  indigotica | Indigo  Flowerpiercer | M | rum | indigo-blue | blue | 0.31  2 | 0.32  4 | 0.203 | 0.161 | 0.324 |
| Dubusia taeniata | Buff-breasted  Mountain-Tanager | M | rum | dark blue | blue | 0.22  7 | 0.24  7 | 0.263 | 0.263 | 0.247 |
| Iridosornis analis | Yellow-throated  Tanager | M | rum | greenish turquoise | blue | 0.17  5 | 0.23  0 | 0.303 | 0.292 | 0.230 |
| Iridosornis porphyrocephalu  s | Purplish-mantled Tanager | M | rum | dark blue | blue | 0.19  1 | 0.21  7 | 0.279 | 0.313 | 0.217 |

| Iridosornis  rufivertex | Golden-crowned  Tanager | M | rum | purplish-blue | blue | 0.29  2 | 0.27  6 | 0.221 | 0.211 | 0.276 |
| --- | --- | --- | --- | --- | --- | --- | --- | --- | --- | --- |
| Pipraeidea  melanonota | Fawn-breasted  Tanager | M | rum | bright turquoise  blue | blue | 0.35  6 | 0.35  1 | 0.170 | 0.123 | 0.351 |
| Porphyrospiza  caerulescens | Blue Finch | M | rum | bright indigo-blue | blue | 0.25  4 | 0.26  0 | 0.252 | 0.234 | 0.260 |
| Stephanophorus  diadematus | Diademed Tanager | M | rum | shining dark blue | blue | 0.30  0 | 0.24  8 | 0.226 | 0.226 | 0.248 |
| Tangara  labradorides | Metallic-green  Tanager | M | rum | opalescent blue-  green | blue | 0.11  0 | 0.24  5 | 0.390 | 0.256 | 0.245 |
| Tangara larvata | Golden-hooded  Tanager | M | rum | light blue | blue | 0.20  4 | 0.26  8 | 0.281 | 0.247 | 0.268 |
| Tangara  nigrocincta | Masked Tanager | M | rum | light blue | blue | 0.27  4 | 0.34  5 | 0.252 | 0.129 | 0.345 |
| Tangara  ruficervix | Golden-naped  Tanager | M | rum | bright blue | blue | 0.21  4 | 0.25  4 | 0.284 | 0.248 | 0.254 |
| Tangara vassorii | Blue-and-black  Tanager | M | rum | dark cobalt blue | blue | 0.43  8 | 0.29  5 | 0.159 | 0.108 | 0.295 |
| Tangara  viridicollis | Silvery Tanager | M | rum | pale turquoise | blue | 0.08  1 | 0.25  6 | 0.354 | 0.308 | 0.256 |
| Tangara  xanthocephala | Saffron-crowned  Tanager | M | rum | black / turquoise  green | blue | 0.08  7 | 0.23  6 | 0.419 | 0.258 | 0.236 |
| Tersina viridis | Swallow Tanager | M | rum | bright turquoise-  blue | blue | 0.23  9 | 0.26  6 | 0.320 | 0.175 | 0.266 |
| Thraupis  glaucocolpa | Glaucous Tanager | M | rum | smoky grey /  greenish | blue | 0.18  8 | 0.24  1 | 0.315 | 0.256 | 0.241 |
| Xenodacnis  parina | Tit-like Dacnis | M | rum | dark blue | blue | 0.27  7 | 0.29  7 | 0.244 | 0.182 | 0.297 |
| Chlorophanes  spiza | Green  Honeycreeper | M | thr | glossy viridian  green | blue | 0.05  1 | 0.30  2 | 0.444 | 0.203 | 0.302 |
| Cyanerpes  cyaneus | Red-legged  Honeycreeper | M | thr | purplish-blue | blue | 0.52  4 | 0.25  6 | 0.140 | 0.080 | 0.256 |
| Dacnis lineata | Black-faced Dacnis | M | thr | turquoise-blue | blue | 0.26  7 | 0.27  3 | 0.285 | 0.175 | 0.273 |
| Dacnis viguieri | Viridian Dacnis | M | thr | verditer blue/  green | blue | 0.27  6 | 0.21  7 | 0.334 | 0.173 | 0.217 |
| Diglossa  albilatera | White-sided  Flowerpiercer | M | thr | dark slate-grey | blue | 0.20  2 | 0.24  7 | 0.282 | 0.270 | 0.247 |
| Diglossa glauca | Deep-blue  Flowerpiercer | M | thr | deep blue | blue | 0.41  8 | 0.27  3 | 0.179 | 0.130 | 0.273 |
| Diglossa  indigotica | Indigo  Flowerpiercer | M | thr | indigo-blue | blue | 0.28  0 | 0.34  9 | 0.218 | 0.153 | 0.349 |
| Porphyrospiza  caerulescens | Blue Finch | M | thr | bright indigo-blue | blue | 0.27  7 | 0.28  3 | 0.234 | 0.207 | 0.283 |
| Tangara chilensis | Paradise Tanager | M | thr | dark blue | blue | 0.52  2 | 0.27  4 | 0.121 | 0.083 | 0.274 |
| Tangara  labradorides | Metallic-green  Tanager | M | thr | opalescent blue-  green | blue | 0.11  0 | 0.27  0 | 0.348 | 0.271 | 0.270 |
| Tangara  mexicana | Turquoise Tanager | M | thr | dark turquoise  blue | blue | 0.38  5 | 0.29  6 | 0.181 | 0.138 | 0.296 |
| Tangara  nigrocincta | Masked Tanager | M | thr | pale lavender | blue | 0.19  2 | 0.33  5 | 0.272 | 0.200 | 0.335 |
| Tangara  peruviana | Black-backed  Tanager | M | thr | bluish-green | blue | 0.09  0 | 0.22  5 | 0.388 | 0.297 | 0.225 |
| Tangara preciosa | Chestnut-backed  Tanager | M | thr | blue-green | blue | 0.08  1 | 0.23  1 | 0.394 | 0.294 | 0.231 |
| Tangara vassorii | Blue-and-black  Tanager | M | thr | dark blue | blue | 0.37  2 | 0.29  4 | 0.193 | 0.142 | 0.294 |
| Tangara velia | Opal-rumped  Tanager | M | thr | deep purplish-blue | blue | 0.29  3 | 0.34  2 | 0.204 | 0.161 | 0.342 |
| Thraupis abbas | Yellow-winged  Tanager | M | thr | blue | blue | 0.26  8 | 0.24  2 | 0.246 | 0.244 | 0.242 |
| Thraupis  bonariensis | Blue-and-yellow  Tanager | M | thr | dull blue | blue | 0.25  2 | 0.26  7 | 0.254 | 0.228 | 0.267 |
| Thraupis ornata | Golden-chevroned  Tanager | M | thr | shining cadet-blue | blue | 0.26  4 | 0.24  9 | 0.242 | 0.244 | 0.249 |
| Xenodacnis  parina | Tit-like Dacnis | M | thr | dark blue | blue | 0.26  4 | 0.29  1 | 0.248 | 0.198 | 0.291 |
| Chlorochrysa | Orange-eared | M | bre | emerald-green | blue | 0.08 | 0.24 | 0.493 | 0.181 | 0.245 |

| calliparaea | Tanager |  |  |  |  | 1 | 5 |  |  |  |
| --- | --- | --- | --- | --- | --- | --- | --- | --- | --- | --- |
| Chlorophanes  spiza | Green  Honeycreeper | M | bre | glossy viridian  green | blue | 0.05  7 | 0.32  5 | 0.416 | 0.201 | 0.325 |
| Cyanerpes  caeruleus | Purple  Honeycreeper | M | bre | lustrous violet-  blue | blue | 0.69  4 | 0.14  0 | 0.166 | 0.000 | 0.140 |
| Cyanerpes  cyaneus | Red-legged  Honeycreeper | M | bre | purplish-blue | blue | 0.60  2 | 0.19  7 | 0.139 | 0.062 | 0.197 |
| Cyanerpes  lucidus | Shining  Honeycreeper | M | bre | ultramarine blue | blue | 0.47  8 | 0.27  6 | 0.167 | 0.079 | 0.276 |
| Cyanerpes  nitidus | Short-billed  Honeycreeper | M | bre | purplish-blue | blue | 0.38  3 | 0.31  7 | 0.168 | 0.132 | 0.317 |
| Dacnis cayana | Blue Dacnis | M | bre | turquoise | blue | 0.20  6 | 0.28  5 | 0.269 | 0.241 | 0.285 |
| Dacnis hartlaubi | Turquoise Dacnis | M | bre | turquoise | blue | 0.27  9 | 0.27  3 | 0.281 | 0.168 | 0.273 |
| Dacnis lineata | Black-faced Dacnis | M | bre | turquoise-blue | blue | 0.25  3 | 0.28  5 | 0.291 | 0.171 | 0.285 |
| Dacnis nigripes | Black-legged  Dacnis | M | bre | light turquoise | blue | 0.21  8 | 0.28  8 | 0.293 | 0.202 | 0.288 |
| Dacnis viguieri | Viridian Dacnis | M | bre | verditer blue/  green | blue | 0.26  2 | 0.22  7 | 0.342 | 0.169 | 0.227 |
| Diglossa  albilatera | White-sided  Flowerpiercer | M | bre | dark slate-grey /  white | blue | 0.22  9 | 0.25  2 | 0.266 | 0.253 | 0.252 |
| Diglossa cyanea | Masked  Flowerpiercer | M | bre | dark blue | blue | 0.27  8 | 0.31  1 | 0.234 | 0.177 | 0.311 |
| Diglossa glauca | Deep-blue  Flowerpiercer | M | bre | deep blue | blue | 0.38  8 | 0.25  8 | 0.194 | 0.160 | 0.258 |
| Diglossa  indigotica | Indigo  Flowerpiercer | M | bre | indigo-blue | blue | 0.29  9 | 0.32  3 | 0.207 | 0.171 | 0.323 |
| Iridosornis  rufivertex | Golden-crowned  Tanager | M | bre | purplish-blue | blue | 0.27  0 | 0.27  9 | 0.231 | 0.220 | 0.279 |
| Stephanophorus  diadematus | Diademed Tanager | M | bre | shining dark blue | blue | 0.38  1 | 0.25  4 | 0.193 | 0.173 | 0.254 |
| Tangara chilensis | Paradise Tanager | M | bre | dark blue | blue | 0.16  4 | 0.25  2 | 0.181 | 0.403 | 0.252 |
| Tangara  cyanoventris | Gilt-edged  Tanager | M | bre | turquoise-blue | blue | 0.15  2 | 0.25  8 | 0.281 | 0.309 | 0.258 |
| Tangara gyrola | Bay-headed  Tanager | M | bre | green / light blue | blue | 0.22  4 | 0.27  1 | 0.344 | 0.161 | 0.271 |
| Tangara  labradorides | Metallic-green  Tanager | M | bre | opalescent blue-  green | blue | 0.10  6 | 0.28  2 | 0.379 | 0.233 | 0.282 |
| Tangara  mexicana | Turquoise Tanager | M | bre | dark turquoise  blue | blue | 0.31  2 | 0.20  9 | 0.250 | 0.229 | 0.209 |
| Tangara  peruviana | Black-backed  Tanager | M | bre | bluish-green | blue | 0.06  5 | 0.24  5 | 0.403 | 0.286 | 0.245 |
| Tangara preciosa | Chestnut-backed  Tanager | M | bre | blue-green | blue | 0.05  8 | 0.25  3 | 0.420 | 0.269 | 0.253 |
| Tangara  ruficervix | Golden-naped  Tanager | M | bre | bright blue | blue | 0.21  8 | 0.26  7 | 0.307 | 0.207 | 0.267 |
| Tangara seledon | Green-headed  Tanager | M | bre | turquoise | blue | 0.21  1 | 0.29  8 | 0.330 | 0.160 | 0.298 |
| Tangara vassorii | Blue-and-black  Tanager | M | bre | dark blue | blue | 0.39  4 | 0.30  7 | 0.172 | 0.127 | 0.307 |
| Tangara velia | Opal-rumped  Tanager | M | bre | purplish blue | blue | 0.42  5 | 0.26  5 | 0.163 | 0.148 | 0.265 |
| Tangara  xanthocephala | Saffron-crowned  Tanager | M | bre | black / turquoise  green | blue | 0.07  4 | 0.20  9 | 0.407 | 0.309 | 0.209 |
| Thraupis  glaucocolpa | Glaucous Tanager | M | bre | grey / turquoise-  blue | blue | 0.17  0 | 0.26  3 | 0.309 | 0.257 | 0.263 |
| Thraupis ornata | Golden-chevroned  Tanager | M | bre | shining cadet-blue | blue | 0.32  9 | 0.25  7 | 0.210 | 0.205 | 0.257 |
| Xenodacnis  parina | Tit-like Dacnis | M | bre | dark blue | blue | 0.31  5 | 0.31  2 | 0.210 | 0.163 | 0.312 |
| Chlorochrysa  calliparaea | Orange-eared  Tanager | M | bel | emerald-green /  dark blue-green | blue | 0.14  3 | 0.27  4 | 0.410 | 0.174 | 0.274 |
| Cyanerpes  caeruleus | Purple  Honeycreeper | M | bel | lustrous violet-  blue | blue | 0.65  2 | 0.19  8 | 0.146 | 0.004 | 0.198 |
| Cyanerpes  cyaneus | Red-legged  Honeycreeper | M | bel | purplish-blue | blue | 0.51  9 | 0.27  1 | 0.142 | 0.069 | 0.271 |

| Cyanerpes  lucidus | Shining  Honeycreeper | M | bel | ultramarine blue | blue | 0.51  3 | 0.26  2 | 0.159 | 0.067 | 0.262 |
| --- | --- | --- | --- | --- | --- | --- | --- | --- | --- | --- |
| Dacnis cayana | Blue Dacnis | M | bel | turquoise | blue | 0.23  5 | 0.28  5 | 0.285 | 0.195 | 0.285 |
| Dacnis hartlaubi | Turquoise Dacnis | M | bel | turquoise | blue | 0.27  0 | 0.27  9 | 0.280 | 0.172 | 0.279 |
| Dacnis nigripes | Black-legged  Dacnis | M | bel | light turquoise | blue | 0.24  4 | 0.28  9 | 0.299 | 0.168 | 0.289 |
| Dacnis viguieri | Viridian Dacnis | M | bel | verditer blue/  green | blue | 0.24  3 | 0.22  4 | 0.345 | 0.188 | 0.224 |
| Diglossa  albilatera | White-sided  Flowerpiercer | M | bel | dark slate-grey | blue | 0.22  1 | 0.24  7 | 0.268 | 0.265 | 0.247 |
| Diglossa cyanea | Masked  Flowerpiercer | M | bel | dark blue | blue | 0.24  6 | 0.28  1 | 0.255 | 0.218 | 0.281 |
| Diglossa glauca | Deep-blue  Flowerpiercer | M | bel | deep blue | blue | 0.29  5 | 0.25  8 | 0.232 | 0.215 | 0.258 |
| Diglossa  indigotica | Indigo  Flowerpiercer | M | bel | indigo-blue | blue | 0.29  7 | 0.32  1 | 0.214 | 0.168 | 0.321 |
| Iridosornis  rufivertex | Golden-crowned  Tanager | M | bel | purplish-blue/  chesnut | blue | 0.21  4 | 0.26  6 | 0.263 | 0.257 | 0.266 |
| Stephanophorus  diadematus | Diademed Tanager | M | bel | no | blue | 0.35  7 | 0.24  6 | 0.204 | 0.193 | 0.246 |
| Tangara  cyanicollis | Blue-necked  Tanager | M | bel | black / deep blue | blue | 0.27  6 | 0.25  5 | 0.242 | 0.227 | 0.255 |
| Tangara gyrola | Bay-headed  Tanager | M | bel | green / light blue | blue | 0.21  5 | 0.22  0 | 0.381 | 0.184 | 0.220 |
| Tangara lavinia | Rufous-winged  Tanager | M | bel | blue | blue | 0.22  9 | 0.26  9 | 0.348 | 0.154 | 0.269 |
| Tangara preciosa | Chestnut-backed  Tanager | M | bel | blue-green /  yellowish-opal | blue | 0.14  5 | 0.25  6 | 0.316 | 0.283 | 0.256 |
| Tangara vassorii | Blue-and-black  Tanager | M | bel | dark blue | blue | 0.38  1 | 0.30  5 | 0.169 | 0.146 | 0.305 |
| Thraupis ornata | Golden-chevroned  Tanager | M | bel | grey / blueish-  green | blue | 0.21  8 | 0.24  4 | 0.264 | 0.273 | 0.244 |
| Xenodacnis  parina | Tit-like Dacnis | M | bel | dark blue | blue | 0.25  2 | 0.28  8 | 0.252 | 0.208 | 0.288 |
| Anisognathus  lacrymosus | Lacrimose  Mountain-Tanager | F | cro | slaty blue | blue | 0.21  5 | 0.27  3 | 0.268 | 0.245 | 0.273 |
| Anisognathus  melanogenys | Black-cheeked  Mountain-Tanager | F | cro | shining blue | blue | 0.17  0 | 0.23  9 | 0.300 | 0.291 | 0.239 |
| Buthraupis  eximia | Black-chested  Mountain-Tanager | F | cro | blue | blue | 0.21  9 | 0.27  3 | 0.264 | 0.244 | 0.273 |
| Conirostrum  albifrons | Capped Conebill | F | cro | dull blue | blue | 0.28  8 | 0.28  5 | 0.223 | 0.205 | 0.285 |
| Conirostrum  bicolor | Bicolored Conebill | F | cro | pale bluish-grey | blue | 0.13  0 | 0.18  5 | 0.326 | 0.359 | 0.185 |
| Cyanicterus  cyanicterus | Blue-backed  Tanager | F | cro | bright cadet-blue | blue | 0.21  6 | 0.20  8 | 0.335 | 0.241 | 0.208 |
| Dacnis cayana | Blue Dacnis | F | cro | blue | blue | 0.21  8 | 0.24  5 | 0.329 | 0.209 | 0.245 |
| Dacnis nigripes | Black-legged  Dacnis | F | cro | brownish olive /  turquoise | blue | 0.17  5 | 0.23  5 | 0.308 | 0.283 | 0.235 |
| Dacnis venusta | Scarlet-thighed  Dacnis | F | cro | greenish olive /  blue | blue | 0.16  8 | 0.22  0 | 0.321 | 0.291 | 0.220 |
| Diglossa cyanea | Masked  Flowerpiercer | F | cro | dark blue | blue | 0.21  4 | 0.27  5 | 0.271 | 0.240 | 0.275 |
| Diglossa  indigotica | Indigo  Flowerpiercer | F | cro | indigo-blue | blue | 0.18  7 | 0.23  1 | 0.288 | 0.294 | 0.231 |
| Iridosornis analis | Yellow-throated  Tanager | F | cro | deep purplish blue | blue | 0.15  2 | 0.23  7 | 0.299 | 0.312 | 0.237 |
| Iridosornis porphyrocephalu  s | Purplish-mantled Tanager | F | cro | dark blue | blue | 0.21  2 | 0.23  8 | 0.267 | 0.283 | 0.238 |
| Tangara  cyanocephala | Red-necked  Tanager | F | cro | blue-violet | blue | 0.25  5 | 0.34  3 | 0.237 | 0.165 | 0.343 |
| Tangara  nigrocincta | Masked Tanager | F | cro | pale lavender | blue | 0.22  8 | 0.32  3 | 0.255 | 0.194 | 0.323 |
| Tangara seledon | Green-headed  Tanager | F | cro | turquoise-green | blue | 0.07  7 | 0.22  7 | 0.443 | 0.252 | 0.227 |

| Tangara vassorii | Blue-and-black  Tanager | F | cro | dark cobalt blue | blue | 0.26  1 | 0.29  1 | 0.242 | 0.207 | 0.291 |
| --- | --- | --- | --- | --- | --- | --- | --- | --- | --- | --- |
| Thraupis abbas | Yellow-winged  Tanager | F | cro | campanula-blue | blue | 0.32  1 | 0.24  6 | 0.220 | 0.213 | 0.246 |
| Thraupis  bonariensis | Blue-and-yellow  Tanager | F | cro | brownish olive /  blue | blue | 0.15  4 | 0.20  1 | 0.313 | 0.331 | 0.201 |
| Thraupis  cyanocephala | Blue-capped  Tanager | F | cro | shining  cornflower-blue | blue | 0.36  2 | 0.27  0 | 0.200 | 0.167 | 0.270 |
| Thraupis ornata | Golden-chevroned  Tanager | F | cro | shining cadet-blue | blue | 0.21  7 | 0.26  9 | 0.265 | 0.250 | 0.269 |
| Anisognathus  melanogenys | Black-cheeked  Mountain-Tanager | F | nap | shining blue | blue | 0.21  8 | 0.25  0 | 0.271 | 0.262 | 0.250 |
| Buthraupis  eximia | Black-chested  Mountain-Tanager | F | nap | blue | blue | 0.33  6 | 0.30  7 | 0.197 | 0.160 | 0.307 |
| Buthraupis  montana | Hooded  Mountain-Tanager | F | nap | shining pale milky  blue | blue | 0.22  1 | 0.24  0 | 0.275 | 0.264 | 0.240 |
| Conirostrum  bicolor | Bicolored Conebill | F | nap | pale bluish-grey | blue | 0.14  2 | 0.19  4 | 0.316 | 0.348 | 0.194 |
| Cyanicterus  cyanicterus | Blue-backed  Tanager | F | nap | bright cadet-blue | blue | 0.24  1 | 0.22  2 | 0.311 | 0.226 | 0.222 |
| Dacnis venusta | Scarlet-thighed  Dacnis | F | nap | greenish olive /  blue | blue | 0.17  7 | 0.22  6 | 0.338 | 0.259 | 0.226 |
| Diglossa cyanea | Masked  Flowerpiercer | F | nap | dark blue | blue | 0.24  0 | 0.28  7 | 0.257 | 0.216 | 0.287 |
| Diglossa  indigotica | Indigo  Flowerpiercer | F | nap | indigo-blue | blue | 0.20  5 | 0.23  4 | 0.279 | 0.283 | 0.234 |
| Iridosornis analis | Yellow-throated  Tanager | F | nap | greenish turquoise | blue | 0.17  1 | 0.26  1 | 0.289 | 0.279 | 0.261 |
| Iridosornis porphyrocephalu  s | Purplish-mantled Tanager | F | nap | dark blue | blue | 0.28  0 | 0.28  9 | 0.230 | 0.201 | 0.289 |
| Tangara  cyanocephala | Red-necked  Tanager | F | nap | blue-violet | blue | 0.24  3 | 0.23  3 | 0.301 | 0.224 | 0.233 |
| Tangara  nigrocincta | Masked Tanager | F | nap | pale lavender | blue | 0.24  9 | 0.33  2 | 0.249 | 0.169 | 0.332 |
| Tangara seledon | Green-headed  Tanager | F | nap | turquoise-green | blue | 0.09  0 | 0.19  4 | 0.455 | 0.260 | 0.194 |
| Tangara vassorii | Blue-and-black  Tanager | F | nap | dark cobalt blue | blue | 0.28  8 | 0.30  2 | 0.222 | 0.189 | 0.302 |
| Thraupis abbas | Yellow-winged  Tanager | F | nap | campanula-blue | blue | 0.36  0 | 0.24  6 | 0.204 | 0.190 | 0.246 |
| Thraupis  cyanocephala | Blue-capped  Tanager | F | nap | shining  cornflower-blue | blue | 0.37  7 | 0.24  6 | 0.203 | 0.174 | 0.246 |
| Thraupis ornata | Golden-chevroned  Tanager | F | nap | shining cadet-blue | blue | 0.27  0 | 0.27  8 | 0.234 | 0.217 | 0.278 |
| Anisognathus  lacrymosus | Lacrimose  Mountain-Tanager | F | man | slaty blue | blue | 0.19  4 | 0.24  7 | 0.281 | 0.279 | 0.247 |
| Anisognathus  melanogenys | Black-cheeked  Mountain-Tanager | F | man | dark grey-blue | blue | 0.19  9 | 0.23  6 | 0.281 | 0.284 | 0.236 |
| Buthraupis  montana | Hooded  Mountain-Tanager | F | man | shining dark blue | blue | 0.64  3 | 0.22  0 | 0.137 | 0.000 | 0.220 |
| Conirostrum  bicolor | Bicolored Conebill | F | man | pale bluish-grey | blue | 0.14  6 | 0.20  2 | 0.316 | 0.337 | 0.202 |
| Diglossa cyanea | Masked  Flowerpiercer | F | man | dark blue | blue | 0.22  2 | 0.26  2 | 0.266 | 0.250 | 0.262 |
| Diglossa  indigotica | Indigo  Flowerpiercer | F | man | indigo-blue | blue | 0.19  8 | 0.23  3 | 0.281 | 0.289 | 0.233 |
| Dubusia taeniata | Buff-breasted  Mountain-Tanager | F | man | black / dark blue | blue | 0.21  5 | 0.25  5 | 0.274 | 0.256 | 0.255 |
| Iridosornis analis | Yellow-throated  Tanager | F | man | greenish turquoise | blue | 0.17  5 | 0.23  7 | 0.297 | 0.292 | 0.237 |
| Iridosornis porphyrocephalu  s | Purplish-mantled Tanager | F | man | dark blue | blue | 0.22  4 | 0.24  9 | 0.266 | 0.260 | 0.249 |
| Stephanophorus  diadematus | Diademed Tanager | F | man | shining dark blue | blue | 0.33  0 | 0.24  5 | 0.217 | 0.208 | 0.245 |
| Tangara  labradorides | Metallic-green  Tanager | F | man | opalescent blue-  green | blue | 0.05  9 | 0.25  7 | 0.426 | 0.258 | 0.257 |
| Tangara vassorii | Blue-and-black | F | man | dark cobalt blue | blue | 0.23 | 0.26 | 0.261 | 0.246 | 0.261 |

|  | Tanager |  |  |  |  | 2 | 1 |  |  |  |
| --- | --- | --- | --- | --- | --- | --- | --- | --- | --- | --- |
| Tangara  xanthocephala | Saffron-crowned  Tanager | F | man | black / turquoise  green | blue | 0.17  0 | 0.22  7 | 0.313 | 0.289 | 0.227 |
| Thraupis abbas | Yellow-winged  Tanager | F | man | smoky blue-grey | blue | 0.32  0 | 0.18  9 | 0.245 | 0.245 | 0.189 |
| Thraupis  glaucocolpa | Glaucous Tanager | F | man | smoky grey /  greenish | blue | 0.18  0 | 0.24  9 | 0.297 | 0.275 | 0.249 |
| Anisognathus  igniventris | Scarlet-bellied  Mountain-Tanager | F | rum | blue | blue | 0.34  2 | 0.32  5 | 0.193 | 0.140 | 0.325 |
| Anisognathus  lacrymosus | Lacrimose  Mountain-Tanager | F | rum | slaty blue | blue | 0.28  2 | 0.27  8 | 0.234 | 0.206 | 0.278 |
| Anisognathus  melanogenys | Black-cheeked  Mountain-Tanager | F | rum | dark grey-blue | blue | 0.20  4 | 0.23  9 | 0.286 | 0.270 | 0.239 |
| Buthraupis  montana | Hooded  Mountain-Tanager | F | rum | shining dark blue | blue | 0.40  7 | 0.23  0 | 0.201 | 0.162 | 0.230 |
| Conirostrum  bicolor | Bicolored Conebill | F | rum | pale bluish-grey | blue | 0.15  8 | 0.21  2 | 0.299 | 0.331 | 0.212 |
| Cyanicterus  cyanicterus | Blue-backed  Tanager | F | rum | bright cadet-blue | blue | 0.23  4 | 0.21  9 | 0.296 | 0.252 | 0.219 |
| Dacnis nigripes | Black-legged  Dacnis | F | rum | brownish olive /  turquoise | blue | 0.18  1 | 0.24  2 | 0.315 | 0.262 | 0.242 |
| Diglossa cyanea | Masked  Flowerpiercer | F | rum | dark blue | blue | 0.22  9 | 0.26  2 | 0.265 | 0.245 | 0.262 |
| Diglossa  indigotica | Indigo  Flowerpiercer | F | rum | indigo-blue | blue | 0.19  8 | 0.22  0 | 0.273 | 0.309 | 0.220 |
| Dubusia taeniata | Buff-breasted  Mountain-Tanager | F | rum | dark blue | blue | 0.22  3 | 0.23  3 | 0.267 | 0.276 | 0.233 |
| Iridosornis analis | Yellow-throated  Tanager | F | rum | greenish turquoise | blue | 0.13  0 | 0.23  0 | 0.336 | 0.304 | 0.230 |
| Iridosornis porphyrocephalu  s | Purplish-mantled Tanager | F | rum | dark blue | blue | 0.17  2 | 0.23  8 | 0.310 | 0.279 | 0.238 |
| Stephanophorus  diadematus | Diademed Tanager | F | rum | shining dark blue | blue | 0.34  4 | 0.26  7 | 0.200 | 0.188 | 0.267 |
| Tangara  labradorides | Metallic-green  Tanager | F | rum | opalescent blue-  green | blue | 0.05  8 | 0.22  1 | 0.458 | 0.263 | 0.221 |
| Tangara larvata | Golden-hooded  Tanager | F | rum | light blue | blue | 0.21  1 | 0.27  8 | 0.311 | 0.200 | 0.278 |
| Tangara  nigrocincta | Masked Tanager | F | rum | light blue | blue | 0.24  4 | 0.30  0 | 0.294 | 0.163 | 0.300 |
| Tangara vassorii | Blue-and-black  Tanager | F | rum | dark cobalt blue | blue | 0.21  8 | 0.25  3 | 0.264 | 0.265 | 0.253 |
| Tangara  xanthocephala | Saffron-crowned  Tanager | F | rum | black / turquoise  green | blue | 0.13  8 | 0.21  6 | 0.352 | 0.294 | 0.216 |
| Thraupis  glaucocolpa | Glaucous Tanager | F | rum | smoky grey /  greenish | blue | 0.19  0 | 0.25  9 | 0.307 | 0.245 | 0.259 |
| Diglossa  indigotica | Indigo  Flowerpiercer | F | thr | indigo-blue | blue | 0.12  1 | 0.20  1 | 0.312 | 0.366 | 0.201 |
| Tangara cayana | Burnished-buff  Tanager | F | thr | blue-violet | blue | 0.10  5 | 0.25  0 | 0.326 | 0.319 | 0.250 |
| Tangara chilensis | Paradise Tanager | F | thr | dark blue | blue | 0.46  3 | 0.27  6 | 0.159 | 0.102 | 0.276 |
| Tangara  labradorides | Metallic-green  Tanager | F | thr | opalescent blue-  green | blue | 0.13  3 | 0.24  5 | 0.338 | 0.284 | 0.245 |
| Tangara  mexicana | Turquoise Tanager | F | thr | dark turquoise  blue | blue | 0.40  2 | 0.27  2 | 0.180 | 0.145 | 0.272 |
| Tangara  nigrocincta | Masked Tanager | F | thr | pale lavender | blue | 0.12  2 | 0.27  9 | 0.344 | 0.254 | 0.279 |
| Tangara vassorii | Blue-and-black  Tanager | F | thr | dark blue | blue | 0.21  9 | 0.26  8 | 0.261 | 0.253 | 0.268 |
| Tangara velia | Opal-rumped  Tanager | F | thr | deep purplish-blue | blue | 0.23  2 | 0.39  1 | 0.224 | 0.153 | 0.391 |
| Tangara  xanthocephala | Saffron-crowned  Tanager | F | thr | black / turquoise  green | blue | 0.13  8 | 0.21  2 | 0.317 | 0.333 | 0.212 |
| Thraupis abbas | Yellow-winged  Tanager | F | thr | blue | blue | 0.23  5 | 0.24  0 | 0.261 | 0.264 | 0.240 |
| Thraupis ornata | Golden-chevroned  Tanager | F | thr | shining cadet-blue | blue | 0.18  0 | 0.24  0 | 0.287 | 0.292 | 0.240 |

| Diglossa cyanea | Masked  Flowerpiercer | F | bre | dark blue | blue | 0.18  7 | 0.24  7 | 0.282 | 0.283 | 0.247 |
| --- | --- | --- | --- | --- | --- | --- | --- | --- | --- | --- |
| Diglossa  indigotica | Indigo  Flowerpiercer | F | bre | indigo-blue | blue | 0.17  8 | 0.21  6 | 0.281 | 0.324 | 0.216 |
| Stephanophorus  diadematus | Diademed Tanager | F | bre | shining dark blue | blue | 0.28  8 | 0.25  4 | 0.234 | 0.224 | 0.254 |
| Tangara chilensis | Paradise Tanager | F | bre | dark blue | blue | 0.12  5 | 0.23  0 | 0.210 | 0.435 | 0.230 |
| Tangara  cyanoventris | Gilt-edged  Tanager | F | bre | turquoise-blue | blue | 0.18  3 | 0.24  5 | 0.341 | 0.232 | 0.245 |
| Tangara gyrola | Bay-headed  Tanager | F | bre | green / light blue | blue | 0.21  3 | 0.26  0 | 0.369 | 0.157 | 0.260 |
| Tangara  labradorides | Metallic-green  Tanager | F | bre | opalescent blue-  green | blue | 0.12  4 | 0.24  0 | 0.349 | 0.287 | 0.240 |
| Tangara  mexicana | Turquoise Tanager | F | bre | dark turquoise  blue | blue | 0.28  7 | 0.19  9 | 0.261 | 0.253 | 0.199 |
| Tangara seledon | Green-headed  Tanager | F | bre | turquoise | blue | 0.18  4 | 0.25  1 | 0.365 | 0.201 | 0.251 |
| Tangara vassorii | Blue-and-black  Tanager | F | bre | dark blue | blue | 0.25  5 | 0.30  0 | 0.240 | 0.205 | 0.300 |
| Tangara velia | Opal-rumped  Tanager | F | bre | purplish blue | blue | 0.34  8 | 0.32  8 | 0.171 | 0.152 | 0.328 |
| Tangara  xanthocephala | Saffron-crowned  Tanager | F | bre | black / turquoise  green | blue | 0.12  7 | 0.20  2 | 0.334 | 0.337 | 0.202 |
| Thraupis  glaucocolpa | Glaucous Tanager | F | bre | grey / turquoise-  blue | blue | 0.18  4 | 0.27  9 | 0.309 | 0.228 | 0.279 |
| Thraupis ornata | Golden-chevroned  Tanager | F | bre | shining cadet-blue | blue | 0.25  6 | 0.24  6 | 0.244 | 0.254 | 0.246 |
| Diglossa cyanea | Masked  Flowerpiercer | F | bel | dark blue | blue | 0.19  3 | 0.24  4 | 0.282 | 0.281 | 0.244 |
| Diglossa  indigotica | Indigo  Flowerpiercer | F | bel | indigo-blue | blue | 0.19  1 | 0.22  2 | 0.275 | 0.312 | 0.222 |
| Stephanophorus  diadematus | Diademed Tanager | F | bel | no | blue | 0.24  2 | 0.24  3 | 0.255 | 0.259 | 0.243 |
| Tangara  cyanicollis | Blue-necked  Tanager | F | bel | black / deep blue | blue | 0.26  3 | 0.29  0 | 0.232 | 0.214 | 0.290 |
| Tangara gyrola | Bay-headed  Tanager | F | bel | green / light blue | blue | 0.20  4 | 0.20  2 | 0.358 | 0.235 | 0.202 |
| Tangara lavinia | Rufous-winged  Tanager | F | bel | blue | blue | 0.17  6 | 0.23  3 | 0.323 | 0.267 | 0.233 |
| Tangara vassorii | Blue-and-black  Tanager | F | bel | dark blue | blue | 0.27  1 | 0.30  6 | 0.229 | 0.194 | 0.306 |
| Thraupis ornata | Golden-chevroned  Tanager | F | bel | grey / blueish-  green | blue | 0.14  9 | 0.22  8 | 0.298 | 0.325 | 0.228 |
| Cnemoscopus  rubrirostris | Gray-hooded Bush  Tanager | M | cro | medium grey | grey | 0.13  1 | 0.24  1 | 0.312 | 0.317 | 0.241 |
| Coereba flaveola | Bananaquit | M | cro | dark grey | grey | 0.18  2 | 0.24  1 | 0.292 | 0.285 | 0.241 |
| Diuca diuca | Common Diuca-  Finch | M | cro | dark gray | grey | 0.16  1 | 0.24  9 | 0.294 | 0.296 | 0.249 |
| Hemispingus  melanotis | Black-eared  Hemispingus | M | cro | grey | grey | 0.18  3 | 0.24  8 | 0.288 | 0.281 | 0.248 |
| Hemispingus reyi | Gray-capped  Hemispingus | M | cro | grey | grey | 0.16  7 | 0.24  3 | 0.291 | 0.299 | 0.243 |
| Hemispingus xanthophthalmu  s | Drab Hemispingus | M | cro | brownish-grey | grey | 0.12  0 | 0.20  8 | 0.310 | 0.362 | 0.208 |
| Idiopsar  brachyurus | Short-tailed Finch | M | cro | leaden gray | grey | 0.17  8 | 0.24  3 | 0.284 | 0.295 | 0.243 |
| Incaspiza laeta | Buff-bridled Inca-  Finch | M | cro | grey | grey | 0.17  2 | 0.24  2 | 0.288 | 0.299 | 0.242 |
| Incaspiza  personata | Rufous-backed  Inca-Finch | M | cro | grey | grey | 0.14  0 | 0.23  0 | 0.300 | 0.330 | 0.230 |
| Melanodera  melanodera | White-bridled  Finch | M | cro | grey | grey | 0.19  0 | 0.25  0 | 0.283 | 0.277 | 0.250 |
| Neothraupis  fasciata | White-banded  Tanager | M | cro | grey | grey | 0.17  4 | 0.25  0 | 0.290 | 0.285 | 0.250 |
| Phrygilus  fruticeti | Mourning Sierra-  Finch | M | cro | grey | grey | 0.18  2 | 0.24  2 | 0.286 | 0.289 | 0.242 |

| Piezorhina  cinerea | Cinereous Finch | M | cro | pale grey | grey | 0.14  8 | 0.24  0 | 0.297 | 0.314 | 0.240 |
| --- | --- | --- | --- | --- | --- | --- | --- | --- | --- | --- |
| Poospiza alticola | Plain-tailed  Warbling-Finch | M | cro | gray brown | grey | 0.15  0 | 0.23  7 | 0.297 | 0.315 | 0.237 |
| Poospiza  cabanisi | Gray-throated  Warbling-Finch | M | cro | grey | grey | 0.14  0 | 0.22  9 | 0.308 | 0.323 | 0.229 |
| Poospiza caesar | Chestnut-breasted  Mountain-Finch | M | cro | gray | grey | 0.16  4 | 0.23  2 | 0.292 | 0.312 | 0.232 |
| Poospiza  hispaniolensis | Collared Warbling-  Finch | M | cro | grey / black | grey | 0.18  0 | 0.25  3 | 0.287 | 0.280 | 0.253 |
| Poospiza  nigrorufa | Black-and-rufous  Warbling-Finch | M | cro | blackish grey | grey | 0.18  2 | 0.23  6 | 0.286 | 0.296 | 0.236 |
| Poospiza  thoracica | Bay-chested  Warbling-Finch | M | cro | grey | grey | 0.12  2 | 0.25  0 | 0.315 | 0.313 | 0.250 |
| Saltator  coerulescens | Grayish Saltator | M | cro | greyish | grey | 0.14  6 | 0.23  4 | 0.299 | 0.322 | 0.234 |
| Sporophila  caerulescens | Double-collared  Seedeater | M | cro | lead-grey | grey | 0.16  8 | 0.23  8 | 0.288 | 0.306 | 0.238 |
| Sporophila  hypoxantha | Tawny-bellied  Seedeater | M | cro | greyish | grey | 0.17  9 | 0.25  0 | 0.285 | 0.286 | 0.250 |
| Sporophila  leucoptera | White-bellied  Seedeater | M | cro | dark grey | grey | 0.16  3 | 0.23  9 | 0.293 | 0.305 | 0.239 |
| Sporophila  minuta | Ruddy-breasted  Seedeater | M | cro | mid-grey | grey | 0.12  8 | 0.23  3 | 0.311 | 0.328 | 0.233 |
| Sporophila  palustris | Marsh Seedeater | M | cro | grey | grey | 0.16  3 | 0.24  5 | 0.291 | 0.301 | 0.245 |
| Sporophila  peruviana | Parrot-billed  Seedeater | M | cro | greyish | grey | 0.15  9 | 0.23  2 | 0.298 | 0.311 | 0.232 |
| Sporophila  plumbea | Plumbeous  Seedeater | M | cro | lead-grey | grey | 0.16  2 | 0.24  7 | 0.291 | 0.300 | 0.247 |
| Sporophila  ruficollis | Dark-throated  Seedeater | M | cro | grey | grey | 0.16  2 | 0.24  0 | 0.291 | 0.307 | 0.240 |
| Sporophila  telasco | Chestnut-throated  Seedeater | M | cro | mid-grey / dusky | grey | 0.14  2 | 0.23  4 | 0.303 | 0.321 | 0.234 |
| Tangara inornata | Plain-colored  Tanager | M | cro | dark gray | grey | 0.17  5 | 0.25  0 | 0.282 | 0.293 | 0.250 |
| Thraupis  glaucocolpa | Glaucous Tanager | M | cro | smoky grey /  greenish | grey | 0.17  0 | 0.23  3 | 0.300 | 0.297 | 0.233 |
| Thraupis sayaca | Sayaca Tanager | M | cro | dull grey / bluish | grey | 0.17  2 | 0.24  0 | 0.293 | 0.295 | 0.240 |
| Coereba flaveola | Bananaquit | M | nap | dark grey | grey | 0.19  1 | 0.24  0 | 0.285 | 0.284 | 0.240 |
| Coryphaspiza  melanotis | Black-masked  Finch | M | nap | greyish | grey | 0.14  4 | 0.22  0 | 0.297 | 0.338 | 0.220 |
| Creurgops  verticalis | Rufous-crested  Tanager | M | nap | leaden grey | grey | 0.12  9 | 0.19  6 | 0.297 | 0.378 | 0.196 |
| Diuca diuca | Common Diuca-  Finch | M | nap | dark gray | grey | 0.18  8 | 0.25  1 | 0.281 | 0.280 | 0.251 |
| Hemispingus  melanotis | Black-eared  Hemispingus | M | nap | grey | grey | 0.19  2 | 0.24  8 | 0.283 | 0.277 | 0.248 |
| Hemispingus xanthophthalmu  s | Drab Hemispingus | M | nap | brownish-grey | grey | 0.14  3 | 0.21  4 | 0.297 | 0.346 | 0.214 |
| Idiopsar  brachyurus | Short-tailed Finch | M | nap | leaden gray | grey | 0.20  2 | 0.24  2 | 0.276 | 0.281 | 0.242 |
| Incaspiza laeta | Buff-bridled Inca-  Finch | M | nap | grey | grey | 0.17  2 | 0.24  6 | 0.288 | 0.294 | 0.246 |
| Melanodera  xanthogramma | Yellow-bridled  Finch | M | nap | grey | grey | 0.23  4 | 0.23  1 | 0.273 | 0.263 | 0.231 |
| Neothraupis  fasciata | White-banded  Tanager | M | nap | grey | grey | 0.19  1 | 0.25  0 | 0.279 | 0.279 | 0.250 |
| Phrygilus  fruticeti | Mourning Sierra-  Finch | M | nap | grey | grey | 0.17  1 | 0.24  1 | 0.291 | 0.297 | 0.241 |
| Piezorhina  cinerea | Cinereous Finch | M | nap | pale grey | grey | 0.16  2 | 0.24  4 | 0.290 | 0.305 | 0.244 |
| Poospiza alticola | Plain-tailed  Warbling-Finch | M | nap | gray brown | grey | 0.16  3 | 0.23  3 | 0.291 | 0.313 | 0.233 |
| Poospiza  cabanisi | Gray-throated  Warbling-Finch | M | nap | grey | grey | 0.12  2 | 0.19  7 | 0.318 | 0.363 | 0.197 |

| Poospiza caesar | Chestnut-breasted  Mountain-Finch | M | nap | gray | grey | 0.17  4 | 0.23  7 | 0.285 | 0.303 | 0.237 |
| --- | --- | --- | --- | --- | --- | --- | --- | --- | --- | --- |
| Poospiza  hispaniolensis | Collared Warbling-  Finch | M | nap | grey | grey | 0.19  8 | 0.25  1 | 0.280 | 0.271 | 0.251 |
| Poospiza  nigrorufa | Black-and-rufous  Warbling-Finch | M | nap | no | grey | 0.17  1 | 0.23  1 | 0.288 | 0.310 | 0.231 |
| Poospiza  thoracica | Bay-chested  Warbling-Finch | M | nap | grey | grey | 0.14  2 | 0.25  1 | 0.303 | 0.304 | 0.251 |
| Saltator  coerulescens | Grayish Saltator | M | nap | greyish | grey | 0.13  3 | 0.22  2 | 0.303 | 0.342 | 0.222 |
| Schistochlamys  melanopis | Black-faced  Tanager | M | nap | grey | grey | 0.20  1 | 0.25  8 | 0.278 | 0.263 | 0.258 |
| Sporophila  albogularis | White-throated  Seedeater | M | nap | no | grey | 0.18  3 | 0.24  2 | 0.281 | 0.294 | 0.242 |
| Sporophila  caerulescens | Double-collared  Seedeater | M | nap | olive-brown | grey | 0.16  5 | 0.23  9 | 0.289 | 0.308 | 0.239 |
| Sporophila  hypoxantha | Tawny-bellied  Seedeater | M | nap | greyish | grey | 0.17  8 | 0.24  9 | 0.284 | 0.290 | 0.249 |
| Sporophila  leucoptera | White-bellied  Seedeater | M | nap | dark grey | grey | 0.20  1 | 0.24  5 | 0.276 | 0.277 | 0.245 |
| Sporophila  minuta | Ruddy-breasted  Seedeater | M | nap | mid-grey | grey | 0.14  3 | 0.23  3 | 0.301 | 0.323 | 0.233 |
| Sporophila  palustris | Marsh Seedeater | M | nap | grey | grey | 0.18  0 | 0.23  9 | 0.282 | 0.299 | 0.239 |
| Sporophila  peruviana | Parrot-billed  Seedeater | M | nap | greyish-brown | grey | 0.17  1 | 0.23  2 | 0.289 | 0.308 | 0.232 |
| Sporophila  plumbea | Plumbeous  Seedeater | M | nap | lead-grey | grey | 0.16  3 | 0.24  7 | 0.288 | 0.301 | 0.247 |
| Sporophila  ruficollis | Dark-throated  Seedeater | M | nap | grey | grey | 0.18  5 | 0.24  7 | 0.281 | 0.287 | 0.247 |
| Sporophila  telasco | Chestnut-throated  Seedeater | M | nap | greyish / dusky | grey | 0.14  4 | 0.23  8 | 0.301 | 0.318 | 0.238 |
| Tangara inornata | Plain-colored  Tanager | M | nap | dark gray | grey | 0.19  4 | 0.24  9 | 0.279 | 0.278 | 0.249 |
| Thraupis  glaucocolpa | Glaucous Tanager | M | nap | smoky grey /  greenish | grey | 0.19  4 | 0.22  8 | 0.298 | 0.280 | 0.228 |
| Thraupis sayaca | Sayaca Tanager | M | nap | dull grey / bluish | grey | 0.18  6 | 0.23  8 | 0.294 | 0.282 | 0.238 |
| Charitospiza  eucosma | Coal-crested Finch | M | man | silvery gray | grey | 0.18  6 | 0.24  9 | 0.280 | 0.286 | 0.249 |
| Coereba flaveola | Bananaquit | M | man | dark grey | grey | 0.19  8 | 0.23  9 | 0.281 | 0.282 | 0.239 |
| Coryphospingus  pileatus | Pileated Finch | M | man | greyish | grey | 0.21  3 | 0.23  7 | 0.271 | 0.280 | 0.237 |
| Creurgops  verticalis | Rufous-crested  Tanager | M | man | leaden grey | grey | 0.18  1 | 0.24  3 | 0.284 | 0.292 | 0.243 |
| Diuca diuca | Common Diuca-  Finch | M | man | dark gray | grey | 0.19  7 | 0.24  0 | 0.276 | 0.286 | 0.240 |
| Hemispingus  verticalis | Black-headed  Hemispingus | M | man | dark grey | grey | 0.16  9 | 0.24  2 | 0.289 | 0.300 | 0.242 |
| Hemispingus xanthophthalmu  s | Drab Hemispingus | M | man | brownish-grey | grey | 0.13  5 | 0.21  1 | 0.304 | 0.349 | 0.211 |
| Idiopsar  brachyurus | Short-tailed Finch | M | man | leaden gray | grey | 0.17  3 | 0.23  9 | 0.286 | 0.302 | 0.239 |
| Melanodera  xanthogramma | Yellow-bridled  Finch | M | man | grey | grey | 0.22  1 | 0.21  3 | 0.284 | 0.282 | 0.213 |
| Neothraupis  fasciata | White-banded  Tanager | M | man | grey | grey | 0.18  4 | 0.24  9 | 0.282 | 0.284 | 0.249 |
| Paroaria  coronata | Red-crested  Cardinal | M | man | grey | grey | 0.19  6 | 0.25  3 | 0.276 | 0.275 | 0.253 |
| Phrygilus  fruticeti | Mourning Sierra-  Finch | M | man | grey | grey | 0.18  7 | 0.24  4 | 0.284 | 0.284 | 0.244 |
| Piezorhina  cinerea | Cinereous Finch | M | man | pale grey | grey | 0.17  0 | 0.24  7 | 0.286 | 0.298 | 0.247 |
| Poospiza alticola | Plain-tailed  Warbling-Finch | M | man | gray brown | grey | 0.17  7 | 0.24  2 | 0.285 | 0.297 | 0.242 |
| Poospiza caesar | Chestnut-breasted  Mountain-Finch | M | man | gray | grey | 0.20  3 | 0.24  1 | 0.274 | 0.282 | 0.241 |

| Poospiza  melanoleuca | Black-capped  Warbling-Finch | M | man | grey | grey | 0.17  6 | 0.25  1 | 0.289 | 0.285 | 0.251 |
| --- | --- | --- | --- | --- | --- | --- | --- | --- | --- | --- |
| Poospiza  nigrorufa | Black-and-rufous  Warbling-Finch | M | man | blue-grey | grey | 0.18  7 | 0.22  8 | 0.281 | 0.304 | 0.228 |
| Poospiza  thoracica | Bay-chested  Warbling-Finch | M | man | grey | grey | 0.13  1 | 0.23  1 | 0.309 | 0.329 | 0.231 |
| Poospiza  torquata | Ringed Warbling-  Finch | M | man | grey | grey | 0.20  7 | 0.24  8 | 0.275 | 0.271 | 0.248 |
| Saltator  aurantiirostris | Golden-billed  Saltator | M | man | grey | grey | 0.16  0 | 0.23  8 | 0.295 | 0.307 | 0.238 |
| Saltator  coerulescens | Grayish Saltator | M | man | greyish | grey | 0.14  1 | 0.22  7 | 0.302 | 0.330 | 0.227 |
| Saltator  nigriceps | Black-cowled  Saltator | M | man | deep gray | grey | 0.17  3 | 0.24  8 | 0.287 | 0.291 | 0.248 |
| Schistochlamys  melanopis | Black-faced  Tanager | M | man | grey | grey | 0.21  0 | 0.25  7 | 0.274 | 0.259 | 0.257 |
| Sicalis  taczanowskii | Sulphur-throated  Finch | M | man | no | grey | 0.14  3 | 0.22  9 | 0.298 | 0.330 | 0.229 |
| Sporophila  albogularis | White-throated  Seedeater | M | man | dark greyish | grey | 0.19  9 | 0.24  7 | 0.276 | 0.279 | 0.247 |
| Sporophila  caerulescens | Double-collared  Seedeater | M | man | olive-brown | grey | 0.15  0 | 0.23  2 | 0.295 | 0.322 | 0.232 |
| Sporophila  hypoxantha | Tawny-bellied  Seedeater | M | man | greyish | grey | 0.18  4 | 0.24  6 | 0.280 | 0.289 | 0.246 |
| Sporophila  minuta | Ruddy-breasted  Seedeater | M | man | mid-grey | grey | 0.14  0 | 0.22  7 | 0.303 | 0.330 | 0.227 |
| Sporophila  palustris | Marsh Seedeater | M | man | grey | grey | 0.17  4 | 0.24  4 | 0.284 | 0.298 | 0.244 |
| Sporophila  plumbea | Plumbeous  Seedeater | M | man | lead-grey | grey | 0.17  9 | 0.24  5 | 0.282 | 0.294 | 0.245 |
| Tangara inornata | Plain-colored  Tanager | M | man | dark gray | grey | 0.18  8 | 0.25  7 | 0.282 | 0.274 | 0.257 |
| Thlypopsis  fulviceps | Fulvous-headed  Tanager | M | man | grey | grey | 0.19  1 | 0.24  0 | 0.282 | 0.287 | 0.240 |
| Thlypopsis  inornata | Buff-bellied  Tanager | M | man | olive-grey | grey | 0.16  1 | 0.24  4 | 0.291 | 0.305 | 0.244 |
| Thlypopsis  pectoralis | Brown-flanked  Tanager | M | man | brownish-grey | grey | 0.19  4 | 0.24  1 | 0.281 | 0.284 | 0.241 |
| Thlypopsis  sordida | Orange-headed  Tanager | M | man | sandy-grey | grey | 0.10  4 | 0.21  4 | 0.321 | 0.361 | 0.214 |
| Thraupis sayaca | Sayaca Tanager | M | man | dull grey / bluish | grey | 0.17  7 | 0.24  2 | 0.297 | 0.284 | 0.242 |
| Charitospiza  eucosma | Coal-crested Finch | M | rum | silvery gray | grey | 0.15  7 | 0.24  3 | 0.291 | 0.309 | 0.243 |
| Conothraupis  speculigera | Black-and-white  Tanager | M | rum | grey | grey | 0.21  4 | 0.23  3 | 0.269 | 0.284 | 0.233 |
| Coryphaspiza  melanotis | Black-masked  Finch | M | rum | greyish | grey | 0.16  5 | 0.21  5 | 0.284 | 0.337 | 0.215 |
| Coryphospingus  pileatus | Pileated Finch | M | rum | greyish | grey | 0.18  7 | 0.22  2 | 0.277 | 0.314 | 0.222 |
| Creurgops  verticalis | Rufous-crested  Tanager | M | rum | leaden grey | grey | 0.19  2 | 0.23  7 | 0.279 | 0.292 | 0.237 |
| Diglossa  brunneiventris | Black-throated  Flowerpiercer | M | rum | grey | grey | 0.23  0 | 0.24  3 | 0.267 | 0.261 | 0.243 |
| Diglossa  carbonaria | Gray-bellied  Flowerpiercer | M | rum | dark grey | grey | 0.21  2 | 0.25  1 | 0.273 | 0.263 | 0.251 |
| Diuca diuca | Common Diuca-  Finch | M | rum | blue gray | grey | 0.19  3 | 0.24  4 | 0.277 | 0.286 | 0.244 |
| Hemispingus  verticalis | Black-headed  Hemispingus | M | rum | dark grey | grey | 0.17  8 | 0.22  3 | 0.281 | 0.318 | 0.223 |
| Hemispingus xanthophthalmu  s | Drab Hemispingus | M | rum | brownish-grey | grey | 0.16  6 | 0.21  0 | 0.284 | 0.340 | 0.210 |
| Idiopsar  brachyurus | Short-tailed Finch | M | rum | leaden gray | grey | 0.20  6 | 0.24  2 | 0.273 | 0.280 | 0.242 |
| Melanodera  xanthogramma | Yellow-bridled  Finch | M | rum | grey | grey | 0.21  1 | 0.20  5 | 0.293 | 0.290 | 0.205 |
| Neothraupis  fasciata | White-banded  Tanager | M | rum | grey | grey | 0.18  7 | 0.24  4 | 0.279 | 0.290 | 0.244 |

| Paroaria  coronata | Red-crested  Cardinal | M | rum | grey | grey | 0.21  5 | 0.25  0 | 0.268 | 0.267 | 0.250 |
| --- | --- | --- | --- | --- | --- | --- | --- | --- | --- | --- |
| Piezorhina  cinerea | Cinereous Finch | M | rum | pale grey | grey | 0.17  0 | 0.23  4 | 0.283 | 0.312 | 0.234 |
| Poospiza alticola | Plain-tailed  Warbling-Finch | M | rum | gray brown | grey | 0.17  9 | 0.24  1 | 0.284 | 0.296 | 0.241 |
| Poospiza caesar | Chestnut-breasted  Mountain-Finch | M | rum | gray | grey | 0.17  6 | 0.23  8 | 0.284 | 0.303 | 0.238 |
| Poospiza  hispaniolensis | Collared Warbling-  Finch | M | rum | brownish-grey | grey | 0.19  5 | 0.25  4 | 0.279 | 0.272 | 0.254 |
| Poospiza  melanoleuca | Black-capped  Warbling-Finch | M | rum | no | grey | 0.18  8 | 0.24  4 | 0.282 | 0.286 | 0.244 |
| Poospiza  nigrorufa | Black-and-rufous  Warbling-Finch | M | rum | blue-grey | grey | 0.19  6 | 0.23  4 | 0.278 | 0.291 | 0.234 |
| Poospiza  torquata | Ringed Warbling-  Finch | M | rum | grey | grey | 0.19  2 | 0.25  6 | 0.280 | 0.272 | 0.256 |
| Saltator  aurantiirostris | Golden-billed  Saltator | M | rum | grey | grey | 0.17  6 | 0.22  8 | 0.287 | 0.308 | 0.228 |
| Saltator  coerulescens | Grayish Saltator | M | rum | greyish | grey | 0.15  9 | 0.21  5 | 0.289 | 0.336 | 0.215 |
| Saltator  nigriceps | Black-cowled  Saltator | M | rum | deep gray / olive | grey | 0.20  3 | 0.24  5 | 0.275 | 0.277 | 0.245 |
| Saltator similis | Green-winged  Saltator | M | rum | grey | grey | 0.17  1 | 0.20  8 | 0.293 | 0.328 | 0.208 |
| Schistochlamys  melanopis | Black-faced  Tanager | M | rum | grey | grey | 0.20  7 | 0.25  1 | 0.274 | 0.267 | 0.251 |
| Sporophila  albogularis | White-throated  Seedeater | M | rum | dark greyish | grey | 0.19  9 | 0.25  0 | 0.276 | 0.276 | 0.250 |
| Sporophila  plumbea | Plumbeous  Seedeater | M | rum | lead-grey | grey | 0.17  3 | 0.23  4 | 0.283 | 0.311 | 0.234 |
| Tangara inornata | Plain-colored  Tanager | M | rum | dark gray | grey | 0.20  4 | 0.25  6 | 0.274 | 0.266 | 0.256 |
| Thlypopsis  fulviceps | Fulvous-headed  Tanager | M | rum | grey | grey | 0.15  5 | 0.22  6 | 0.298 | 0.321 | 0.226 |
| Thlypopsis  inornata | Buff-bellied  Tanager | M | rum | olive-grey | grey | 0.14  6 | 0.22  9 | 0.296 | 0.329 | 0.229 |
| Thlypopsis  pectoralis | Brown-flanked  Tanager | M | rum | brownish-grey | grey | 0.17  4 | 0.21  9 | 0.283 | 0.324 | 0.219 |
| Thlypopsis  sordida | Orange-headed  Tanager | M | rum | sandy-grey | grey | 0.13  7 | 0.21  8 | 0.305 | 0.340 | 0.218 |
| Thraupis sayaca | Sayaca Tanager | M | rum | dull grey / bluish | grey | 0.19  5 | 0.24  0 | 0.295 | 0.269 | 0.240 |
| Cnemoscopus  rubrirostris | Gray-hooded Bush  Tanager | M | thr | medium grey | grey | 0.12  4 | 0.24  0 | 0.309 | 0.326 | 0.240 |
| Embernagra  platensis | Great Pampa-  Finch | M | thr | grey | grey | 0.14  5 | 0.24  8 | 0.297 | 0.310 | 0.248 |
| Phrygilus  plebejus | Ash-breasted  Sierra-Finch | M | thr | greyish-white | grey | 0.18  0 | 0.25  2 | 0.281 | 0.286 | 0.252 |
| Tangara inornata | Plain-colored  Tanager | M | thr | dark gray | grey | 0.17  6 | 0.24  9 | 0.285 | 0.290 | 0.249 |
| Thraupis  glaucocolpa | Glaucous Tanager | M | thr | smoky grey | grey | 0.16  7 | 0.25  1 | 0.290 | 0.292 | 0.251 |
| Thraupis sayaca | Sayaca Tanager | M | thr | light grey / bluish | grey | 0.16  5 | 0.25  0 | 0.289 | 0.296 | 0.250 |
| Xenospingus  concolor | Slender-billed  Finch | M | thr | paler grey | grey | 0.15  9 | 0.25  1 | 0.291 | 0.298 | 0.251 |
| Cnemoscopus  rubrirostris | Gray-hooded Bush  Tanager | M | bre | medium grey | grey | 0.12  4 | 0.14  9 | 0.344 | 0.382 | 0.149 |
| Conirostrum  bicolor | Bicolored Conebill | M | bre | pale greyish-buff | grey | 0.10  0 | 0.22  5 | 0.320 | 0.355 | 0.225 |
| Embernagra  platensis | Great Pampa-  Finch | M | bre | grey | grey | 0.10  8 | 0.24  2 | 0.313 | 0.337 | 0.242 |
| Hemispingus  verticalis | Black-headed  Hemispingus | M | bre | pale grey | grey | 0.17  6 | 0.24  8 | 0.285 | 0.290 | 0.248 |
| Idiopsar  brachyurus | Short-tailed Finch | M | bre | leaden gray | grey | 0.17  1 | 0.24  9 | 0.283 | 0.296 | 0.249 |
| Incaspiza laeta | Buff-bridled Inca-  Finch | M | bre | grey | grey | 0.14  4 | 0.24  4 | 0.297 | 0.315 | 0.244 |
| Incaspiza | Rufous-backed | M | bre | grey | grey | 0.13 | 0.24 | 0.300 | 0.322 | 0.243 |

| personata | Inca-Finch |  |  |  |  | 5 | 3 |  |  |  |
| --- | --- | --- | --- | --- | --- | --- | --- | --- | --- | --- |
| Incaspiza pulchra | Great Inca-Finch | M | bre | grey | grey | 0.16  8 | 0.25  4 | 0.286 | 0.292 | 0.254 |
| Neothraupis  fasciata | White-banded  Tanager | M | bre | grey | grey | 0.14  8 | 0.25  1 | 0.294 | 0.308 | 0.251 |
| Phrygilus  plebejus | Ash-breasted  Sierra-Finch | M | bre | greyish-white | grey | 0.15  5 | 0.24  5 | 0.290 | 0.310 | 0.245 |
| Piezorhina  cinerea | Cinereous Finch | M | bre | pale grey | grey | 0.13  3 | 0.24  4 | 0.301 | 0.322 | 0.244 |
| Poospiza  hypochondria | Rufous-sided  Warbling-Finch | M | bre | grey | grey | 0.11  3 | 0.23  1 | 0.310 | 0.346 | 0.231 |
| Saltator atriceps | Black-headed  Saltator | M | bre | gray | grey | 0.16  2 | 0.25  4 | 0.288 | 0.295 | 0.254 |
| Saltator  atripennis | Black-winged  Saltator | M | bre | dove-grey | grey | 0.15  6 | 0.25  4 | 0.290 | 0.299 | 0.254 |
| Saltator  aurantiirostris | Golden-billed  Saltator | M | bre | grey | grey | 0.08  1 | 0.20  0 | 0.325 | 0.394 | 0.200 |
| Saltator  coerulescens | Grayish Saltator | M | bre | greyish | grey | 0.08  6 | 0.20  5 | 0.321 | 0.388 | 0.205 |
| Saltator  maximus | Buff-throated  Saltator | M | bre | grayish | grey | 0.09  4 | 0.21  5 | 0.320 | 0.371 | 0.215 |
| Saltator  nigriceps | Black-cowled  Saltator | M | bre | gray | grey | 0.19  1 | 0.25  2 | 0.278 | 0.279 | 0.252 |
| Sporophila  plumbea | Plumbeous  Seedeater | M | bre | pale grey | grey | 0.15  7 | 0.24  9 | 0.288 | 0.306 | 0.249 |
| Thlypopsis  fulviceps | Fulvous-headed  Tanager | M | bre | grey | grey | 0.13  1 | 0.24  6 | 0.303 | 0.320 | 0.246 |
| Thraupis sayaca | Sayaca Tanager | M | bre | light grey / bluish | grey | 0.17  0 | 0.25  1 | 0.295 | 0.285 | 0.251 |
| Xenospingus  concolor | Slender-billed  Finch | M | bre | paler grey | grey | 0.14  4 | 0.24  8 | 0.296 | 0.312 | 0.248 |
| Diglossa  carbonaria | Gray-bellied  Flowerpiercer | M | bel | grey | grey | 0.18  1 | 0.25  0 | 0.283 | 0.285 | 0.250 |
| Idiopsar  brachyurus | Short-tailed Finch | M | bel | leaden gray | grey | 0.16  4 | 0.24  4 | 0.289 | 0.303 | 0.244 |
| Neothraupis  fasciata | White-banded  Tanager | M | bel | grey | grey | 0.13  9 | 0.24  6 | 0.300 | 0.315 | 0.246 |
| Phrygilus  plebejus | Ash-breasted  Sierra-Finch | M | bel | greyish-white | grey | 0.15  4 | 0.24  9 | 0.292 | 0.305 | 0.249 |
| Piezorhina  cinerea | Cinereous Finch | M | bel | pale grey / white | grey | 0.13  5 | 0.24  8 | 0.300 | 0.317 | 0.248 |
| Saltator atriceps | Black-headed  Saltator | M | bel | gray | grey | 0.16  3 | 0.25  0 | 0.288 | 0.298 | 0.250 |
| Saltator  atripennis | Black-winged  Saltator | M | bel | grey-white | grey | 0.14  5 | 0.25  4 | 0.295 | 0.306 | 0.254 |
| Saltator  maximus | Buff-throated  Saltator | M | bel | grayish | grey | 0.11  8 | 0.22  5 | 0.309 | 0.348 | 0.225 |
| Saltator  nigriceps | Black-cowled  Saltator | M | bel | gray / buffy | grey | 0.12  6 | 0.23  2 | 0.306 | 0.336 | 0.232 |
| Schistochlamys  melanopis | Black-faced  Tanager | M | bel | grey | grey | 0.17  3 | 0.25  7 | 0.283 | 0.286 | 0.257 |
| Sporophila  plumbea | Plumbeous  Seedeater | M | bel | lead-grey | grey | 0.14  8 | 0.25  8 | 0.294 | 0.300 | 0.258 |
| Thraupis  glaucocolpa | Glaucous Tanager | M | bel | white | grey | 0.14  6 | 0.25  2 | 0.307 | 0.294 | 0.252 |
| Urothraupis  stolzmanni | Black-backed Bush  Tanager | M | bel | gray | grey | 0.16  8 | 0.24  6 | 0.290 | 0.296 | 0.246 |
| Xenospingus  concolor | Slender-billed  Finch | M | bel | paler grey | grey | 0.15  0 | 0.24  6 | 0.296 | 0.308 | 0.246 |
| Acanthidops  bairdii | Peg-billed Finch | F | cro | brownish-olive | grey | 0.13  0 | 0.20  9 | 0.313 | 0.347 | 0.209 |
| Camarhynchus  parvulus | Small Tree-Finch | F | cro | greyish-brown | grey | 0.14  6 | 0.22  5 | 0.299 | 0.330 | 0.225 |
| Cnemoscopus  rubrirostris | Gray-hooded Bush  Tanager | F | cro | medium grey | grey | 0.13  2 | 0.24  6 | 0.311 | 0.312 | 0.246 |
| Coereba flaveola | Bananaquit | F | cro | dark grey | grey | 0.18  6 | 0.24  2 | 0.289 | 0.282 | 0.242 |
| Creurgops  verticalis | Rufous-crested  Tanager | F | cro | leaden grey | grey | 0.17  3 | 0.24  4 | 0.290 | 0.293 | 0.244 |

| Diuca diuca | Common Diuca-  Finch | F | cro | dark gray | grey | 0.12  1 | 0.20  6 | 0.308 | 0.364 | 0.206 |
| --- | --- | --- | --- | --- | --- | --- | --- | --- | --- | --- |
| Hemispingus  melanotis | Black-eared  Hemispingus | F | cro | grey | grey | 0.19  3 | 0.21  7 | 0.294 | 0.296 | 0.217 |
| Heterospingus  xanthopygius | Scarlet-browed  Tanager | F | cro | dark leaden grey | grey | 0.16  3 | 0.23  5 | 0.298 | 0.304 | 0.235 |
| Incaspiza  personata | Rufous-backed  Inca-Finch | F | cro | grey | grey | 0.14  4 | 0.22  4 | 0.301 | 0.331 | 0.224 |
| Lophospingus  griseocristatus | Gray-crested Finch | F | cro | gray | grey | 0.16  0 | 0.24  7 | 0.292 | 0.301 | 0.247 |
| Neothraupis  fasciata | White-banded  Tanager | F | cro | grey | grey | 0.13  0 | 0.22  4 | 0.306 | 0.339 | 0.224 |
| Phrygilus  atriceps | Black-hooded  Sierra-Finch | F | cro | black | grey | 0.16  2 | 0.23  2 | 0.294 | 0.312 | 0.232 |
| Piezorhina  cinerea | Cinereous Finch | F | cro | pale grey | grey | 0.14  0 | 0.23  5 | 0.301 | 0.325 | 0.235 |
| Poospiza alticola | Plain-tailed  Warbling-Finch | F | cro | gray brown | grey | 0.17  8 | 0.23  4 | 0.284 | 0.303 | 0.234 |
| Poospiza caesar | Chestnut-breasted  Mountain-Finch | F | cro | gray | grey | 0.16  2 | 0.23  4 | 0.296 | 0.308 | 0.234 |
| Poospiza  thoracica | Bay-chested  Warbling-Finch | F | cro | grey | grey | 0.11  9 | 0.23  4 | 0.312 | 0.335 | 0.234 |
| Ramphocelus  passerinii | Scarlet-rumped  Tanager | F | cro | grey | grey | 0.12  7 | 0.20  9 | 0.304 | 0.360 | 0.209 |
| Saltator  coerulescens | Grayish Saltator | F | cro | greyish | grey | 0.15  8 | 0.22  6 | 0.293 | 0.322 | 0.226 |
| Tachyphonus  luctuosus | White-shouldered  Tanager | F | cro | grey | grey | 0.13  8 | 0.20  6 | 0.325 | 0.332 | 0.206 |
| Tachyphonus  phoenicius | Red-shouldered  Tanager | F | cro | brownish-grey | grey | 0.15  3 | 0.23  6 | 0.299 | 0.312 | 0.236 |
| Tachyphonus  rufiventer | Yellow-crested  Tanager | F | cro | grey | grey | 0.11  5 | 0.16  5 | 0.335 | 0.385 | 0.165 |
| Tangara inornata | Plain-colored  Tanager | F | cro | dark gray | grey | 0.16  4 | 0.24  2 | 0.292 | 0.302 | 0.242 |
| Thraupis  glaucocolpa | Glaucous Tanager | F | cro | smoky grey /  greenish | grey | 0.17  7 | 0.24  1 | 0.291 | 0.291 | 0.241 |
| Thraupis sayaca | Sayaca Tanager | F | cro | dull grey / bluish | grey | 0.16  5 | 0.23  7 | 0.297 | 0.301 | 0.237 |
| Tiaris canorus | Cuban Grassquit | F | cro | grey / greenish | grey | 0.14  2 | 0.21  7 | 0.294 | 0.346 | 0.217 |
| Acanthidops  bairdii | Peg-billed Finch | F | nap | brownish-olive | grey | 0.14  7 | 0.21  4 | 0.303 | 0.336 | 0.214 |
| Camarhynchus  parvulus | Small Tree-Finch | F | nap | greyish-brown | grey | 0.15  4 | 0.22  5 | 0.295 | 0.327 | 0.225 |
| Coereba flaveola | Bananaquit | F | nap | dark grey | grey | 0.18  0 | 0.24  2 | 0.291 | 0.287 | 0.242 |
| Compsospiza  garleppi | Cochabamba  Mountain-Finch | F | nap | leaden grey | grey | 0.16  5 | 0.25  2 | 0.293 | 0.290 | 0.252 |
| Conirostrum  albifrons | Capped Conebill | F | nap | blue-tinged  greyish | grey | 0.20  7 | 0.25  0 | 0.268 | 0.275 | 0.250 |
| Coryphospingus  pileatus | Pileated Finch | F | nap | greyish-brown | grey | 0.16  2 | 0.22  0 | 0.291 | 0.327 | 0.220 |
| Creurgops  verticalis | Rufous-crested  Tanager | F | nap | leaden grey | grey | 0.18  7 | 0.24  2 | 0.282 | 0.289 | 0.242 |
| Diuca diuca | Common Diuca-  Finch | F | nap | dark gray | grey | 0.16  8 | 0.22  0 | 0.290 | 0.323 | 0.220 |
| Hemispingus  melanotis | Black-eared  Hemispingus | F | nap | grey | grey | 0.20  5 | 0.22  9 | 0.279 | 0.287 | 0.229 |
| Heterospingus  xanthopygius | Scarlet-browed  Tanager | F | nap | dark leaden grey | grey | 0.17  5 | 0.23  0 | 0.289 | 0.306 | 0.230 |
| Lophospingus  griseocristatus | Gray-crested Finch | F | nap | gray | grey | 0.17  6 | 0.24  1 | 0.285 | 0.299 | 0.241 |
| Neothraupis  fasciata | White-banded  Tanager | F | nap | grey | grey | 0.14  4 | 0.22  6 | 0.298 | 0.332 | 0.226 |
| Piezorhina  cinerea | Cinereous Finch | F | nap | pale grey | grey | 0.14  9 | 0.24  4 | 0.296 | 0.312 | 0.244 |
| Poospiza alticola | Plain-tailed  Warbling-Finch | F | nap | gray brown | grey | 0.17  0 | 0.22  5 | 0.284 | 0.321 | 0.225 |
| Poospiza caesar | Chestnut-breasted | F | nap | gray | grey | 0.17 | 0.23 | 0.287 | 0.309 | 0.235 |

|  | Mountain-Finch |  |  |  |  | 0 | 5 |  |  |  |
| --- | --- | --- | --- | --- | --- | --- | --- | --- | --- | --- |
| Poospiza  melanoleuca | Black-capped  Warbling-Finch | F | nap | blue-grey | grey | 0.20  9 | 0.24  1 | 0.274 | 0.276 | 0.241 |
| Poospiza  thoracica | Bay-chested  Warbling-Finch | F | nap | grey | grey | 0.15  0 | 0.24  0 | 0.294 | 0.316 | 0.240 |
| Poospiza  torquata | Ringed Warbling-  Finch | F | nap | grey | grey | 0.16  5 | 0.24  2 | 0.292 | 0.301 | 0.242 |
| Saltator  coerulescens | Grayish Saltator | F | nap | greyish | grey | 0.15  7 | 0.22  0 | 0.300 | 0.323 | 0.220 |
| Schistochlamys  melanopis | Black-faced  Tanager | F | nap | grey | grey | 0.18  4 | 0.25  8 | 0.285 | 0.273 | 0.258 |
| Tachyphonus  luctuosus | White-shouldered  Tanager | F | nap | grey | grey | 0.16  0 | 0.20  0 | 0.312 | 0.328 | 0.200 |
| Tachyphonus  phoenicius | Red-shouldered  Tanager | F | nap | brownish-grey | grey | 0.18  3 | 0.23  4 | 0.283 | 0.299 | 0.234 |
| Tangara inornata | Plain-colored  Tanager | F | nap | dark gray | grey | 0.17  7 | 0.24  2 | 0.285 | 0.296 | 0.242 |
| Thraupis  glaucocolpa | Glaucous Tanager | F | nap | smoky grey /  greenish | grey | 0.17  9 | 0.24  3 | 0.296 | 0.282 | 0.243 |
| Thraupis sayaca | Sayaca Tanager | F | nap | dull grey / bluish | grey | 0.17  1 | 0.23  2 | 0.300 | 0.297 | 0.232 |
| Acanthidops  bairdii | Peg-billed Finch | F | man | brownish-olive | grey | 0.18  8 | 0.22  9 | 0.283 | 0.300 | 0.229 |
| Camarhynchus  parvulus | Small Tree-Finch | F | man | greyish-brown | grey | 0.14  7 | 0.23  0 | 0.297 | 0.326 | 0.230 |
| Coereba flaveola | Bananaquit | F | man | dark grey | grey | 0.20  4 | 0.23  7 | 0.279 | 0.280 | 0.237 |
| Compsospiza  garleppi | Cochabamba  Mountain-Finch | F | man | leaden grey | grey | 0.16  3 | 0.25  1 | 0.295 | 0.292 | 0.251 |
| Coryphospingus  pileatus | Pileated Finch | F | man | greyish-brown | grey | 0.18  3 | 0.23  0 | 0.283 | 0.305 | 0.230 |
| Creurgops  verticalis | Rufous-crested  Tanager | F | man | leaden grey | grey | 0.20  0 | 0.24  4 | 0.277 | 0.278 | 0.244 |
| Diuca diuca | Common Diuca-  Finch | F | man | dark gray | grey | 0.16  7 | 0.22  5 | 0.288 | 0.320 | 0.225 |
| Heterospingus  xanthopygius | Scarlet-browed  Tanager | F | man | dark leaden grey | grey | 0.18  6 | 0.22  7 | 0.282 | 0.305 | 0.227 |
| Lophospingus  griseocristatus | Gray-crested Finch | F | man | gray / olive | grey | 0.16  8 | 0.24  2 | 0.287 | 0.303 | 0.242 |
| Neothraupis  fasciata | White-banded  Tanager | F | man | grey | grey | 0.18  1 | 0.23  8 | 0.284 | 0.298 | 0.238 |
| Paroaria  coronata | Red-crested  Cardinal | F | man | grey | grey | 0.19  0 | 0.24  4 | 0.278 | 0.288 | 0.244 |
| Piezorhina  cinerea | Cinereous Finch | F | man | pale grey | grey | 0.19  0 | 0.24  7 | 0.280 | 0.284 | 0.247 |
| Poospiza alticola | Plain-tailed  Warbling-Finch | F | man | gray brown | grey | 0.14  7 | 0.22  5 | 0.298 | 0.330 | 0.225 |
| Poospiza caesar | Chestnut-breasted  Mountain-Finch | F | man | gray | grey | 0.18  5 | 0.24  2 | 0.282 | 0.291 | 0.242 |
| Poospiza  melanoleuca | Black-capped  Warbling-Finch | F | man | blue-grey | grey | 0.19  4 | 0.25  4 | 0.280 | 0.272 | 0.254 |
| Poospiza  thoracica | Bay-chested  Warbling-Finch | F | man | grey | grey | 0.15  2 | 0.22  9 | 0.297 | 0.323 | 0.229 |
| Poospiza  torquata | Ringed Warbling-  Finch | F | man | grey | grey | 0.13  7 | 0.23  6 | 0.302 | 0.325 | 0.236 |
| Saltator  aurantiirostris | Golden-billed  Saltator | F | man | grey | grey | 0.16  4 | 0.23  3 | 0.293 | 0.309 | 0.233 |
| Saltator  coerulescens | Grayish Saltator | F | man | greyish | grey | 0.15  1 | 0.21  2 | 0.300 | 0.337 | 0.212 |
| Schistochlamys  melanopis | Black-faced  Tanager | F | man | grey | grey | 0.19  9 | 0.26  0 | 0.279 | 0.262 | 0.260 |
| Tangara inornata | Plain-colored  Tanager | F | man | dark gray | grey | 0.18  5 | 0.24  6 | 0.281 | 0.288 | 0.246 |
| Thlypopsis  sordida | Orange-headed  Tanager | F | man | sandy-grey | grey | 0.10  1 | 0.21  7 | 0.323 | 0.359 | 0.217 |
| Thraupis sayaca | Sayaca Tanager | F | man | dull grey / bluish | grey | 0.17  9 | 0.23  5 | 0.297 | 0.289 | 0.235 |
| Charitospiza  eucosma | Coal-crested Finch | F | rum | gray-gold | grey | 0.19  7 | 0.24  5 | 0.273 | 0.284 | 0.245 |

| Compsospiza  garleppi | Cochabamba  Mountain-Finch | F | rum | leaden grey | grey | 0.18  3 | 0.25  1 | 0.285 | 0.281 | 0.251 |
| --- | --- | --- | --- | --- | --- | --- | --- | --- | --- | --- |
| Creurgops  verticalis | Rufous-crested  Tanager | F | rum | leaden grey | grey | 0.15  9 | 0.24  3 | 0.293 | 0.305 | 0.243 |
| Diglossa  brunneiventris | Black-throated  Flowerpiercer | F | rum | grey | grey | 0.21  1 | 0.24  7 | 0.273 | 0.268 | 0.247 |
| Diuca diuca | Common Diuca-  Finch | F | rum | blue gray | grey | 0.15  9 | 0.22  4 | 0.292 | 0.324 | 0.224 |
| Lophospingus  griseocristatus | Gray-crested Finch | F | rum | blue gray | grey | 0.19  1 | 0.24  3 | 0.278 | 0.287 | 0.243 |
| Neothraupis  fasciata | White-banded  Tanager | F | rum | grey | grey | 0.17  5 | 0.23  5 | 0.287 | 0.303 | 0.235 |
| Paroaria  coronata | Red-crested  Cardinal | F | rum | grey | grey | 0.21  0 | 0.24  6 | 0.270 | 0.274 | 0.246 |
| Piezorhina  cinerea | Cinereous Finch | F | rum | pale grey | grey | 0.19  4 | 0.24  7 | 0.277 | 0.281 | 0.247 |
| Poospiza alticola | Plain-tailed  Warbling-Finch | F | rum | gray brown | grey | 0.14  7 | 0.22  3 | 0.299 | 0.331 | 0.223 |
| Poospiza caesar | Chestnut-breasted  Mountain-Finch | F | rum | gray | grey | 0.18  4 | 0.24  5 | 0.282 | 0.289 | 0.245 |
| Poospiza  melanoleuca | Black-capped  Warbling-Finch | F | rum | blue-grey | grey | 0.21  1 | 0.25  3 | 0.272 | 0.264 | 0.253 |
| Poospiza  torquata | Ringed Warbling-  Finch | F | rum | grey | grey | 0.16  7 | 0.24  3 | 0.288 | 0.302 | 0.243 |
| Saltator  aurantiirostris | Golden-billed  Saltator | F | rum | grey | grey | 0.17  0 | 0.23  6 | 0.291 | 0.302 | 0.236 |
| Saltator  coerulescens | Grayish Saltator | F | rum | greyish | grey | 0.13  6 | 0.21  7 | 0.306 | 0.341 | 0.217 |
| Saltator similis | Green-winged  Saltator | F | rum | grey | grey | 0.16  0 | 0.21  2 | 0.300 | 0.328 | 0.212 |
| Schistochlamys  melanopis | Black-faced  Tanager | F | rum | grey | grey | 0.20  6 | 0.25  8 | 0.275 | 0.261 | 0.258 |
| Tangara inornata | Plain-colored  Tanager | F | rum | dark gray | grey | 0.18  7 | 0.24  4 | 0.279 | 0.290 | 0.244 |
| Thlypopsis  sordida | Orange-headed  Tanager | F | rum | sandy-grey | grey | 0.11  2 | 0.21  3 | 0.320 | 0.355 | 0.213 |
| Thraupis sayaca | Sayaca Tanager | F | rum | dull grey / bluish | grey | 0.19  8 | 0.23  2 | 0.302 | 0.269 | 0.232 |
| Cnemoscopus  rubrirostris | Gray-hooded Bush  Tanager | F | thr | medium grey | grey | 0.12  9 | 0.24  5 | 0.307 | 0.319 | 0.245 |
| Conirostrum  albifrons | Capped Conebill | F | thr | pale grey / bluish | grey | 0.16  2 | 0.24  3 | 0.289 | 0.306 | 0.243 |
| Dacnis cayana | Blue Dacnis | F | thr | grey | grey | 0.18  2 | 0.23  9 | 0.297 | 0.281 | 0.239 |
| Dacnis venusta | Scarlet-thighed  Dacnis | F | thr | grayish | grey | 0.16  9 | 0.23  3 | 0.313 | 0.285 | 0.233 |
| Heterospingus  xanthopygius | Scarlet-browed  Tanager | F | thr | paler grey | grey | 0.15  2 | 0.23  5 | 0.296 | 0.317 | 0.235 |
| Lanio aurantius | Black-throated  Shrike-Tanager | F | thr | grey | grey | 0.11  9 | 0.21  7 | 0.308 | 0.356 | 0.217 |
| Lophospingus  griseocristatus | Gray-crested Finch | F | thr | paler gray | grey | 0.16  1 | 0.25  2 | 0.289 | 0.298 | 0.252 |
| Loxigilla noctis | Lesser Antillean  Bullfinch | F | thr | brown / greyer | grey | 0.11  9 | 0.21  5 | 0.309 | 0.357 | 0.215 |
| Loxipasser  anoxanthus | Yellow-shouldered  Grassquit | F | thr | grey / green | grey | 0.13  3 | 0.21  6 | 0.303 | 0.347 | 0.216 |
| Phrygilus  fruticeti | Mourning Sierra-  Finch | F | thr | grey | grey | 0.17  1 | 0.24  3 | 0.288 | 0.298 | 0.243 |
| Ramphocelus  passerinii | Scarlet-rumped  Tanager | F | thr | grey | grey | 0.11  4 | 0.20  3 | 0.313 | 0.370 | 0.203 |
| Tangara inornata | Plain-colored  Tanager | F | thr | dark gray | grey | 0.15  5 | 0.24  8 | 0.295 | 0.302 | 0.248 |
| Thraupis  glaucocolpa | Glaucous Tanager | F | thr | smoky grey | grey | 0.17  0 | 0.25  1 | 0.286 | 0.292 | 0.251 |
| Thraupis sayaca | Sayaca Tanager | F | thr | light grey / bluish | grey | 0.14  8 | 0.24  8 | 0.297 | 0.307 | 0.248 |
| Xenospingus  concolor | Slender-billed  Finch | F | thr | paler grey | grey | 0.15  0 | 0.25  5 | 0.293 | 0.301 | 0.255 |
| Cnemoscopus | Gray-hooded Bush | F | bre | medium grey | grey | 0.10 | 0.16 | 0.345 | 0.388 | 0.163 |

| rubrirostris | Tanager |  |  |  |  | 5 | 3 |  |  |  |
| --- | --- | --- | --- | --- | --- | --- | --- | --- | --- | --- |
| Conirostrum  albifrons | Capped Conebill | F | bre | pale grey / bluish | grey | 0.16  4 | 0.24  7 | 0.286 | 0.303 | 0.247 |
| Conirostrum  bicolor | Bicolored Conebill | F | bre | pale greyish-buff | grey | 0.10  0 | 0.13  3 | 0.359 | 0.409 | 0.133 |
| Heterospingus  xanthopygius | Scarlet-browed  Tanager | F | bre | paler grey | grey | 0.18  3 | 0.23  6 | 0.283 | 0.298 | 0.236 |
| Incaspiza laeta | Buff-bridled Inca-  Finch | F | bre | grey | grey | 0.12  0 | 0.23  5 | 0.306 | 0.339 | 0.235 |
| Incaspiza  personata | Rufous-backed  Inca-Finch | F | bre | grey | grey | 0.11  0 | 0.23  1 | 0.311 | 0.348 | 0.231 |
| Incaspiza pulchra | Great Inca-Finch | F | bre | grey | grey | 0.14  6 | 0.24  6 | 0.295 | 0.314 | 0.246 |
| Lophospingus  griseocristatus | Gray-crested Finch | F | bre | paler gray | grey | 0.15  8 | 0.25  0 | 0.289 | 0.303 | 0.250 |
| Lophospingus  pusillus | Black-crested  Finch | F | bre | greyish | grey | 0.14  6 | 0.24  0 | 0.295 | 0.319 | 0.240 |
| Loxipasser  anoxanthus | Yellow-shouldered  Grassquit | F | bre | grey / green | grey | 0.13  4 | 0.21  5 | 0.308 | 0.343 | 0.215 |
| Neothraupis  fasciata | White-banded  Tanager | F | bre | grey | grey | 0.14  6 | 0.24  7 | 0.295 | 0.313 | 0.247 |
| Phrygilus  fruticeti | Mourning Sierra-  Finch | F | bre | white | grey | 0.16  3 | 0.23  8 | 0.288 | 0.310 | 0.238 |
| Piezorhina  cinerea | Cinereous Finch | F | bre | pale grey | grey | 0.12  5 | 0.23  4 | 0.304 | 0.337 | 0.234 |
| Poospiza  hypochondria | Rufous-sided  Warbling-Finch | F | bre | grey | grey | 0.12  7 | 0.23  9 | 0.303 | 0.331 | 0.239 |
| Saltator atriceps | Black-headed  Saltator | F | bre | gray | grey | 0.16  6 | 0.24  8 | 0.288 | 0.298 | 0.248 |
| Saltator  atripennis | Black-winged  Saltator | F | bre | dove-grey | grey | 0.16  5 | 0.25  3 | 0.288 | 0.294 | 0.253 |
| Saltator  aurantiirostris | Golden-billed  Saltator | F | bre | grey | grey | 0.07  9 | 0.19  6 | 0.326 | 0.398 | 0.196 |
| Saltator  coerulescens | Grayish Saltator | F | bre | greyish | grey | 0.10  0 | 0.20  3 | 0.320 | 0.377 | 0.203 |
| Saltator  maximus | Buff-throated  Saltator | F | bre | grayish / buff | grey | 0.10  2 | 0.21  7 | 0.315 | 0.366 | 0.217 |
| Sicalis  taczanowskii | Sulphur-throated  Finch | F | bre | pale greyish-  brown | grey | 0.15  9 | 0.23  9 | 0.293 | 0.308 | 0.239 |
| Tangara  viridicollis | Silvery Tanager | F | bre | grey | grey | 0.11  1 | 0.21  3 | 0.339 | 0.337 | 0.213 |
| Thraupis sayaca | Sayaca Tanager | F | bre | light grey / bluish | grey | 0.15  7 | 0.25  0 | 0.298 | 0.294 | 0.250 |
| Tiaris canorus | Cuban Grassquit | F | bre | grey | grey | 0.11  7 | 0.23  1 | 0.310 | 0.342 | 0.231 |
| Xenospingus  concolor | Slender-billed  Finch | F | bre | paler grey | grey | 0.15  1 | 0.25  3 | 0.292 | 0.304 | 0.253 |
| Heterospingus  xanthopygius | Scarlet-browed  Tanager | F | bel | paler grey | grey | 0.15  7 | 0.22  5 | 0.295 | 0.323 | 0.225 |
| Lophospingus  griseocristatus | Gray-crested Finch | F | bel | paler gray | grey | 0.14  9 | 0.25  2 | 0.293 | 0.306 | 0.252 |
| Lophospingus  pusillus | Black-crested  Finch | F | bel | greyish | grey | 0.15  2 | 0.25  0 | 0.293 | 0.305 | 0.250 |
| Loxipasser  anoxanthus | Yellow-shouldered  Grassquit | F | bel | greenish / yellow | grey | 0.13  7 | 0.22  4 | 0.305 | 0.334 | 0.224 |
| Neothraupis  fasciata | White-banded  Tanager | F | bel | grey | grey | 0.14  1 | 0.24  1 | 0.300 | 0.318 | 0.241 |
| Piezorhina  cinerea | Cinereous Finch | F | bel | pale grey / white | grey | 0.12  4 | 0.24  3 | 0.304 | 0.328 | 0.243 |
| Saltator atriceps | Black-headed  Saltator | F | bel | gray | grey | 0.15  7 | 0.24  6 | 0.292 | 0.305 | 0.246 |
| Saltator  atripennis | Black-winged  Saltator | F | bel | grey-white | grey | 0.17  7 | 0.25  0 | 0.283 | 0.289 | 0.250 |
| Saltator  maximus | Buff-throated  Saltator | F | bel | grayish | grey | 0.09  9 | 0.20  6 | 0.317 | 0.378 | 0.206 |
| Schistochlamys  melanopis | Black-faced  Tanager | F | bel | grey | grey | 0.16  3 | 0.25  4 | 0.288 | 0.295 | 0.254 |
| Sicalis  taczanowskii | Sulphur-throated  Finch | F | bel | pale greyish-  brown | grey | 0.18  5 | 0.25  4 | 0.278 | 0.283 | 0.254 |

| Tangara  viridicollis | Silvery Tanager | F | bel | grey | grey | 0.13  8 | 0.25  1 | 0.311 | 0.300 | 0.251 |
| --- | --- | --- | --- | --- | --- | --- | --- | --- | --- | --- |
| Tiaris canorus | Cuban Grassquit | F | bel | olive-grey | grey | 0.10  7 | 0.22  5 | 0.316 | 0.353 | 0.225 |
| Urothraupis  stolzmanni | Black-backed Bush  Tanager | F | bel | gray | grey | 0.17  0 | 0.24  8 | 0.288 | 0.294 | 0.248 |
| Xenospingus  concolor | Slender-billed  Finch | F | bel | paler grey | grey | 0.14  8 | 0.25  1 | 0.295 | 0.306 | 0.251 |
| Acanthidops  bairdii | Peg-billed Finch | M | cro | dark grey | slate | 0.20  2 | 0.24  3 | 0.278 | 0.277 | 0.243 |
| Bangsia arcaei | Blue-and-gold  Tanager | M | cro | bright blue | slate | 0.34  6 | 0.25  8 | 0.216 | 0.181 | 0.258 |
| Catamenia analis | Band-tailed  Seedeater | M | cro | grey | slate | 0.19  0 | 0.26  5 | 0.284 | 0.261 | 0.265 |
| Catamenia  homochroa | Paramo Seedeater | M | cro | dark slate gray | slate | 0.17  2 | 0.24  3 | 0.294 | 0.291 | 0.243 |
| Catamenia  inornata | Plain-colored  Seedeater | M | cro | grey | slate | 0.14  0 | 0.24  6 | 0.303 | 0.311 | 0.246 |
| Conirostrum  cinereum | Cinereous Conebill | M | cro | slate-gray | slate | 0.13  6 | 0.23  4 | 0.310 | 0.320 | 0.234 |
| Conirostrum  rufum | Rufous-browed  Conebill | M | cro | plumbeous grey | slate | 0.13  9 | 0.24  9 | 0.311 | 0.302 | 0.249 |
| Conirostrum  speciosum | Chestnut-vented  Conebill | M | cro | dark greyish-blue | slate | 0.21  1 | 0.29  5 | 0.276 | 0.218 | 0.295 |
| Delothraupis  castaneoventris | Chestnut-bellied  Mountain-Tanager | M | cro | silvery sky blue | slate | 0.29  5 | 0.29  3 | 0.227 | 0.185 | 0.293 |
| Diglossa baritula | Cinnamon-bellied  Flowerpiercer | M | cro | slate-blackish | slate | 0.23  9 | 0.26  2 | 0.265 | 0.233 | 0.262 |
| Diglossa  caerulescens | Bluish  Flowerpiercer | M | cro | dull bluish-grey | slate | 0.23  3 | 0.26  5 | 0.266 | 0.236 | 0.265 |
| Diglossa duidae | Scaled  Flowerpiercer | M | cro | slaty black | slate | 0.21  0 | 0.24  9 | 0.284 | 0.258 | 0.249 |
| Diglossa  plumbea | Slaty  Flowerpiercer | M | cro | blackish-grey /  blue | slate | 0.22  2 | 0.25  5 | 0.271 | 0.252 | 0.255 |
| Eucometis  penicillata | Gray-headed  Tanager | M | cro | grey | slate | 0.06  4 | 0.23  3 | 0.336 | 0.367 | 0.233 |
| Euneornis  campestris | Orangequit | M | cro | grey-blue | slate | 0.23  9 | 0.29  0 | 0.264 | 0.207 | 0.290 |
| Haplospiza  unicolor | Uniform Finch | M | cro | blue-grey | slate | 0.13  7 | 0.27  0 | 0.308 | 0.284 | 0.270 |
| Oreomanes  fraseri | Giant Conebill | M | cro | plumbeous | slate | 0.15  4 | 0.24  5 | 0.299 | 0.302 | 0.245 |
| Phrygilus  alaudinus | Band-tailed Sierra-  Finch | M | cro | grey | slate | 0.19  6 | 0.24  2 | 0.279 | 0.283 | 0.242 |
| Phrygilus  erythronotus | White-throated  Sierra-Finch | M | cro | blue-grey | slate | 0.18  0 | 0.24  9 | 0.284 | 0.287 | 0.249 |
| Phrygilus gayi | Gray-hooded  Sierra-Finch | M | cro | bluish-grey | slate | 0.25  0 | 0.26  3 | 0.256 | 0.231 | 0.263 |
| Phrygilus  patagonicus | Patagonian Sierra-  Finch | M | cro | dark blue gray | slate | 0.25  8 | 0.26  5 | 0.254 | 0.223 | 0.265 |
| Phrygilus  punensis | Peruvian Sierra-  Finch | M | cro | grey | slate | 0.20  3 | 0.25  4 | 0.277 | 0.265 | 0.254 |
| Phrygilus  unicolor | Plumbeous Sierra-  Finch | M | cro | lead-grey | slate | 0.18  4 | 0.25  2 | 0.285 | 0.279 | 0.252 |
| Poospiza cinerea | Cinereous  Warbling-Finch | M | cro | grey | slate | 0.14  3 | 0.25  3 | 0.303 | 0.300 | 0.253 |
| Poospiza  erythrophrys | Rusty-browed  Warbling-Finch | M | cro | blue-grey | slate | 0.13  1 | 0.24  6 | 0.308 | 0.316 | 0.246 |
| Poospiza whitii | Black-and- chestnut  Warbling-Finch | M | cro | slate-grey | slate | 0.18  5 | 0.24  8 | 0.286 | 0.280 | 0.248 |
| Saltator  fuliginosus | Black-throated  Grosbeak | M | cro | deep slate-blue | slate | 0.22  4 | 0.25  7 | 0.273 | 0.246 | 0.257 |
| Saltator grossus | Slate-colored  Grosbeak | M | cro | slaty blue | slate | 0.22  7 | 0.25  2 | 0.269 | 0.253 | 0.252 |
| Sporophila  castaneiventris | Chestnut-bellied  Seedeater | M | cro | blue-grey | slate | 0.18  4 | 0.26  0 | 0.286 | 0.270 | 0.260 |
| Sporophila  intermedia | Gray Seedeater | M | cro | medium-grey /  blue | slate | 0.17  5 | 0.25  0 | 0.287 | 0.288 | 0.250 |

| Thraupis  episcopus | Blue-gray Tanager | M | cro | pale grey / blue  wash | slate | 0.18  8 | 0.25  4 | 0.295 | 0.263 | 0.254 |
| --- | --- | --- | --- | --- | --- | --- | --- | --- | --- | --- |
| Xenospingus  concolor | Slender-billed  Finch | M | cro | slate-grey | slate | 0.19  4 | 0.25  4 | 0.279 | 0.273 | 0.254 |
| Acanthidops  bairdii | Peg-billed Finch | M | nap | dark grey | slate | 0.20  6 | 0.24  4 | 0.274 | 0.276 | 0.244 |
| Bangsia arcaei | Blue-and-gold  Tanager | M | nap | dark blue | slate | 0.33  7 | 0.25  2 | 0.220 | 0.191 | 0.252 |
| Catamenia analis | Band-tailed  Seedeater | M | nap | grey | slate | 0.20  8 | 0.26  5 | 0.274 | 0.253 | 0.265 |
| Catamenia  homochroa | Paramo Seedeater | M | nap | dark slate gray | slate | 0.18  7 | 0.24  5 | 0.284 | 0.285 | 0.245 |
| Catamenia  inornata | Plain-colored  Seedeater | M | nap | grey | slate | 0.14  7 | 0.24  6 | 0.298 | 0.309 | 0.246 |
| Conirostrum  cinereum | Cinereous Conebill | M | nap | slate-gray | slate | 0.14  5 | 0.24  0 | 0.303 | 0.311 | 0.240 |
| Conirostrum  rufum | Rufous-browed  Conebill | M | nap | plumbeous grey | slate | 0.18  0 | 0.24  9 | 0.288 | 0.284 | 0.249 |
| Conirostrum  sitticolor | Blue-backed  Conebill | M | nap | blue | slate | 0.23  3 | 0.25  6 | 0.273 | 0.239 | 0.256 |
| Conirostrum  speciosum | Chestnut-vented  Conebill | M | nap | dark greyish-blue | slate | 0.23  0 | 0.27  9 | 0.265 | 0.226 | 0.279 |
| Delothraupis  castaneoventris | Chestnut-bellied  Mountain-Tanager | M | nap | dark blue | slate | 0.28  3 | 0.28  9 | 0.233 | 0.196 | 0.289 |
| Diglossa baritula | Cinnamon-bellied  Flowerpiercer | M | nap | slate-blackish | slate | 0.25  3 | 0.26  1 | 0.258 | 0.228 | 0.261 |
| Diglossa  caerulescens | Bluish  Flowerpiercer | M | nap | dull bluish-grey | slate | 0.23  4 | 0.26  5 | 0.265 | 0.237 | 0.265 |
| Diglossa duidae | Scaled  Flowerpiercer | M | nap | slaty black | slate | 0.22  8 | 0.24  5 | 0.267 | 0.260 | 0.245 |
| Diglossa  plumbea | Slaty  Flowerpiercer | M | nap | blackish-grey /  blue | slate | 0.21  7 | 0.24  8 | 0.272 | 0.263 | 0.248 |
| Eucometis  penicillata | Gray-headed  Tanager | M | nap | grey | slate | 0.05  8 | 0.23  0 | 0.341 | 0.371 | 0.230 |
| Euneornis  campestris | Orangequit | M | nap | grey-blue | slate | 0.25  2 | 0.28  8 | 0.254 | 0.205 | 0.288 |
| Haplospiza  unicolor | Uniform Finch | M | nap | blue-grey | slate | 0.16  7 | 0.26  7 | 0.294 | 0.273 | 0.267 |
| Nemosia pileata | Hooded Tanager | M | nap | light blue | slate | 0.21  4 | 0.27  1 | 0.275 | 0.240 | 0.271 |
| Oreomanes  fraseri | Giant Conebill | M | nap | plumbeous | slate | 0.18  7 | 0.24  3 | 0.283 | 0.286 | 0.243 |
| Phrygilus  alaudinus | Band-tailed Sierra-  Finch | M | nap | lead-grey | slate | 0.16  9 | 0.24  0 | 0.288 | 0.303 | 0.240 |
| Phrygilus  erythronotus | White-throated  Sierra-Finch | M | nap | dark gray | slate | 0.18  9 | 0.25  0 | 0.280 | 0.281 | 0.250 |
| Phrygilus gayi | Gray-hooded  Sierra-Finch | M | nap | bluish-grey | slate | 0.21  5 | 0.21  7 | 0.284 | 0.285 | 0.217 |
| Phrygilus  patagonicus | Patagonian Sierra-  Finch | M | nap | dark blue gray | slate | 0.24  4 | 0.24  5 | 0.263 | 0.247 | 0.245 |
| Phrygilus  unicolor | Plumbeous Sierra-  Finch | M | nap | lead-grey | slate | 0.20  0 | 0.25  4 | 0.276 | 0.270 | 0.254 |
| Poospiza cinerea | Cinereous  Warbling-Finch | M | nap | grey | slate | 0.17  4 | 0.25  1 | 0.288 | 0.287 | 0.251 |
| Poospiza whitii | Black-and- chestnut  Warbling-Finch | M | nap | slate-grey | slate | 0.20  1 | 0.24  8 | 0.278 | 0.274 | 0.248 |
| Saltator  fuliginosus | Black-throated  Grosbeak | M | nap | deep slate-blue | slate | 0.22  4 | 0.24  7 | 0.268 | 0.260 | 0.247 |
| Saltator grossus | Slate-colored  Grosbeak | M | nap | slaty blue | slate | 0.23  8 | 0.25  3 | 0.263 | 0.245 | 0.253 |
| Sporophila  castaneiventris | Chestnut-bellied  Seedeater | M | nap | blue-grey | slate | 0.19  6 | 0.26  0 | 0.280 | 0.265 | 0.260 |
| Sporophila  intermedia | Gray Seedeater | M | nap | medium-grey /  blue | slate | 0.18  4 | 0.25  2 | 0.284 | 0.280 | 0.252 |
| Thraupis  episcopus | Blue-gray Tanager | M | nap | pale grey / blue  wash | slate | 0.19  9 | 0.26  4 | 0.286 | 0.251 | 0.264 |
| Xenospingus  concolor | Slender-billed  Finch | M | nap | slate-grey | slate | 0.18  9 | 0.25  0 | 0.282 | 0.279 | 0.250 |

| Acanthidops  bairdii | Peg-billed Finch | M | man | dark grey | slate | 0.20  2 | 0.24  6 | 0.279 | 0.273 | 0.246 |
| --- | --- | --- | --- | --- | --- | --- | --- | --- | --- | --- |
| Bangsia arcaei | Blue-and-gold  Tanager | M | man | dark blue | slate | 0.32  2 | 0.25  5 | 0.227 | 0.196 | 0.255 |
| Catamblyrhynch  us diadema | Plushcap | M | man | blue gray | slate | 0.21  9 | 0.25  3 | 0.271 | 0.257 | 0.253 |
| Catamenia analis | Band-tailed  Seedeater | M | man | grey / browner | slate | 0.20  1 | 0.26  5 | 0.276 | 0.258 | 0.265 |
| Catamenia  homochroa | Paramo Seedeater | M | man | dark slate gray | slate | 0.17  7 | 0.24  3 | 0.287 | 0.293 | 0.243 |
| Catamenia  inornata | Plain-colored  Seedeater | M | man | grey | slate | 0.12  8 | 0.23  0 | 0.308 | 0.334 | 0.230 |
| Conirostrum  cinereum | Cinereous Conebill | M | man | slate-gray | slate | 0.14  4 | 0.23  6 | 0.305 | 0.315 | 0.236 |
| Conirostrum  leucogenys | White-eared  Conebill | M | man | dark bluish-grey | slate | 0.23  5 | 0.28  7 | 0.264 | 0.214 | 0.287 |
| Conirostrum  rufum | Rufous-browed  Conebill | M | man | plumbeous grey | slate | 0.19  5 | 0.24  8 | 0.281 | 0.275 | 0.248 |
| Conirostrum  sitticolor | Blue-backed  Conebill | M | man | blue | slate | 0.35  5 | 0.28  7 | 0.201 | 0.158 | 0.287 |
| Conirostrum  speciosum | Chestnut-vented  Conebill | M | man | dark greyish-blue | slate | 0.22  5 | 0.27  8 | 0.267 | 0.230 | 0.278 |
| Delothraupis  castaneoventris | Chestnut-bellied  Mountain-Tanager | M | man | dark blue | slate | 0.26  8 | 0.27  7 | 0.241 | 0.214 | 0.277 |
| Diglossa baritula | Cinnamon-bellied  Flowerpiercer | M | man | slate gray | slate | 0.23  6 | 0.27  1 | 0.263 | 0.230 | 0.271 |
| Diglossa  caerulescens | Bluish  Flowerpiercer | M | man | dull bluish-grey | slate | 0.24  2 | 0.26  8 | 0.262 | 0.229 | 0.268 |
| Diglossa duidae | Scaled  Flowerpiercer | M | man | slaty black | slate | 0.23  2 | 0.24  6 | 0.266 | 0.255 | 0.246 |
| Diglossa  plumbea | Slaty  Flowerpiercer | M | man | blackish-grey /  blue | slate | 0.23  8 | 0.26  5 | 0.263 | 0.234 | 0.265 |
| Euneornis  campestris | Orangequit | M | man | grey-blue | slate | 0.25  4 | 0.28  6 | 0.252 | 0.208 | 0.286 |
| Haplospiza  unicolor | Uniform Finch | M | man | blue-grey | slate | 0.17  0 | 0.26  0 | 0.291 | 0.279 | 0.260 |
| Hemispingus  goeringi | Slaty-backed  Hemispingus | M | man | slaty | slate | 0.18  6 | 0.24  0 | 0.285 | 0.289 | 0.240 |
| Nemosia pileata | Hooded Tanager | M | man | light blue | slate | 0.21  5 | 0.26  9 | 0.273 | 0.242 | 0.269 |
| Oreomanes  fraseri | Giant Conebill | M | man | plumbeous | slate | 0.21  0 | 0.24  2 | 0.274 | 0.274 | 0.242 |
| Phrygilus  erythronotus | White-throated  Sierra-Finch | M | man | dark gray | slate | 0.20  5 | 0.24  3 | 0.274 | 0.279 | 0.243 |
| Phrygilus  unicolor | Plumbeous Sierra-  Finch | M | man | lead-grey | slate | 0.20  9 | 0.25  1 | 0.273 | 0.267 | 0.251 |
| Poospiza cinerea | Cinereous  Warbling-Finch | M | man | blue-grey | slate | 0.21  0 | 0.24  6 | 0.272 | 0.272 | 0.246 |
| Poospiza whitii | Black-and- chestnut  Warbling-Finch | M | man | slate-grey | slate | 0.16  8 | 0.24  8 | 0.294 | 0.290 | 0.248 |
| Pyrrhocoma  ruficeps | Chestnut-headed  Tanager | M | man | dark grey | slate | 0.21  3 | 0.24  6 | 0.272 | 0.269 | 0.246 |
| Saltator  fuliginosus | Black-throated  Grosbeak | M | man | deep slate-blue | slate | 0.23  4 | 0.26  0 | 0.265 | 0.240 | 0.260 |
| Saltator grossus | Slate-colored  Grosbeak | M | man | slaty blue | slate | 0.22  8 | 0.24  3 | 0.264 | 0.265 | 0.243 |
| Schistochlamys  ruficapillus | Cinnamon Tanager | M | man | grey | slate | 0.19  0 | 0.25  3 | 0.282 | 0.275 | 0.253 |
| Sporophila  castaneiventris | Chestnut-bellied  Seedeater | M | man | blue-grey | slate | 0.19  6 | 0.25  5 | 0.279 | 0.269 | 0.255 |
| Sporophila  intermedia | Gray Seedeater | M | man | medium-grey /  blue | slate | 0.20  0 | 0.25  1 | 0.276 | 0.273 | 0.251 |
| Tangara heinei | Black-capped  Tanager | M | man | shining grey-blue | slate | 0.05  2 | 0.34  8 | 0.339 | 0.261 | 0.348 |
| Thraupis  episcopus | Blue-gray Tanager | M | man | darker bluish-grey | slate | 0.20  8 | 0.27  3 | 0.291 | 0.228 | 0.273 |
| Xenospingus  concolor | Slender-billed  Finch | M | man | slate-grey | slate | 0.18  7 | 0.25  4 | 0.283 | 0.276 | 0.254 |

| Acanthidops  bairdii | Peg-billed Finch | M | rum | dark grey | slate | 0.21  3 | 0.24  7 | 0.273 | 0.267 | 0.247 |
| --- | --- | --- | --- | --- | --- | --- | --- | --- | --- | --- |
| Bangsia arcaei | Blue-and-gold  Tanager | M | rum | dark blue | slate | 0.24  0 | 0.24  3 | 0.259 | 0.258 | 0.243 |
| Catamblyrhynch  us diadema | Plushcap | M | rum | blue gray | slate | 0.21  3 | 0.24  6 | 0.272 | 0.270 | 0.246 |
| Catamenia analis | Band-tailed  Seedeater | M | rum | blue-grey | slate | 0.20  7 | 0.26  5 | 0.273 | 0.255 | 0.265 |
| Catamenia  homochroa | Paramo Seedeater | M | rum | dark slate gray | slate | 0.18  3 | 0.23  6 | 0.284 | 0.297 | 0.236 |
| Catamenia  inornata | Plain-colored  Seedeater | M | rum | grey | slate | 0.17  3 | 0.24  8 | 0.287 | 0.292 | 0.248 |
| Conirostrum  cinereum | Cinereous Conebill | M | rum | slate-gray | slate | 0.16  3 | 0.22  7 | 0.294 | 0.315 | 0.227 |
| Conirostrum  leucogenys | White-eared  Conebill | M | rum | dark bluish-grey | slate | 0.22  2 | 0.25  8 | 0.268 | 0.251 | 0.258 |
| Conirostrum  rufum | Rufous-browed  Conebill | M | rum | plumbeous grey | slate | 0.20  8 | 0.22  1 | 0.270 | 0.300 | 0.221 |
| Conirostrum  sitticolor | Blue-backed  Conebill | M | rum | blue | slate | 0.26  5 | 0.25  2 | 0.243 | 0.239 | 0.252 |
| Conirostrum  speciosum | Chestnut-vented  Conebill | M | rum | dark greyish-blue | slate | 0.23  4 | 0.28  8 | 0.264 | 0.214 | 0.288 |
| Delothraupis  castaneoventris | Chestnut-bellied  Mountain-Tanager | M | rum | dark blue | slate | 0.26  8 | 0.27  0 | 0.242 | 0.220 | 0.270 |
| Diglossa baritula | Cinnamon-bellied  Flowerpiercer | M | rum | slate gray | slate | 0.23  7 | 0.26  1 | 0.262 | 0.240 | 0.261 |
| Diglossa  caerulescens | Bluish  Flowerpiercer | M | rum | dull bluish-grey | slate | 0.25  6 | 0.26  3 | 0.257 | 0.224 | 0.263 |
| Diglossa duidae | Scaled  Flowerpiercer | M | rum | slaty black | slate | 0.21  9 | 0.24  4 | 0.269 | 0.268 | 0.244 |
| Diglossa  plumbea | Slaty  Flowerpiercer | M | rum | blackish-grey /  blue | slate | 0.22  4 | 0.24  8 | 0.267 | 0.261 | 0.248 |
| Euneornis  campestris | Orangequit | M | rum | grey-blue | slate | 0.26  8 | 0.29  5 | 0.246 | 0.191 | 0.295 |
| Haplospiza  unicolor | Uniform Finch | M | rum | blue-grey | slate | 0.19  5 | 0.25  8 | 0.283 | 0.264 | 0.258 |
| Hemispingus  goeringi | Slaty-backed  Hemispingus | M | rum | dark grey | slate | 0.18  6 | 0.23  0 | 0.282 | 0.302 | 0.230 |
| Nemosia pileata | Hooded Tanager | M | rum | light blue | slate | 0.21  6 | 0.26  7 | 0.273 | 0.244 | 0.267 |
| Oreomanes  fraseri | Giant Conebill | M | rum | plumbeous | slate | 0.19  1 | 0.23  3 | 0.278 | 0.298 | 0.233 |
| Phrygilus  alaudinus | Band-tailed Sierra-  Finch | M | rum | grey | slate | 0.17  3 | 0.23  1 | 0.286 | 0.311 | 0.231 |
| Phrygilus  erythronotus | White-throated  Sierra-Finch | M | rum | dark gray | slate | 0.20  4 | 0.25  0 | 0.274 | 0.271 | 0.250 |
| Phrygilus  unicolor | Plumbeous Sierra-  Finch | M | rum | lead-grey | slate | 0.19  5 | 0.25  1 | 0.278 | 0.276 | 0.251 |
| Poospiza cinerea | Cinereous  Warbling-Finch | M | rum | blue-grey | slate | 0.18  2 | 0.25  9 | 0.284 | 0.275 | 0.259 |
| Poospiza whitii | Black-and- chestnut  Warbling-Finch | M | rum | slate-grey | slate | 0.21  7 | 0.24  9 | 0.271 | 0.263 | 0.249 |
| Pyrrhocoma  ruficeps | Chestnut-headed  Tanager | M | rum | dark grey | slate | 0.21  1 | 0.25  0 | 0.273 | 0.266 | 0.250 |
| Saltator  fuliginosus | Black-throated  Grosbeak | M | rum | deep slate-blue | slate | 0.24  2 | 0.25  1 | 0.263 | 0.245 | 0.251 |
| Saltator grossus | Slate-colored  Grosbeak | M | rum | slaty blue | slate | 0.22  6 | 0.24  4 | 0.266 | 0.264 | 0.244 |
| Schistochlamys  ruficapillus | Cinnamon Tanager | M | rum | grey | slate | 0.19  1 | 0.24  3 | 0.278 | 0.288 | 0.243 |
| Sporophila  castaneiventris | Chestnut-bellied  Seedeater | M | rum | blue-grey | slate | 0.19  5 | 0.26  0 | 0.278 | 0.267 | 0.260 |
| Sporophila  intermedia | Gray Seedeater | M | rum | medium-grey /  blue | slate | 0.20  1 | 0.25  0 | 0.276 | 0.273 | 0.250 |
| Tangara heinei | Black-capped  Tanager | M | rum | shining grey-blue | slate | 0.11  7 | 0.31  2 | 0.300 | 0.271 | 0.312 |
| Tangara  vitriolina | Scrub Tanager | M | rum | greyish blue | slate | 0.07  7 | 0.24  6 | 0.350 | 0.327 | 0.246 |

| Thraupis  episcopus | Blue-gray Tanager | M | rum | darker bluish-grey | slate | 0.22  4 | 0.29  5 | 0.265 | 0.217 | 0.295 |
| --- | --- | --- | --- | --- | --- | --- | --- | --- | --- | --- |
| Xenospingus  concolor | Slender-billed  Finch | M | rum | slate-grey | slate | 0.20  5 | 0.24  5 | 0.273 | 0.276 | 0.245 |
| Acanthidops  bairdii | Peg-billed Finch | M | thr | dark grey | slate | 0.19  3 | 0.24  7 | 0.280 | 0.280 | 0.247 |
| Bangsia arcaei | Blue-and-gold  Tanager | M | thr | dark blue | slate | 0.22  9 | 0.23  3 | 0.273 | 0.265 | 0.233 |
| Catamenia analis | Band-tailed  Seedeater | M | thr | grey | slate | 0.17  4 | 0.26  1 | 0.288 | 0.277 | 0.261 |
| Catamenia  homochroa | Paramo Seedeater | M | thr | dark slate gray | slate | 0.15  7 | 0.24  6 | 0.298 | 0.299 | 0.246 |
| Catamenia  inornata | Plain-colored  Seedeater | M | thr | grey | slate | 0.15  2 | 0.25  1 | 0.295 | 0.303 | 0.251 |
| Conirostrum  leucogenys | White-eared  Conebill | M | thr | pale grey / bluish | slate | 0.18  9 | 0.25  5 | 0.283 | 0.273 | 0.255 |
| Diglossa baritula | Cinnamon-bellied  Flowerpiercer | M | thr | slate-blackish /  cinnamon-rufous | slate | 0.14  6 | 0.20  3 | 0.292 | 0.359 | 0.203 |
| Diglossa  caerulescens | Bluish  Flowerpiercer | M | thr | dull bluish-grey | slate | 0.21  9 | 0.27  0 | 0.273 | 0.237 | 0.270 |
| Diglossa duidae | Scaled  Flowerpiercer | M | thr | slate-grey | slate | 0.21  2 | 0.24  7 | 0.273 | 0.269 | 0.247 |
| Diglossa  plumbea | Slaty  Flowerpiercer | M | thr | dark slate-grey | slate | 0.19  0 | 0.24  7 | 0.281 | 0.282 | 0.247 |
| Haplospiza  unicolor | Uniform Finch | M | thr | blue-grey | slate | 0.15  9 | 0.25  9 | 0.294 | 0.287 | 0.259 |
| Phrygilus gayi | Gray-hooded  Sierra-Finch | M | thr | bluish-grey | slate | 0.22  2 | 0.25  9 | 0.267 | 0.252 | 0.259 |
| Phrygilus  patagonicus | Patagonian Sierra-  Finch | M | thr | dark blue gray | slate | 0.23  3 | 0.25  6 | 0.262 | 0.249 | 0.256 |
| Phrygilus  punensis | Peruvian Sierra-  Finch | M | thr | grey | slate | 0.19  6 | 0.25  3 | 0.277 | 0.273 | 0.253 |
| Phrygilus  unicolor | Plumbeous Sierra-  Finch | M | thr | lead-grey | slate | 0.17  5 | 0.25  0 | 0.285 | 0.290 | 0.250 |
| Saltator  fuliginosus | Black-throated  Grosbeak | M | thr | deep slate-blue | slate | 0.20  2 | 0.23  5 | 0.287 | 0.277 | 0.235 |
| Tangara  vitriolina | Scrub Tanager | M | thr | pale bluish-grey | slate | 0.12  1 | 0.24  3 | 0.314 | 0.322 | 0.243 |
| Thraupis  cyanocephala | Blue-capped  Tanager | M | thr | dull bluish-grey | slate | 0.17  5 | 0.25  5 | 0.286 | 0.284 | 0.255 |
| Thraupis  episcopus | Blue-gray Tanager | M | thr | pale grey / blue  wash | slate | 0.19  1 | 0.25  7 | 0.286 | 0.266 | 0.257 |
| Acanthidops  bairdii | Peg-billed Finch | M | bre | dark grey | slate | 0.18  9 | 0.24  9 | 0.280 | 0.281 | 0.249 |
| Catamenia analis | Band-tailed  Seedeater | M | bre | grey | slate | 0.17  5 | 0.26  0 | 0.285 | 0.281 | 0.260 |
| Catamenia  homochroa | Paramo Seedeater | M | bre | dark slate gray | slate | 0.16  8 | 0.24  4 | 0.289 | 0.298 | 0.244 |
| Catamenia  inornata | Plain-colored  Seedeater | M | bre | grey | slate | 0.13  3 | 0.24  6 | 0.302 | 0.318 | 0.246 |
| Diglossa  plumbea | Slaty  Flowerpiercer | M | bre | dark slate-grey | slate | 0.19  4 | 0.25  5 | 0.280 | 0.271 | 0.255 |
| Euneornis  campestris | Orangequit | M | bre | grey-blue | slate | 0.23  7 | 0.28  7 | 0.264 | 0.213 | 0.287 |
| Haplospiza  unicolor | Uniform Finch | M | bre | blue-grey | slate | 0.19  7 | 0.25  8 | 0.276 | 0.269 | 0.258 |
| Phrygilus  erythronotus | White-throated  Sierra-Finch | M | bre | grey | slate | 0.16  6 | 0.25  0 | 0.285 | 0.299 | 0.250 |
| Phrygilus  unicolor | Plumbeous Sierra-  Finch | M | bre | lead-grey | slate | 0.19  2 | 0.25  5 | 0.276 | 0.277 | 0.255 |
| Pyrrhocoma  ruficeps | Chestnut-headed  Tanager | M | bre | dark grey | slate | 0.19  1 | 0.25  1 | 0.281 | 0.277 | 0.251 |
| Tangara  vitriolina | Scrub Tanager | M | bre | pale bluish-grey | slate | 0.06  6 | 0.26  2 | 0.341 | 0.330 | 0.262 |
| Thraupis  cyanocephala | Blue-capped  Tanager | M | bre | dull bluish-grey | slate | 0.17  4 | 0.26  0 | 0.286 | 0.280 | 0.260 |
| Thraupis  episcopus | Blue-gray Tanager | M | bre | pale grey / blue  wash | slate | 0.20  8 | 0.28  1 | 0.269 | 0.243 | 0.281 |
| Acanthidops | Peg-billed Finch | M | bel | dark grey | slate | 0.17 | 0.25 | 0.284 | 0.286 | 0.251 |

| bairdii |  |  |  |  |  | 9 | 1 |  |  |  |
| --- | --- | --- | --- | --- | --- | --- | --- | --- | --- | --- |
| Catamenia  homochroa | Paramo Seedeater | M | bel | dark slate gray | slate | 0.15  3 | 0.24  0 | 0.296 | 0.312 | 0.240 |
| Diglossa  plumbea | Slaty  Flowerpiercer | M | bel | dark slate-grey | slate | 0.17  3 | 0.24  8 | 0.286 | 0.293 | 0.248 |
| Euneornis  campestris | Orangequit | M | bel | grey-blue | slate | 0.23  1 | 0.27  4 | 0.265 | 0.230 | 0.274 |
| Haplospiza  unicolor | Uniform Finch | M | bel | blue-grey | slate | 0.15  9 | 0.25  5 | 0.295 | 0.292 | 0.255 |
| Phrygilus  unicolor | Plumbeous Sierra-  Finch | M | bel | lead-grey | slate | 0.17  9 | 0.25  0 | 0.283 | 0.287 | 0.250 |
| Pyrrhocoma  ruficeps | Chestnut-headed  Tanager | M | bel | dark grey | slate | 0.19  4 | 0.24  7 | 0.277 | 0.282 | 0.247 |
| Saltator  fuliginosus | Black-throated  Grosbeak | M | bel | deep slate-blue | slate | 0.22  3 | 0.24  8 | 0.268 | 0.261 | 0.248 |
| Saltator grossus | Slate-colored  Grosbeak | M | bel | slaty blue | slate | 0.20  9 | 0.24  9 | 0.272 | 0.270 | 0.249 |
| Tangara heinei | Black-capped  Tanager | M | bel | gray-blue | slate | 0.12  0 | 0.31  0 | 0.301 | 0.270 | 0.310 |
| Tangara  vitriolina | Scrub Tanager | M | bel | pale glaucous | slate | 0.07  0 | 0.21  8 | 0.339 | 0.374 | 0.218 |
| Thraupis  cyanocephala | Blue-capped  Tanager | M | bel | dull bluish-grey | slate | 0.16  7 | 0.25  9 | 0.286 | 0.288 | 0.259 |
| Thraupis  episcopus | Blue-gray Tanager | M | bel | pale grey / blue  wash | slate | 0.20  0 | 0.28  9 | 0.273 | 0.237 | 0.289 |
| Bangsia arcaei | Blue-and-gold  Tanager | F | cro | bright blue | slate | 0.32  5 | 0.27  8 | 0.218 | 0.180 | 0.278 |
| Conirostrum  cinereum | Cinereous Conebill | F | cro | slate-gray | slate | 0.14  9 | 0.22  4 | 0.303 | 0.324 | 0.224 |
| Conirostrum  ferrugineiventre | White-browed  Conebill | F | cro | slaty | slate | 0.17  1 | 0.24  4 | 0.296 | 0.290 | 0.244 |
| Conirostrum  rufum | Rufous-browed  Conebill | F | cro | plumbeous grey | slate | 0.15  6 | 0.24  0 | 0.297 | 0.306 | 0.240 |
| Conirostrum  speciosum | Chestnut-vented  Conebill | F | cro | bluish-grey | slate | 0.17  1 | 0.26  4 | 0.298 | 0.267 | 0.264 |
| Delothraupis  castaneoventris | Chestnut-bellied  Mountain-Tanager | F | cro | silvery sky blue | slate | 0.27  7 | 0.29  5 | 0.237 | 0.190 | 0.295 |
| Diglossa  caerulescens | Bluish  Flowerpiercer | F | cro | dull bluish-grey | slate | 0.20  0 | 0.25  5 | 0.279 | 0.266 | 0.255 |
| Diglossa duidae | Scaled  Flowerpiercer | F | cro | slaty black | slate | 0.19  9 | 0.24  6 | 0.290 | 0.265 | 0.246 |
| Eucometis  penicillata | Gray-headed  Tanager | F | cro | grey | slate | 0.09  5 | 0.23  8 | 0.321 | 0.345 | 0.238 |
| Euneornis  campestris | Orangequit | F | cro | olive-grey | slate | 0.14  2 | 0.24  3 | 0.311 | 0.303 | 0.243 |
| Nemosia pileata | Hooded Tanager | F | cro | light blue | slate | 0.16  6 | 0.26  2 | 0.299 | 0.272 | 0.262 |
| Phrygilus  erythronotus | White-throated  Sierra-Finch | F | cro | dark gray | slate | 0.18  0 | 0.24  6 | 0.284 | 0.289 | 0.246 |
| Phrygilus  patagonicus | Patagonian Sierra-  Finch | F | cro | dark blue gray | slate | 0.22  7 | 0.26  1 | 0.266 | 0.247 | 0.261 |
| Phrygilus  punensis | Peruvian Sierra-  Finch | F | cro | grey | slate | 0.16  3 | 0.23  9 | 0.294 | 0.305 | 0.239 |
| Poospiza cinerea | Cinereous  Warbling-Finch | F | cro | grey | slate | 0.16  9 | 0.25  1 | 0.288 | 0.292 | 0.251 |
| Poospiza  erythrophrys | Rusty-browed  Warbling-Finch | F | cro | blue-grey | slate | 0.15  1 | 0.24  5 | 0.294 | 0.311 | 0.245 |
| Poospiza whitii | Black-and- chestnut  Warbling-Finch | F | cro | slate-grey / olive- brown | slate | 0.15  5 | 0.22  9 | 0.300 | 0.316 | 0.229 |
| Saltator  fuliginosus | Black-throated  Grosbeak | F | cro | deep slate-blue | slate | 0.19  5 | 0.24  9 | 0.282 | 0.274 | 0.249 |
| Saltator grossus | Slate-colored  Grosbeak | F | cro | slaty blue | slate | 0.19  4 | 0.25  1 | 0.285 | 0.270 | 0.251 |
| Thraupis  episcopus | Blue-gray Tanager | F | cro | pale grey / blue  wash | slate | 0.18  1 | 0.24  8 | 0.296 | 0.274 | 0.248 |
| Xenospingus  concolor | Slender-billed  Finch | F | cro | slate-grey | slate | 0.16  5 | 0.25  7 | 0.291 | 0.287 | 0.257 |

| Bangsia arcaei | Blue-and-gold  Tanager | F | nap | dark blue | slate | 0.32  1 | 0.26  6 | 0.221 | 0.192 | 0.266 |
| --- | --- | --- | --- | --- | --- | --- | --- | --- | --- | --- |
| Conirostrum  cinereum | Cinereous Conebill | F | nap | slate-gray | slate | 0.14  6 | 0.22  5 | 0.302 | 0.326 | 0.225 |
| Conirostrum  ferrugineiventre | White-browed  Conebill | F | nap | blue gray | slate | 0.18  8 | 0.25  0 | 0.285 | 0.277 | 0.250 |
| Conirostrum  rufum | Rufous-browed  Conebill | F | nap | plumbeous grey | slate | 0.16  6 | 0.24  3 | 0.293 | 0.297 | 0.243 |
| Conirostrum  sitticolor | Blue-backed  Conebill | F | nap | blue | slate | 0.20  0 | 0.24  5 | 0.282 | 0.274 | 0.245 |
| Conirostrum  speciosum | Chestnut-vented  Conebill | F | nap | bluish-grey | slate | 0.19  4 | 0.26  7 | 0.285 | 0.254 | 0.267 |
| Delothraupis  castaneoventris | Chestnut-bellied  Mountain-Tanager | F | nap | dark blue | slate | 0.26  5 | 0.27  3 | 0.244 | 0.218 | 0.273 |
| Diglossa  caerulescens | Bluish  Flowerpiercer | F | nap | dull bluish-grey | slate | 0.21  3 | 0.24  8 | 0.272 | 0.267 | 0.248 |
| Diglossa duidae | Scaled  Flowerpiercer | F | nap | slaty black | slate | 0.24  3 | 0.25  3 | 0.265 | 0.238 | 0.253 |
| Eucometis  penicillata | Gray-headed  Tanager | F | nap | grey / green | slate | 0.06  4 | 0.23  3 | 0.338 | 0.365 | 0.233 |
| Euneornis  campestris | Orangequit | F | nap | olive-grey | slate | 0.14  8 | 0.23  9 | 0.306 | 0.306 | 0.239 |
| Nemosia pileata | Hooded Tanager | F | nap | light blue | slate | 0.19  6 | 0.26  2 | 0.282 | 0.261 | 0.262 |
| Phrygilus  erythronotus | White-throated  Sierra-Finch | F | nap | dark gray | slate | 0.17  6 | 0.24  9 | 0.286 | 0.288 | 0.249 |
| Phrygilus  patagonicus | Patagonian Sierra-  Finch | F | nap | dark blue gray | slate | 0.21  7 | 0.24  9 | 0.271 | 0.263 | 0.249 |
| Phrygilus  punensis | Peruvian Sierra-  Finch | F | nap | grey | slate | 0.17  8 | 0.24  0 | 0.286 | 0.296 | 0.240 |
| Poospiza cinerea | Cinereous  Warbling-Finch | F | nap | grey | slate | 0.13  7 | 0.25  6 | 0.302 | 0.305 | 0.256 |
| Poospiza whitii | Black-and- chestnut  Warbling-Finch | F | nap | slate-grey / olive- brown | slate | 0.15  5 | 0.22  2 | 0.299 | 0.324 | 0.222 |
| Saltator  fuliginosus | Black-throated  Grosbeak | F | nap | deep slate-blue | slate | 0.21  7 | 0.25  1 | 0.272 | 0.261 | 0.251 |
| Saltator grossus | Slate-colored  Grosbeak | F | nap | slaty blue | slate | 0.21  7 | 0.25  1 | 0.272 | 0.259 | 0.251 |
| Thraupis  episcopus | Blue-gray Tanager | F | nap | pale grey / blue  wash | slate | 0.19  0 | 0.25  1 | 0.293 | 0.265 | 0.251 |
| Xenospingus  concolor | Slender-billed  Finch | F | nap | slate-grey | slate | 0.18  2 | 0.25  5 | 0.283 | 0.279 | 0.255 |
| Bangsia arcaei | Blue-and-gold  Tanager | F | man | dark blue | slate | 0.26  8 | 0.24  8 | 0.247 | 0.238 | 0.248 |
| Catamblyrhynch  us diadema | Plushcap | F | man | blue gray | slate | 0.21  8 | 0.25  2 | 0.270 | 0.261 | 0.252 |
| Conirostrum  cinereum | Cinereous Conebill | F | man | slate-gray | slate | 0.16  4 | 0.22  4 | 0.294 | 0.319 | 0.224 |
| Conirostrum  ferrugineiventre | White-browed  Conebill | F | man | blue gray | slate | 0.18  5 | 0.26  3 | 0.286 | 0.265 | 0.263 |
| Conirostrum  rufum | Rufous-browed  Conebill | F | man | plumbeous grey | slate | 0.16  6 | 0.24  1 | 0.294 | 0.299 | 0.241 |
| Conirostrum  sitticolor | Blue-backed  Conebill | F | man | blue | slate | 0.26  9 | 0.25  9 | 0.243 | 0.229 | 0.259 |
| Delothraupis  castaneoventris | Chestnut-bellied  Mountain-Tanager | F | man | dark blue | slate | 0.26  5 | 0.28  8 | 0.242 | 0.205 | 0.288 |
| Diglossa  caerulescens | Bluish  Flowerpiercer | F | man | dull bluish-grey | slate | 0.22  6 | 0.27  0 | 0.268 | 0.236 | 0.270 |
| Diglossa duidae | Scaled  Flowerpiercer | F | man | slaty black | slate | 0.24  6 | 0.24  6 | 0.261 | 0.247 | 0.246 |
| Hemispingus  goeringi | Slaty-backed  Hemispingus | F | man | slaty | slate | 0.19  3 | 0.23  8 | 0.282 | 0.287 | 0.238 |
| Nemosia pileata | Hooded Tanager | F | man | light blue | slate | 0.19  1 | 0.26  2 | 0.283 | 0.264 | 0.262 |
| Phrygilus  erythronotus | White-throated  Sierra-Finch | F | man | dark gray | slate | 0.18  6 | 0.24  7 | 0.282 | 0.285 | 0.247 |
| Poospiza cinerea | Cinereous  Warbling-Finch | F | man | blue-grey | slate | 0.14  5 | 0.25  8 | 0.300 | 0.297 | 0.258 |

| Poospiza whitii | Black-and- chestnut  Warbling-Finch | F | man | slate-grey / olive- brown | slate | 0.15  3 | 0.22  9 | 0.300 | 0.318 | 0.229 |
| --- | --- | --- | --- | --- | --- | --- | --- | --- | --- | --- |
| Saltator  fuliginosus | Black-throated  Grosbeak | F | man | deep slate-blue | slate | 0.22  0 | 0.25  1 | 0.271 | 0.257 | 0.251 |
| Saltator grossus | Slate-colored  Grosbeak | F | man | slaty blue | slate | 0.21  6 | 0.25  4 | 0.273 | 0.256 | 0.254 |
| Schistochlamys  ruficapillus | Cinnamon Tanager | F | man | grey | slate | 0.17  2 | 0.22  8 | 0.293 | 0.307 | 0.228 |
| Thraupis  episcopus | Blue-gray Tanager | F | man | darker bluish-grey | slate | 0.18  8 | 0.25  3 | 0.295 | 0.264 | 0.253 |
| Xenospingus  concolor | Slender-billed  Finch | F | man | slate-grey | slate | 0.18  5 | 0.25  3 | 0.282 | 0.281 | 0.253 |
| Bangsia arcaei | Blue-and-gold  Tanager | F | rum | dark blue | slate | 0.32  4 | 0.26  7 | 0.220 | 0.188 | 0.267 |
| Catamblyrhynch  us diadema | Plushcap | F | rum | blue gray | slate | 0.20  4 | 0.23  7 | 0.273 | 0.287 | 0.237 |
| Catamenia analis | Band-tailed  Seedeater | F | rum | brown / dark  brown | slate | 0.17  2 | 0.23  1 | 0.289 | 0.308 | 0.231 |
| Conirostrum  cinereum | Cinereous Conebill | F | rum | slate-gray | slate | 0.15  9 | 0.21  8 | 0.293 | 0.330 | 0.218 |
| Conirostrum  ferrugineiventre | White-browed  Conebill | F | rum | blue gray | slate | 0.19  7 | 0.26  0 | 0.281 | 0.262 | 0.260 |
| Conirostrum  rufum | Rufous-browed  Conebill | F | rum | plumbeous grey | slate | 0.17  3 | 0.23  6 | 0.286 | 0.305 | 0.236 |
| Conirostrum  sitticolor | Blue-backed  Conebill | F | rum | blue | slate | 0.22  1 | 0.24  2 | 0.262 | 0.275 | 0.242 |
| Delothraupis  castaneoventris | Chestnut-bellied  Mountain-Tanager | F | rum | dark blue | slate | 0.22  9 | 0.25  2 | 0.259 | 0.260 | 0.252 |
| Diglossa  caerulescens | Bluish  Flowerpiercer | F | rum | dull bluish-grey | slate | 0.24  5 | 0.26  1 | 0.260 | 0.234 | 0.261 |
| Diglossa duidae | Scaled  Flowerpiercer | F | rum | slaty black | slate | 0.24  1 | 0.24  4 | 0.263 | 0.252 | 0.244 |
| Hemispingus  goeringi | Slaty-backed  Hemispingus | F | rum | dark grey | slate | 0.18  9 | 0.23  1 | 0.281 | 0.299 | 0.231 |
| Nemosia pileata | Hooded Tanager | F | rum | light blue | slate | 0.20  2 | 0.25  9 | 0.278 | 0.260 | 0.259 |
| Phrygilus  erythronotus | White-throated  Sierra-Finch | F | rum | dark gray | slate | 0.22  6 | 0.24  9 | 0.266 | 0.258 | 0.249 |
| Poospiza cinerea | Cinereous  Warbling-Finch | F | rum | blue-grey | slate | 0.16  6 | 0.26  0 | 0.292 | 0.281 | 0.260 |
| Poospiza whitii | Black-and- chestnut  Warbling-Finch | F | rum | slate-grey / olive- brown | slate | 0.19  5 | 0.23  6 | 0.281 | 0.287 | 0.236 |
| Saltator  fuliginosus | Black-throated  Grosbeak | F | rum | deep slate-blue | slate | 0.20  7 | 0.25  0 | 0.276 | 0.267 | 0.250 |
| Saltator grossus | Slate-colored  Grosbeak | F | rum | slaty blue | slate | 0.21  0 | 0.24  6 | 0.275 | 0.270 | 0.246 |
| Schistochlamys  ruficapillus | Cinnamon Tanager | F | rum | grey | slate | 0.19  1 | 0.23  2 | 0.279 | 0.298 | 0.232 |
| Tangara  vitriolina | Scrub Tanager | F | rum | greyish blue | slate | 0.11  1 | 0.23  1 | 0.335 | 0.322 | 0.231 |
| Thraupis  episcopus | Blue-gray Tanager | F | rum | darker bluish-grey | slate | 0.22  4 | 0.26  6 | 0.286 | 0.224 | 0.266 |
| Xenospingus  concolor | Slender-billed  Finch | F | rum | slate-grey | slate | 0.19  5 | 0.25  4 | 0.277 | 0.274 | 0.254 |
| Bangsia arcaei | Blue-and-gold  Tanager | F | thr | dark blue | slate | 0.23  4 | 0.22  0 | 0.273 | 0.273 | 0.220 |
| Diglossa  caerulescens | Bluish  Flowerpiercer | F | thr | dull bluish-grey | slate | 0.16  2 | 0.22  0 | 0.288 | 0.329 | 0.220 |
| Diglossa duidae | Scaled  Flowerpiercer | F | thr | slate-grey | slate | 0.22  3 | 0.25  2 | 0.272 | 0.254 | 0.252 |
| Embernagra  platensis | Great Pampa-  Finch | F | thr | grey | slate | 0.12  1 | 0.23  4 | 0.304 | 0.341 | 0.234 |
| Phrygilus  patagonicus | Patagonian Sierra-  Finch | F | thr | dark blue gray | slate | 0.22  0 | 0.25  2 | 0.268 | 0.260 | 0.252 |
| Saltator  fuliginosus | Black-throated  Grosbeak | F | thr | deep slate-blue | slate | 0.17  0 | 0.22  3 | 0.294 | 0.313 | 0.223 |
| Tangara | Scrub Tanager | F | thr | pale bluish-grey | slate | 0.12 | 0.23 | 0.316 | 0.323 | 0.236 |

| vitriolina |  |  |  |  |  | 5 | 6 |  |  |  |
| --- | --- | --- | --- | --- | --- | --- | --- | --- | --- | --- |
| Thraupis  cyanocephala | Blue-capped  Tanager | F | thr | dull bluish-grey | slate | 0.15  4 | 0.25  2 | 0.295 | 0.300 | 0.252 |
| Thraupis  episcopus | Blue-gray Tanager | F | thr | pale grey / blue  wash | slate | 0.16  4 | 0.25  0 | 0.292 | 0.293 | 0.250 |
| Diglossa  caerulescens | Bluish  Flowerpiercer | F | bre | dull bluish-grey | slate | 0.22  1 | 0.26  9 | 0.267 | 0.242 | 0.269 |
| Diglossa duidae | Scaled  Flowerpiercer | F | bre | slate-grey | slate | 0.23  0 | 0.25  3 | 0.267 | 0.250 | 0.253 |
| Embernagra  platensis | Great Pampa-  Finch | F | bre | grey | slate | 0.10  9 | 0.23  8 | 0.310 | 0.343 | 0.238 |
| Phrygilus  erythronotus | White-throated  Sierra-Finch | F | bre | grey | slate | 0.17  3 | 0.25  1 | 0.283 | 0.293 | 0.251 |
| Saltator  fuliginosus | Black-throated  Grosbeak | F | bre | deep slate-blue | slate | 0.20  3 | 0.23  7 | 0.280 | 0.280 | 0.237 |
| Saltator grossus | Slate-colored  Grosbeak | F | bre | slaty blue | slate | 0.14  4 | 0.22  6 | 0.301 | 0.330 | 0.226 |
| Thraupis  cyanocephala | Blue-capped  Tanager | F | bre | dull bluish-grey | slate | 0.16  3 | 0.26  1 | 0.289 | 0.286 | 0.261 |
| Thraupis  episcopus | Blue-gray Tanager | F | bre | pale grey / blue  wash | slate | 0.18  6 | 0.27  7 | 0.289 | 0.248 | 0.277 |
| Saltator  fuliginosus | Black-throated  Grosbeak | F | bel | deep slate-blue | slate | 0.20  0 | 0.24  4 | 0.278 | 0.277 | 0.244 |
| Saltator grossus | Slate-colored  Grosbeak | F | bel | slaty blue | slate | 0.16  5 | 0.23  4 | 0.290 | 0.311 | 0.234 |
| Tangara  vitriolina | Scrub Tanager | F | bel | pale glaucous | slate | 0.08  9 | 0.23  8 | 0.329 | 0.344 | 0.238 |
| Thraupis  cyanocephala | Blue-capped  Tanager | F | bel | dull bluish-grey | slate | 0.16  4 | 0.25  6 | 0.289 | 0.291 | 0.256 |
| Thraupis  episcopus | Blue-gray Tanager | F | bel | pale grey / blue  wash | slate | 0.17  9 | 0.26  3 | 0.289 | 0.269 | 0.263 |

**Table S2.** Scoring of the plumage colour measurements for males of each species with double scoring species involved. Species with double scoring are marked in red and excluded from the second analysis.

| Latin name | cod  ing | Latin name | cod  ing | Latin name | cod  ing | Latin name | cod  ing | Latin name | cod  ing | Latin  name | cod  ing |
| --- | --- | --- | --- | --- | --- | --- | --- | --- | --- | --- | --- |
| Acanthidops bairdii | 2 | Dacnis nigripes | 3 | Incaspiza pulchra | 1 | Poospiza caesar | 1 | Sporophila albogularis | 1 | Tangara icteroceph  ala | 0 |
| Anisognathu  s igniventris | 3 | Dacnis  venusta | 3 | Iridophanes  pulcherrimus | 3 | Poospiza  cinerea | 2 | Sporophila  americana | 0 | Tangara  inornata | 3 |
| Anisognathu  s lacrymosus | 3 | Dacnis  viguieri | 3 | Iridosornis  analis | 3 | Poospiza  erythrophrys | 2 | Sporophila  bouvreuil | 0 | Tangara  johannae | 3 |
| Anisognathu s  melanogenys | 3 | Delothraupis castaneovent  ris | 2 | Iridosornis jelskii | 3 | Poospiza hispaniolensi  s | 1 | Sporophila bouvronides | 0 | Tangara labradorid  es | 3 |
| Anisognathu s  somptuosus | 3 | Diglossa albilatera | 3 | Iridosornis porphyrocep  halus | 3 | Poospiza hypochondri  a | 1 | Sporophila caerulescens | 1 | Tangara larvata | 3 |
| Bangsia arcaei | 2 | Diglossa baritula | 2 | Iridosornis rufivertex | 3 | Poospiza lateralis | 2 | Sporophila castaneivent  ris | 2 | Tangara lavinia | 3 |
| Bangsia edwardsi | 3 | Diglossa brunneiventri  s | 1 | Lanio aurantius | 0 | Poospiza melanoleuca | 1 | Sporophila cinnamomea | 1 | Tangara mexicana | 3 |
| Buthraupis eximia | 3 | Diglossa caerulescens | 2 | Lanio fulvus | 0 | Poospiza nigrorufa | 1 | Sporophila collaris | 1 | Tangara nigrocinct  a | 3 |
| Buthraupis montana | 3 | Diglossa carbonaria | 1 | Lanio leucothorax | 0 | Poospiza ornata | 1 | Sporophila corvina | 0 | Tangara nigroviridi  s | 3 |
| Calochaetes | 0 | Diglossa | 3 | Lanio | 0 | Poospiza | 1 | Sporophila | 1 | Tangara | 3 |

| coccineus |  | cyanea |  | versicolor |  | thoracica |  | frontalis |  | parzudakii |  |
| --- | --- | --- | --- | --- | --- | --- | --- | --- | --- | --- | --- |
| Camarhynch us pallidus | 0 | Diglossa duidae | 2 | Lophospingu s griseocristat  us | 1 | Poospiza torquata | 1 | Sporophila hypoxantha | 1 | Tangara peruviana | 3 |
| Camarhynch  us parvulus | 0 | Diglossa  glauca | 3 | Lophospingu  s pusillus | 1 | Poospiza  whitii | 2 | Sporophila  intermedia | 2 | Tangara  preciosa | 3 |
| Camarhynch us psittacula | 0 | Diglossa gloriosa | 1 | Loxigilla barbadensis | 1 | Porphyrospiz a  caerulescens | 3 | Sporophila leucoptera | 1 | Tangara punctata | 3 |
| Catamblyrhy nchus  diadema | 2 | Diglossa gloriosissima | 2 | Loxigilla noctis | 1 | Pyrrhocoma ruficeps | 2 | Sporophila lineola | 0 | Tangara ruficervix | 3 |
| Catamenia  analis | 2 | Diglossa  humeralis | 1 | Loxigilla  portoricensis | 0 | Ramphocelu  s bresilius | 0 | Sporophila  luctuosa | 0 | Tangara  rufigenis | 3 |
| Catamenia  homochroa | 2 | Diglossa  indigotica | 3 | Loxigilla  violacea | 0 | Ramphocelu  s carbo | 0 | Sporophila  minuta | 1 | Tangara  rufigula | 0 |
| Catamenia  inornata | 2 | Diglossa  lafresnayii | 2 | Loxipasser  anoxanthus | 0 | Ramphocelu  s dimidiatus | 0 | Sporophila  murallae | 0 | Tangara  schrankii | 3 |
| Certhidea olivacea | 0 | Diglossa major | 2 | Melanodera melanodera | 2 | Ramphocelu s  flammigerus | 0 | Sporophila nigricollis | 0 | Tangara seledon | 3 |
| Charitospiza eucosma | 1 | Diglossa mystacalis | 2 | Melanodera xanthogram ma | 1 | Ramphocelu s melanogaste  r | 0 | Sporophila nigrorufa | 0 | Tangara varia | 3 |
| Chlorochrysa calliparaea | 3 | Diglossa plumbea | 2 | Melanospiza richardsoni | 0 | Ramphocelu s  nigrogularis | 0 | Sporophila palustris | 1 | Tangara vassorii | 3 |
| Chlorochrysa  phoenicotis | 0 | Diglossa  sittoides | 2 | Melopyrrha  nigra | 0 | Ramphocelu  s passerinii | 0 | Sporophila  peruviana | 1 | Tangara  velia | 3 |
| Chlorophane s spiza | 3 | Diglossa venezuelensis | 0 | Nemosia pileata | 2 | Ramphocelu s sanguinolent  us | 0 | Sporophila plumbea | 1 | Tangara viridicollis | 3 |
| Chlorornis  riefferii | 0 | Diuca diuca | 1 | Neothraupis  fasciata | 1 | Rhodospingu  s cruentus | 0 | Sporophila  ruficollis | 1 | Tangara  vitriolina | 3 |
| Chrysothlypi s  chrysomelas | 0 | Donacospiza albifrons | 1 | Nesospiza acunhae | 0 | Rowettia goughensis | 0 | Sporophila simplex | 1 | Tangara xanthocep  hala | 3 |
| Chrysothlypi s salmoni | 0 | Dubusia taeniata | 3 | Nesospiza questi | 0 | Saltator albicollis | 0 | Sporophila telasco | 1 | Tangara xanthogast  ra | 0 |
| Cissopis  leverianus | 0 | Emberizoides  herbicola | 1 | Nesospiza  wilkinsi | 0 | Saltator  atriceps | 1 | Sporophila  torqueola | 0 | Tersina  viridis | 3 |
| Cnemoscopu s rubrirostris | 1 | Embernagra platensis | 2 | Orchesticus abeillei | 0 | Saltator atricollis | 0 | Stephanoph orus  diadematus | 3 | Thlypopsis fulviceps | 1 |
| Coereba  flaveola | 1 | Eucometis  penicillata | 2 | Oreomanes  fraseri | 2 | Saltator  atripennis | 1 | Tachyphonu  s coronatus | 0 | Thlypopsis  inornata | 1 |
| Compsospiza garleppi | 1 | Euneornis campestris | 2 | Oryzoborus angolensis | 0 | Saltator aurantiirostri  s | 1 | Tachyphonu s cristatus | 0 | Thlypopsis ornata | 1 |
| Conirostrum  albifrons | 3 | Geospiza  conirostris | 0 | Oryzoborus  crassirostris | 0 | Saltator  coerulescens | 1 | Tachyphonu  s delatrii | 0 | Thlypopsis  pectoralis | 1 |
| Conirostrum  bicolor | 3 | Geospiza  difficilis | 0 | Oryzoborus  maximiliani | 0 | Saltator  fuliginosus | 2 | Tachyphonu  s luctuosus | 0 | Thlypopsis  ruficeps | 0 |
| Conirostrum  cinereum | 2 | Geospiza  fortis | 0 | Oryzoborus  nuttingi | 0 | Saltator  grossus | 2 | Tachyphonu  s phoenicius | 0 | Thlypopsis  sordida | 1 |
| Conirostrum ferrugineive  ntre | 2 | Geospiza fuliginosa | 0 | Parkerthraus tes  humeralis | 2 | Saltator maxillosus | 1 | Tachyphonu s rufiventer | 0 | Thraupis abbas | 3 |
| Conirostrum leucogenys | 2 | Geospiza magnirostris | 0 | Paroaria capitata | 0 | Saltator maximus | 1 | Tachyphonu s rufus | 0 | Thraupis bonariensi  s | 3 |
| Conirostrum margaritae | 2 | Geospiza scandens | 0 | Paroaria coronata | 1 | Saltator nigriceps | 1 | Tachyphonu s surinamus | 0 | Thraupis cyanoceph  ala | 3 |
| Conirostrum | 2 | Gubernatrix | 0 | Paroaria | 1 | Saltator | 1 | Tangara | 3 | Thraupis | 3 |

| rufum |  | cristata |  | dominicana |  | orenocensis |  | argyrofenge  s |  | episcopus |  |
| --- | --- | --- | --- | --- | --- | --- | --- | --- | --- | --- | --- |
| Conirostrum sitticolor | 2 | Haplospiza unicolor | 2 | Paroaria gularis | 0 | Saltator similis | 1 | Tangara arthus | 0 | Thraupis glaucocolp  a | 3 |
| Conirostrum  speciosum | 2 | Hemispingus  atropileus | 0 | Phrygilus  alaudinus | 2 | Saltator  striatipectus | 1 | Tangara  callophrys | 3 | Thraupis  ornata | 3 |
| Conothraupi  s speculigera | 1 | Hemispingus  calophrys | 0 | Phrygilus  atriceps | 1 | Saltatricula  multicolor | 1 | Tangara  cayana | 3 | Thraupis  sayaca | 3 |
| Coryphaspiza  melanotis | 1 | Hemispingus  frontalis | 0 | Phrygilus  dorsalis | 2 | Schistochlam  ys melanopis | 1 | Tangara  chilensis | 3 | Tiaris  bicolor | 1 |
| Coryphospin gus  cucullatus | 0 | Hemispingus goeringi | 2 | Phrygilus erythronotus | 2 | Schistochlam ys  ruficapillus | 2 | Tangara chrysotis | 0 | Tiaris canorus | 1 |
| Coryphospin  gus pileatus | 1 | Hemispingus  melanotis | 1 | Phrygilus  fruticeti | 1 | Sericossypha  albocristata | 0 | Tangara  cucullata | 3 | Tiaris  fuliginosus | 0 |
| Creurgops  dentatus | 2 | Hemispingus  reyi | 1 | Phrygilus  gayi | 2 | Sicalis  auriventris | 0 | Tangara  cyanicollis | 3 | Tiaris  obscurus | 1 |
| Creurgops verticalis | 1 | Hemispingus superciliaris | 0 | Phrygilus patagonicus | 2 | Sicalis citrina | 0 | Tangara cyanocephal  a | 3 | Tiaris olivaceus | 0 |
| Cyanerpes caeruleus | 3 | Hemispingus verticalis | 1 | Phrygilus plebejus | 1 | Sicalis columbiana | 0 | Tangara cyanoptera | 3 | Trichothra upis  melanops | 0 |
| Cyanerpes cyaneus | 3 | Hemispingus xanthophthal  mus | 1 | Phrygilus punensis | 2 | Sicalis flaveola | 0 | Tangara cyanotis | 3 | Urothraupi s  stolzmanni | 1 |
| Cyanerpes  lucidus | 3 | Hemithraupis  flavicollis | 0 | Phrygilus  unicolor | 2 | Sicalis  lebruni | 1 | Tangara  cyanoventris | 3 | Volatinia  jacarina | 0 |
| Cyanerpes  nitidus | 3 | Hemithraupis  guira | 0 | Piezorhina  cinerea | 1 | Sicalis lutea | 0 | Tangara  desmaresti | 0 | Xenodacni  s parina | 3 |
| Cyanicterus cyanicterus | 3 | Hemithraupis ruficapilla | 0 | Pinaroloxias inornata | 0 | Sicalis luteocephala | 1 | Tangara dowii | 3 | Xenosping us  concolor | 2 |
| Cypsnagra  hirundinacea | 0 | Heterospingu  s rubrifrons | 1 | Pipraeidea  melanonota | 3 | Sicalis  luteola | 0 | Tangara  fastuosa | 3 |  |  |
| Dacnis cayana | 3 | Heterospingu s  xanthopygius | 0 | Platyspiza crassirostris | 0 | Sicalis olivascens | 0 | Tangara florida | 0 |  |  |
| Dacnis  flaviventer | 0 | Idiopsar  brachyurus | 1 | Poospiza  alticola | 1 | Sicalis  raimondii | 1 | Tangara  guttata | 3 |  |  |
| Dacnis  hartlaubi | 3 | Incaspiza  laeta | 1 | Poospiza  boliviana | 1 | Sicalis  taczanowskii | 1 | Tangara  gyrola | 3 |  |  |
| Dacnis  lineata | 3 | Incaspiza  personata | 1 | Poospiza  cabanisi | 1 | Sicalis  uropygialis | 1 | Tangara  heinei | 3 |  |  |

**Table S3.** Scoring of the plumage colour measurements for females of each species with double scoring species involved. Species with double scoring are marked in red and excluded from the second analysis.

| Latin name | cod  ing | Latin name | cod  ing | Latin name | cod  ing | Latin name | cod  ing | Latin name | cod  ing | Latin  name | cod  ing |
| --- | --- | --- | --- | --- | --- | --- | --- | --- | --- | --- | --- |
| Acanthidops bairdii | 1 | Dacnis nigripes | 3 | Incaspiza pulchra | 1 | Poospiza caesar | 1 | Sporophila albogularis | 0 | Tangara icteroceph  ala | 0 |
| Anisognathu  s igniventris | 3 | Dacnis  venusta | 3 | Iridophanes  pulcherrimus | 3 | Poospiza  cinerea | 2 | Sporophila  americana | 0 | Tangara  inornata | 3 |
| Anisognathu  s lacrymosus | 3 | Dacnis  viguieri | 0 | Iridosornis  analis | 3 | Poospiza  erythrophrys | 2 | Sporophila  bouvreuil | 0 | Tangara  johannae | 3 |
| Anisognathu s  melanogenys | 3 | Delothraupis castaneovent  ris | 2 | Iridosornis jelskii | 3 | Poospiza hispaniolensi  s | 0 | Sporophila bouvronides | 0 | Tangara labradorid  es | 3 |
| Anisognathu s  somptuosus | 3 | Diglossa albilatera | 0 | Iridosornis porphyrocep  halus | 3 | Poospiza hypochondri  a | 1 | Sporophila caerulescens | 0 | Tangara larvata | 3 |

| Bangsia arcaei | 2 | Diglossa baritula | 0 | Iridosornis rufivertex | 3 | Poospiza lateralis | 2 | Sporophila castaneivent  ris | 0 | Tangara lavinia | 3 |
| --- | --- | --- | --- | --- | --- | --- | --- | --- | --- | --- | --- |
| Bangsia edwardsi | 3 | Diglossa brunneiventri  s | 1 | Lanio aurantius | 1 | Poospiza melanoleuca | 1 | Sporophila cinnamomea | 0 | Tangara mexicana | 3 |
| Buthraupis eximia | 3 | Diglossa caerulescens | 2 | Lanio fulvus | 0 | Poospiza nigrorufa | 1 | Sporophila collaris | 0 | Tangara nigrocinct  a | 3 |
| Buthraupis montana | 3 | Diglossa carbonaria | 1 | Lanio leucothorax | 0 | Poospiza ornata | 1 | Sporophila corvina | 0 | Tangara nigroviridi  s | 3 |
| Calochaetes  coccineus | 0 | Diglossa  cyanea | 3 | Lanio  versicolor | 0 | Poospiza  thoracica | 1 | Sporophila  frontalis | 0 | Tangara  parzudakii | 3 |
| Camarhynch us pallidus | 0 | Diglossa duidae | 2 | Lophospingu s griseocristat  us | 1 | Poospiza torquata | 1 | Sporophila hypoxantha | 0 | Tangara peruviana | 0 |
| Camarhynch  us parvulus | 1 | Diglossa  glauca | 3 | Lophospingu  s pusillus | 1 | Poospiza  whitii | 2 | Sporophila  intermedia | 0 | Tangara  preciosa | 0 |
| Camarhynch us psittacula | 0 | Diglossa gloriosa | 1 | Loxigilla barbadensis | 1 | Porphyrospiz a  caerulescens | 1 | Sporophila leucoptera | 0 | Tangara punctata | 3 |
| Catamblyrhy nchus  diadema | 2 | Diglossa gloriosissima | 2 | Loxigilla noctis | 1 | Pyrrhocoma ruficeps | 0 | Sporophila lineola | 0 | Tangara ruficervix | 3 |
| Catamenia  analis | 2 | Diglossa  humeralis | 1 | Loxigilla  portoricensis | 0 | Ramphocelu  s bresilius | 0 | Sporophila  luctuosa | 0 | Tangara  rufigenis | 3 |
| Catamenia  homochroa | 1 | Diglossa  indigotica | 3 | Loxigilla  violacea | 0 | Ramphocelu  s carbo | 0 | Sporophila  minuta | 0 | Tangara  rufigula | 0 |
| Catamenia  inornata | 1 | Diglossa  lafresnayii | 2 | Loxipasser  anoxanthus | 1 | Ramphocelu  s dimidiatus | 0 | Sporophila  murallae | 0 | Tangara  schrankii | 3 |
| Certhidea olivacea | 0 | Diglossa major | 2 | Melanodera melanodera | 0 | Ramphocelu s  flammigerus | 0 | Sporophila nigricollis | 0 | Tangara seledon | 3 |
| Charitospiza eucosma | 1 | Diglossa mystacalis | 2 | Melanodera xanthogram ma | 0 | Ramphocelu s melanogaste  r | 0 | Sporophila nigrorufa | 0 | Tangara varia | 0 |
| Chlorochrysa calliparaea | 3 | Diglossa plumbea | 0 | Melanospiza richardsoni | 1 | Ramphocelu s  nigrogularis | 0 | Sporophila palustris | 0 | Tangara vassorii | 3 |
| Chlorochrysa  phoenicotis | 0 | Diglossa  sittoides | 0 | Melopyrrha  nigra | 0 | Ramphocelu  s passerinii | 1 | Sporophila  peruviana | 0 | Tangara  velia | 3 |
| Chlorophane s spiza | 0 | Diglossa venezuelensis | 0 | Nemosia pileata | 2 | Ramphocelu s sanguinolent  us | 0 | Sporophila plumbea | 0 | Tangara viridicollis | 1 |
| Chlorornis  riefferii | 0 | Diuca diuca | 1 | Neothraupis  fasciata | 1 | Rhodospingu  s cruentus | 0 | Sporophila  ruficollis | 0 | Tangara  vitriolina | 3 |
| Chrysothlypi s  chrysomelas | 0 | Donacospiza albifrons | 1 | Nesospiza acunhae | 0 | Rowettia goughensis | 0 | Sporophila simplex | 0 | Tangara xanthocep  hala | 3 |
| Chrysothlypi s salmoni | 0 | Dubusia taeniata | 3 | Nesospiza questi | 0 | Saltator albicollis | 0 | Sporophila telasco | 0 | Tangara xanthogast  ra | 0 |
| Cissopis  leverianus | 0 | Emberizoides  herbicola | 1 | Nesospiza  wilkinsi | 0 | Saltator  atriceps | 1 | Sporophila  torqueola | 0 | Tersina  viridis | 0 |
| Cnemoscopu s rubrirostris | 1 | Embernagra platensis | 2 | Orchesticus abeillei | 0 | Saltator atricollis | 0 | Stephanoph orus  diadematus | 3 | Thlypopsis fulviceps | 1 |
| Coereba  flaveola | 1 | Eucometis  penicillata | 2 | Oreomanes  fraseri | 2 | Saltator  atripennis | 1 | Tachyphonu  s coronatus | 1 | Thlypopsis  inornata | 1 |
| Compsospiza garleppi | 1 | Euneornis campestris | 2 | Oryzoborus angolensis | 0 | Saltator aurantiirostri  s | 1 | Tachyphonu s cristatus | 0 | Thlypopsis ornata | 1 |
| Conirostrum  albifrons | 3 | Geospiza  conirostris | 0 | Oryzoborus  crassirostris | 0 | Saltator  coerulescens | 1 | Tachyphonu  s delatrii | 0 | Thlypopsis  pectoralis | 1 |
| Conirostrum  bicolor | 3 | Geospiza  difficilis | 0 | Oryzoborus  maximiliani | 0 | Saltator  fuliginosus | 2 | Tachyphonu  s luctuosus | 1 | Thlypopsis  ruficeps | 0 |

| Conirostrum  cinereum | 2 | Geospiza  fortis | 0 | Oryzoborus  nuttingi | 0 | Saltator  grossus | 2 | Tachyphonu  s phoenicius | 1 | Thlypopsis  sordida | 1 |
| --- | --- | --- | --- | --- | --- | --- | --- | --- | --- | --- | --- |
| Conirostrum ferrugineive  ntre | 2 | Geospiza fuliginosa | 1 | Parkerthraus tes  humeralis | 2 | Saltator maxillosus | 1 | Tachyphonu s rufiventer | 1 | Thraupis abbas | 3 |
| Conirostrum leucogenys | 2 | Geospiza magnirostris | 1 | Paroaria capitata | 0 | Saltator maximus | 1 | Tachyphonu s rufus | 0 | Thraupis bonariensi  s | 3 |
| Conirostrum margaritae | 2 | Geospiza scandens | 0 | Paroaria coronata | 1 | Saltator nigriceps | 1 | Tachyphonu s surinamus | 1 | Thraupis cyanoceph  ala | 3 |
| Conirostrum rufum | 2 | Gubernatrix cristata | 0 | Paroaria dominicana | 1 | Saltator orenocensis | 1 | Tangara argyrofenge  s | 3 | Thraupis episcopus | 3 |
| Conirostrum sitticolor | 2 | Haplospiza unicolor | 0 | Paroaria gularis | 0 | Saltator similis | 1 | Tangara arthus | 0 | Thraupis glaucocolp  a | 3 |
| Conirostrum  speciosum | 2 | Hemispingus  atropileus | 0 | Phrygilus  alaudinus | 0 | Saltator  striatipectus | 1 | Tangara  callophrys | 3 | Thraupis  ornata | 3 |
| Conothraupi  s speculigera | 0 | Hemispingus  calophrys | 0 | Phrygilus  atriceps | 1 | Saltatricula  multicolor | 1 | Tangara  cayana | 3 | Thraupis  sayaca | 3 |
| Coryphaspiza  melanotis | 1 | Hemispingus  frontalis | 0 | Phrygilus  dorsalis | 2 | Schistochlam  ys melanopis | 1 | Tangara  chilensis | 3 | Tiaris  bicolor | 1 |
| Coryphospin gus  cucullatus | 0 | Hemispingus goeringi | 2 | Phrygilus erythronotus | 2 | Schistochlam ys  ruficapillus | 2 | Tangara chrysotis | 0 | Tiaris canorus | 1 |
| Coryphospin  gus pileatus | 1 | Hemispingus  melanotis | 1 | Phrygilus  fruticeti | 1 | Sericossypha  albocristata | 0 | Tangara  cucullata | 3 | Tiaris  fuliginosus | 0 |
| Creurgops  dentatus | 2 | Hemispingus  reyi | 1 | Phrygilus  gayi | 2 | Sicalis  auriventris | 0 | Tangara  cyanicollis | 3 | Tiaris  obscurus | 1 |
| Creurgops verticalis | 1 | Hemispingus superciliaris | 0 | Phrygilus patagonicus | 2 | Sicalis citrina | 0 | Tangara cyanocephal  a | 3 | Tiaris olivaceus | 0 |
| Cyanerpes caeruleus | 3 | Hemispingus verticalis | 1 | Phrygilus plebejus | 1 | Sicalis columbiana | 0 | Tangara cyanoptera | 3 | Trichothra upis  melanops | 0 |
| Cyanerpes cyaneus | 0 | Hemispingus xanthophthal  mus | 1 | Phrygilus punensis | 2 | Sicalis flaveola | 0 | Tangara cyanotis | 3 | Urothraupi s  stolzmanni | 1 |
| Cyanerpes  lucidus | 3 | Hemithraupis  flavicollis | 0 | Phrygilus  unicolor | 0 | Sicalis  lebruni | 1 | Tangara  cyanoventris | 3 | Volatinia  jacarina | 0 |
| Cyanerpes  nitidus | 3 | Hemithraupis  guira | 0 | Piezorhina  cinerea | 1 | Sicalis lutea | 0 | Tangara  desmaresti | 0 | Xenodacni  s parina | 3 |
| Cyanicterus cyanicterus | 3 | Hemithraupis ruficapilla | 0 | Pinaroloxias inornata | 0 | Sicalis luteocephala | 1 | Tangara dowii | 3 | Xenosping us  concolor | 2 |
| Cypsnagra  hirundinacea | 0 | Heterospingu  s rubrifrons | 1 | Pipraeidea  melanonota | 3 | Sicalis  luteola | 0 | Tangara  fastuosa | 3 |  |  |
| Dacnis cayana | 3 | Heterospingu s  xanthopygius | 1 | Platyspiza crassirostris | 0 | Sicalis olivascens | 1 | Tangara florida | 0 |  |  |
| Dacnis  flaviventer | 0 | Idiopsar  brachyurus | 1 | Poospiza  alticola | 1 | Sicalis  raimondii | 1 | Tangara  guttata | 3 |  |  |
| Dacnis  hartlaubi | 1 | Incaspiza  laeta | 1 | Poospiza  boliviana | 1 | Sicalis  taczanowskii | 1 | Tangara  gyrola | 3 |  |  |
| Dacnis  lineata | 1 | Incaspiza  personata | 1 | Poospiza  cabanisi | 1 | Sicalis  uropygialis | 1 | Tangara  heinei | 3 |  |  |

**Table S4.** Reverse Jump analysis results for male; analysis done with the table containing double scored species

|  | transition rate | colour transitions | ESS | Me an | Me dia  n | Mo de | 95% HPD | % Zero | number of zero | total number | % not zero |
| --- | --- | --- | --- | --- | --- | --- | --- | --- | --- | --- | --- |
| male_ 1 | q01 | any other colour -> grey | 6579  1.80  0 | 0.0  44 | 0.0  43 | 0.0  42 | [0.029,  0.0604] | 0.000 | 1.000 | 218000.00  0 | 100.000 |
|  | q02 | any other | 1450 | 0.0 | 0.0 | 0.0 | [0, | 6.552 | 14284.00 | 218000.00 | 93.448 |

|  |  | colour -> slate | 5.70  0 | 41 | 42 | 00 | 0.0573] |  | 0 | 0 |  |
| --- | --- | --- | --- | --- | --- | --- | --- | --- | --- | --- | --- |
|  | q03 | any other colour -> blue | 2845  3.50  0 | 0.0  00 | 0.0  00 | 0.0  00 | [0, 0] | 99.104 | 216047.0  00 | 218000.00  0 | 0.896 |
|  | q10 | grey -> any other colour | 6634  7.30  0 | 0.0  44 | 0.0  43 | 0.0  42 | [0.029,  0.0605] | 0.000 | 0.000 | 218000.00  0 | 100.000 |
|  | q12 | grey -> slate | 6510  6.50  0 | 0.0  44 | 0.0  43 | 0.0  42 | [0.0291,  0.0604] | 0.000 | 0.000 | 218000.00  0 | 100.000 |
|  | q13 | grey -> blue | 1265  47.4  00 | 0.0  00 | 0.0  00 | 0.0  00 | [0, 0] | 99.871 | 217718.0  00 | 218000.00  0 | 0.129 |
|  | q20 | slate -> any other colour | 4839  7.30  0 | 0.0  30 | 0.0  38 | 0.0  00 | [0,  0.0561] | 29.555 | 64430.00  0 | 218000.00  0 | 70.445 |
|  | q21 | slate -> grey | 6652  4.90  0 | 0.0  44 | 0.0  43 | 0.0  42 | [0.029,  0604] | 0.002 | 4.000 | 218000.00  0 | 99.998 |
|  | q23 | slate -> blue | 7593  0.00  0 | 0.0  44 | 0.0  43 | 0.0  42 | [0.0293,  0.0602] | 0.000 | 0.000 | 218000.00  0 | 100.000 |
|  | q30 | blue -> any other colour | 7589  4.20  0 | 0.0  44 | 0.0  43 | 0.0  42 | [0.0293,  0.0602] | 0.000 | 0.000 | 218000.00  0 | 100.000 |
|  | q31 | blue -> grey | 9985  9.70  0 | 0.0  00 | 0.0  00 | 0.0  00 | [0,  0.0578] | 99.970 | 217934.0  00 | 218000.00  0 | 0.030 |
|  | q32 | blue -> slate | 3328  .400 | 0.0  01 | 0.0  00 | 0.0  00 | [0,  0.0818] | 98.111 | 213882.0  00 | 218000.00  0 | 1.889 |
| male_ 2 | q01 | any other colour -> grey | 7261  6.50  0 | 0.0  44 | 0.0  43 | 0.0  38 | [0.029,  0.0604] | 0.002 | 4.000 | 218000.00  0 | 99.998 |
|  | q02 | any other colour -> slate | 1060  9.00  0 | 0.0  40 | 0.0  42 | 0.0  00 | [0,  0.0573] | 6.806 | 14837.00  0 | 218000.00  0 | 93.194 |
|  | q03 | any other colour -> blue | 2886  5.70  0 | 0.0  00 | 0.0  00 | 0.0  00 | [0, 0] | 99.106 | 216051.0  00 | 218000.00  0 | 0.894 |
|  | q10 | grey -> any other colour | 7254  5.70  0 | 0.0  44 | 0.0  43 | 0.0  38 | [0.029,  0.0604] | 0.000 | 0.000 | 218000.00  0 | 100.000 |
|  | q12 | grey -> slate | 6619  4.80  0 | 0.0  44 | 0.0  43 | 0.0  38 | [0.0292,  0.0604] | 0.000 | 0.000 | 218000.00  0 | 100.000 |
|  | q13 | grey -> blue | 1346  40.8  00 | 0.0  00 | 0.0  00 | 0.0  00 | [0, 0] | 99.874 | 217725.0  00 | 218000.00  0 | 0.126 |
|  | q20 | slate -> any other colour | 3813  3.30  0 | 0.0  30 | 0.0  38 | 0.0  00 | [0,  0.0561] | 29.513 | 64338.00  0 | 218000.00  0 | 70.487 |
|  | q21 | slate -> grey | 7260  4.40  0 | 0.0  44 | 0.0  43 | 0.0  38 | [0.029,  0.0604] | 0.003 | 6.000 | 218000.00  0 | 99.997 |
|  | q23 | slate -> blue | 7748  1.60  0 | 0.0  44 | 0.0  43 | 0.0  38 | [0.0294,  0.0602] | 0.001 | 3.000 | 218000.00  0 | 99.999 |
|  | q30 | blue -> any other colour | 7657  5.00  0 | 0.0  44 | 0.0  43 | 0.0  38 | [0.0294,  0.0602] | 0.000 | 0.000 | 218000.00  0 | 100.000 |
|  | q31 | blue -> grey | 1404  47.3  00 | 0.0  00 | 0.0  00 | 0.0  00 | [0, 0] | 99.981 | 217959.0  00 | 218000.00  0 | 0.019 |
|  | q32 | blue -> slate | 2530  .400 | 0.0  01 | 0.0  00 | 0.0  00 | [0, 0] | 97.830 | 213270.0  00 | 218000.00  0 | 2.170 |
| male_ 3 | q01 | any other colour -> grey | 6807  6.50  0 | 0.0  44 | 0.0  43 | 0.0  43 | [0.0291,  0.0607] | 0.002 | 4.000 | 218000.00  0 | 99.998 |

|  | q02 | any other colour -> slate | 1245  9.70  0 | 0.0  40 | 0.0  42 | 0.0  00 | [0,  0.0574] | 6.757 | 14730.00  0 | 218000.00  0 | 93.243 |
| --- | --- | --- | --- | --- | --- | --- | --- | --- | --- | --- | --- |
|  | q03 | any other colour -> blue | 2523  5.50  0 | 0.0  00 | 0.0  00 | 0.0  00 | [0, 0] | 99.094 | 216026.0  00 | 218000.00  0 | 0.906 |
|  | q10 | grey -> any other colour | 6826  2.60  0 | 0.0  44 | 0.0  43 | 0.0  43 | [0.0291,  0.0607] | 0.000 | 0.000 | 218000.00  0 | 100.000 |
|  | q12 | grey -> slate | 6675  6.90  0 | 0.0  44 | 0.0  43 | 0.0  43 | [0.0291,  0.0605] | 0.000 | 0.000 | 218000.00  0 | 100.000 |
|  | q13 | grey -> blue | 9073  2.20  0 | 0.0  00 | 0.0  00 | 0.0  00 | [0, 0] | 99.834 | 217638.0  00 | 218000.00  0 | 0.166 |
|  | q20 | slate -> any other colour | 4473  0.80  0 | 0.0  30 | 0.0  38 | 0.0  00 | [0,  0.0561] | 29.667 | 64675.00  0 | 218000.00  0 | 70.333 |
|  | q21 | slate -> grey | 6837  5.80  0 | 0.0  44 | 0.0  43 | 0.0  43 | [0.0291,  0.0607] | 0.003 | 6.000 | 218000.00  0 | 99.997 |
|  | q23 | slate -> blue | 7633  4.10  0 | 0.0  44 | 0.0  43 | 0.0  43 | [0.0291,  0.0601] | 0.005 | 10.000 | 218000.00  0 | 99.995 |
|  | q30 | blue -> any other colour | 7624  7.60  0 | 0.0  44 | 0.0  43 | 0.0  43 | [0.0291,  0.0601] | 0.000 | 0.000 | 218000.00  0 | 100.000 |
|  | q31 | blue -> grey | 1582  34.7  00 | 0.0  00 | 0.0  00 | 0.0  00 | [0, 0] | 99.978 | 217951.0  00 | 218000.00  0 | 0.022 |
|  | q32 | blue -> slate | 2428  .400 | 0.0  01 | 0.0  00 | 0.0  00 | [0, 0] | 97.913 | 213451.0  00 | 218000.00  0 | 2.087 |

**Table S5.** Reverse Jump analysis results for female; analysis done with the table containing double scores species

|  | transition  rate | colour  transitions | ESS | Mea  n | Med  ian | Mod  e | 95% HPD | %  Zero | number  of zero | total  number | % not  zero |
| --- | --- | --- | --- | --- | --- | --- | --- | --- | --- | --- | --- |
| female_ | q01 | any other colour | 1033 | 0.06 | 0.06 | 0.06 | [0.048, | 0.00 | 0.000 | 218000. | 100.000 |
| 1 |  | -> grey | 42.20 | 5 | 4 | 4 | 0.0807] | 0 |  | 000 |  |
|  |  |  | 0 |  |  |  |  |  |  |  |  |
|  | q02 | any other colour | 1343 | 0.00 | 0.00 | 0.00 | [0, 0] | 96.6 | 210758.0 | 218000. | 3.322 |
|  |  | -> slate | 60.50 | 2 | 0 | 0 |  | 78 | 00 | 000 |  |
|  |  |  | 0 |  |  |  |  |  |  |  |  |
|  | q03 | any other colour | 8683 | 0.00 | 0.00 | 0.00 | [0, 0] | 99.9 | 217908.0 | 218000. | 0.042 |
|  |  | -> blue | 9.800 | 0 | 0 | 0 |  | 58 | 00 | 000 |  |
|  | q10 | grey -> any other | 1025 | 0.06 | 0.06 | 0.06 | [0.0481, | 0.00 | 0.000 | 218000. | 100.000 |
|  |  | colour | 56.30 | 5 | 4 | 4 | 0.0807] | 0 |  | 000 |  |
|  |  |  | 0 |  |  |  |  |  |  |  |  |
|  | q12 | grey -> slate | 1440 | 0.06 | 0.06 | 0.06 | [0.0484, | 0.00 | 2.000 | 218000. | 99.999 |
|  |  |  | 16.00 | 4 | 4 | 4 | 0.0807] | 1 |  | 000 |  |
|  |  |  | 0 |  |  |  |  |  |  |  |  |
|  | q13 | grey -> blue | 9287 | 0.00 | 0.00 | 0.00 | [0, 0] | 99.9 | 217991.0 | 218000. | 0.004 |
|  |  |  | 0.600 | 0 | 0 | 0 |  | 96 | 00 | 000 |  |
|  | q20 | slate -> any | 7110 | 0.06 | 0.06 | 0.00 | [0.0479, | 0.53 | 1162.000 | 218000. | 99.467 |
|  |  | other colour | 5.400 | 4 | 4 | 0 | 0.0807] | 3 |  | 000 |  |
|  | q21 | slate -> grey | 1056 | 0.06 | 0.06 | 0.00 | [0.048, | 0.05 | 128.000 | 218000. | 99.941 |
|  |  |  | 26.00 | 4 | 4 | 0 | 0.0807] | 9 |  | 000 |  |
|  |  |  | 0 |  |  |  |  |  |  |  |  |
|  | q23 | slate -> blue | 1370 | 0.06 | 0.06 | 0.06 | [0.0484, | 0.00 | 1.000 | 218000. | 100.000 |
|  |  |  | 94.20 | 4 | 4 | 4 | 0.0807] | 0 |  | 000 |  |
|  |  |  | 0 |  |  |  |  |  |  |  |  |
|  | q30 | blue -> any other | 1355 | 0.06 | 0.06 | 0.06 | [0.0484, | 0.00 | 0.000 | 218000. | 100.000 |
|  |  | colour | 17.90 | 4 | 4 | 4 | 0.0807] | 0 |  | 000 |  |
|  |  |  | 0 |  |  |  |  |  |  |  |  |

|  | q31 | blue -> grey | 1849  4.200 | 0.00  0 | 0.00  0 | 0.00  0 | [0, 0] | 99.8  93 | 217767.0  00 | 218000.  000 | 0.107 |
| --- | --- | --- | --- | --- | --- | --- | --- | --- | --- | --- | --- |
|  | q32 | blue -> slate | 1652  9.100 | 0.00  0 | 0.00  0 | 0.00  0 | [0, 0] | 99.9  13 | 217811.0  00 | 218000.  000 | 0.087 |
| female_ | q01 | any other colour | 1041 | 0.06 | 0.06 | 0.06 | [0.0482, | 0.00 | 0.000 | 218000. | 100.000 |
| 2 |  | -> grey | 50.60 | 5 | 4 | 4 | 0.0808] | 0 |  | 000 |  |
|  |  |  | 0 |  |  |  |  |  |  |  |  |
|  | q02 | any other colour | 1319 | 0.00 | 0.00 | 0.00 | [0, 0] | 96.6 | 210623.0 | 218000. | 3.384 |
|  |  | -> slate | 91.20 | 2 | 0 | 0 |  | 16 | 00 | 000 |  |
|  |  |  | 0 |  |  |  |  |  |  |  |  |
|  | q03 | any other colour | 6737 | 0.00 | 0.00 | 0.00 | [0, 0] | 99.9 | 217940.0 | 218000. | 0.028 |
|  |  | -> blue | 1.700 | 0 | 0 | 0 |  | 72 | 00 | 000 |  |
|  | q10 | grey -> any other | 1037 | 0.06 | 0.06 | 0.06 | [0.0482, | 0.00 | 0.000 | 218000. | 100.000 |
|  |  | colour | 34.70 | 5 | 4 | 4 | 0.0807] | 0 |  | 000 |  |
|  |  |  | 0 |  |  |  |  |  |  |  |  |
|  | q12 | grey -> slate | 1396 | 0.06 | 0.06 | 0.06 | [0.0481, | 0.00 | 6.000 | 218000. | 99.997 |
|  |  |  | 47.80 | 4 | 4 | 4 | 0.0804] | 3 |  | 000 |  |
|  |  |  | 0 |  |  |  |  |  |  |  |  |
|  | q13 | grey -> blue | 1533 | 0.00 | 0.00 | 0.00 | [0, 0] | 99.9 | 217994.0 | 218000. | 0.003 |
|  |  |  | 73.50 | 0 | 0 | 0 |  | 97 | 00 | 000 |  |
|  |  |  | 0 |  |  |  |  |  |  |  |  |
|  | q20 | slate -> any | 5874 | 0.06 | 0.06 | 0.00 | [0.0481, | 0.47 | 1026.000 | 218000. | 99.529 |
|  |  | other colour | 2.400 | 4 | 4 | 0 | 0.0808] | 1 |  | 000 |  |
|  | q21 | slate -> grey | 1046 | 0.06 | 0.06 | 0.00 | [0.0482, | 0.07 | 158.000 | 218000. | 99.928 |
|  |  |  | 05.60 | 4 | 4 | 0 | 0.0808] | 2 |  | 000 |  |
|  |  |  | 0 |  |  |  |  |  |  |  |  |
|  | q23 | slate -> blue | 1387 | 0.06 | 0.06 | 0.06 | [0.0481, | 0.00 | 0.000 | 218000. | 100.000 |
|  |  |  | 04.70 | 4 | 4 | 4 | 0.0804] | 0 |  | 000 |  |
|  |  |  | 0 |  |  |  |  |  |  |  |  |
|  | q30 | blue -> any other | 1402 | 0.06 | 0.06 | 0.06 | [0.0482, | 0.00 | 0.000 | 218000. | 100.000 |
|  |  | colour | 20.60 | 4 | 4 | 4 | 0.0804] | 0 |  | 000 |  |
|  |  |  | 0 |  |  |  |  |  |  |  |  |
|  | q31 | blue -> grey | 8026. | 0.00 | 0.00 | 0.00 | [0, 0] | 99.9 | 217834.0 | 218000. | 0.076 |
|  |  |  | 400 | 0 | 0 | 0 |  | 24 | 00 | 000 |  |
|  | q32 | blue -> slate | 1655 | 0.00 | 0.00 | 0.00 | [0, 0] | 99.9 | 217835.0 | 218000. | 0.076 |
|  |  |  | 4.800 | 0 | 0 | 0 |  | 24 | 00 | 000 |  |
| female_ | q01 | any other colour | 9941 | 0.06 | 0.06 | 0.06 | [0.048, | 0.00 | 0.000 | 218000. | 100.000 |
| 3 |  | -> grey | 7.200 | 5 | 4 | 4 | 0.0808] | 0 |  | 000 |  |
|  | q02 | any other colour | 1313 | 0.00 | 0.00 | 0.00 | [0, 0] | 96.6 | 210695.0 | 218000. | 3.351 |
|  |  | -> slate | 06.20 | 2 | 0 | 0 |  | 49 | 00 | 000 |  |
|  |  |  | 0 |  |  |  |  |  |  |  |  |
|  | q03 | any other colour | 9443 | 0.00 | 0.00 | 0.00 | [0, 0] | 99.9 | 217906.0 | 218000. | 0.043 |
|  |  | -> blue | 2.100 | 0 | 0 | 0 |  | 57 | 00 | 000 |  |
|  | q10 | grey -> any other | 9870 | 0.06 | 0.06 | 0.06 | [0.0479, | 0.00 | 1.000 | 218000. | 100.000 |
|  |  | colour | 3.800 | 5 | 4 | 4 | 0.0806] | 0 |  | 000 |  |
|  | q12 | grey -> slate | 1437 | 0.06 | 639. | 0.06 | [0.0481, | 0.00 | 1.000 | 218000. | 100.000 |
|  |  |  | 92.40 | 4 | 000 | 4 | 0.0805] | 0 |  | 000 |  |
|  |  |  | 0 |  |  |  |  |  |  |  |  |
|  | q13 | grey -> blue | 1718 | 0.00 | 0.00 | 0.00 | [0, 0] | 99.9 | 217981.0 | 218000. | 0.009 |
|  |  |  | 2.100 | 0 | 0 | 0 |  | 91 | 00 | 000 |  |
|  | q20 | slate -> any | 6321 | 0.06 | 0.06 | 0.00 | [0.048, | 0.48 | 1064.000 | 218000. | 99.512 |
|  |  | other colour | 0.200 | 4 | 4 | 0 | 0.0808] | 8 |  | 000 |  |
|  | q21 | slate -> grey | 1016 | 0.06 | 0.06 | 0.00 | [0.0479, | 0.06 | 132.000 | 218000. | 99.939 |
|  |  |  | 12.80 | 4 | 4 | 0 | 0.0806] | 1 |  | 000 |  |
|  |  |  | 0 |  |  |  |  |  |  |  |  |
|  | q23 | slate -> blue | 1339 | 0.06 | 0.06 | 0.06 | [0.0479, | 0.00 | 11.000 | 218000. | 99.995 |
|  |  |  | 40.10 | 4 | 4 | 4 | 0.0803] | 5 |  | 000 |  |
|  |  |  | 0 |  |  |  |  |  |  |  |  |
|  | q30 | blue -> any other | 1380 | 0.06 | 0.06 | 0.06 | [0.0479, | 0.00 | 0.000 | 218000. | 100.000 |
|  |  | colour | 57.80 | 4 | 4 | 4 | 0.0803] | 0 |  | 000 |  |
|  |  |  | 0 |  |  |  |  |  |  |  |  |
|  | q31 | blue -> grey | 1282 | 0.00 | 0.00 | 0.00 | [0, 0] | 99.9 | 217809.0 | 218000. | 0.088 |
|  |  |  | 2.800 | 0 | 0 | 0 |  | 12 | 00 | 000 |  |
|  | q32 | blue -> slate | 1569 | 0.00 | 0.00 | 0.00 | [0,0] | 99.9 | 217843.0 | 218000. | 0.072 |
|  |  |  | 5.900 | 0 | 0 | 0 |  | 28 | 00 | 000 |  |

**Table S6**. Reverse Jump analysis results for male; analysis done with the table not containing double scored species

|  | trans ition  rate | colour transitions | ESS | Mea n | Medi an | Mod e | 95% HPD | %  Zero | number of zero | total number | % not zero |
| --- | --- | --- | --- | --- | --- | --- | --- | --- | --- | --- | --- |
| male | q01 | any other colour - | 55006 | 0.040 | 0.03 | 0.03 | [0.0271, | 0.00 | 0.000 | 218000.000 | 100.000 |
| _1 |  | > grey | .000 |  | 9 | 7 | 0.0544] | 0 |  |  |  |
|  | q02 | any other colour - | 86926 | 0.033 | 0.03 | 0.00 | [0, | 98.3 | 214362.000 | 218000.000 | 1.669 |
|  |  | > slate | .000 |  | 7 | 0 | 0.0496] | 31 |  |  |  |
|  | q03 | any other colour - | 43413 | 0.000 | 0.00 | 0.00 | [0, 0] | 99.9 | 217861.000 | 218000.000 | 0.064 |
|  |  | > blue | .000 |  | 0 | 0 |  | 36 |  |  |  |
|  | q10 | grey -> any other | 55016 | 0.040 | 0.03 | 0.03 | [0.0272, | 0.00 | 0.000 | 218000.000 | 100.000 |
|  |  | colour | .000 |  | 9 | 7 | 0.0545] | 0 |  |  |  |
|  | q12 | grey -> slate | 55446 | 0.040 | 0.03 | 0.03 | [0.0272, | 0.00 | 6.000 | 218000.000 | 99.997 |
|  |  |  | .000 |  | 9 | 7 | 0.0544] | 3 |  |  |  |
|  | q13 | grey -> blue | 50429 | 0.000 | 0.00 | 0.00 | [0, 0] | 99.8 | 217688.000 | 218000.000 | 0.143 |
|  |  |  | .000 |  | 0 | 0 |  | 57 |  |  |  |
|  | q20 | slate -> any other | 10190 | 0.032 | 0.03 | 0.00 | [0, | 6.45 | 14073.000 | 218000.000 | 93.544 |
|  |  | colour | 0.000 |  | 7 | 0 | 0.0502] | 6 |  |  |  |
|  | q21 | slate -> grey | 55579 | 0.040 | 0.03 | 0.03 | [0.0272, | 1.29 | 2819.000 | 218000.000 | 98.707 |
|  |  |  | .000 |  | 9 | 7 | 0.0544] | 3 |  |  |  |
|  | q23 | slate -> blue | 86990 | 0.040 | 0.03 | 0.00 | [0.0277, | 5.42 | 11827.000 | 218000.000 | 94.575 |
|  |  |  | .000 |  | 9 | 0 | 0.054] | 5 |  |  |  |
|  | q30 | blue -> any other | 92878 | 9287 | 0.03 | 0.03 | [0.0272, | 0.00 | 20.000 | 218000.000 | 99.991 |
|  |  | colour | .000 | 8.000 | 9 | 7 | 0.0534] | 9 |  |  |  |
|  | q31 | blue -> grey | 15410 | 0.000 | 0.00 | 0.00 | [0, 0] | 94.0 | 205101.000 | 218000.000 | 5.917 |
|  |  |  | 0.000 |  | 0 | 0 |  | 83 |  |  |  |
|  | q32 | blue -> slate | 20358 | 0.000 | 0.00 | 0.00 | [0, 0] | 98.7 | 215246.000 | 218000.000 | 1.263 |
|  |  |  | .000 |  | 0 | 0 |  | 37 |  |  |  |
| male | q01 | any other colour - | 54079 | 0.040 | 0.03 | 0.03 | [0.0274, | 0.00 | 6.000 | 218000.000 | 99.997 |
| _2 |  | > grey | .000 |  | 9 | 8 | 0.0547] | 3 |  |  |  |
|  | q02 | any other colour - | 88908 | 0.033 | 0.03 | 0.00 | [0, | 14.9 | 32523.000 | 218000.000 | 85.081 |
|  |  | > slate | .000 |  | 7 | 0 | 0.0496] | 19 |  |  |  |
|  | q03 | any other colour - | 45276 | 0.000 | 0.00 | 0.00 | [0, 0] | 99.4 | 216816.000 | 218000.000 | 0.543 |
|  |  | > blue | .000 |  | 0 | 0 |  | 57 |  |  |  |
|  | q10 | grey -> any other | 54182 | 0.040 | 0.03 | 0.03 | [0.0274, | 0.00 | 0.000 | 218000.000 | 100.000 |
|  |  | colour | .000 |  | 9 | 8 | 0.0547] | 0 |  |  |  |
|  | q12 | grey -> slate | 54228 | 0.040 | 0.03 | 0.03 | [0.0274, | 0.00 | 0.000 | 218000.000 | 100.000 |
|  |  |  | .000 |  | 9 | 8 | 0.0547] | 0 |  |  |  |
|  | q13 | grey -> blue | 29370 | 0.000 | 0.00 | 0.00 | [0, 0] | 99.8 | 217572.000 | 218000.000 | 0.196 |
|  |  |  | .000 |  | 0 | 0 |  | 04 |  |  |  |
|  | q20 | slate -> any other | 10270 | 0.032 | 0.03 | 0.00 | [0, | 17.8 | 38847.000 | 218000.000 | 82.180 |
|  |  | colour | 0.000 |  | 7 | 0 | 0/0504] | 20 |  |  |  |
|  | q21 | slate -> grey | 55315 | 0.040 | 0.03 | 0.03 | [0.0274, | 0.00 | 6.000 | 218000.000 | 99.997 |
|  |  |  | .000 |  | 9 | 8 | 0.0547] | 3 |  |  |  |
|  | q23 | slate -> blue | 86564 | 0.040 | 0.03 | 0.00 | [0.0275, | 0.03 | 78.000 | 218000.000 | 99.964 |
|  |  |  | .000 |  | 9 | 0 | 0.0538] | 6 |  |  |  |
|  | q30 | blue -> any other | 96608 | 9460 | 0.03 | 0.03 | [0.0275, | 0.00 | 0.000 | 218000.000 | 100.000 |
|  |  | colour | .000 | 8.000 | 9 | 8 | 0.0538] | 0 |  |  |  |
|  | q31 | blue -> grey | 88186 | 0.000 | 0.00 | 0.00 | [0, 0] | 99.9 | 217925.000 | 218000.000 | 0.034 |
|  |  |  | .000 |  | 0 | 0 |  | 66 |  |  |  |
|  | q32 | blue -> slate | 21723 | 0.000 | 0.00 | 0.00 | [0, 0] | 99.4 | 216840.000 | 218000.000 | 0.532 |
|  |  |  | .000 |  | 0 | 0 |  | 68 |  |  |  |
| male | q01 | any other colour - | 57759 | 0.404 | 0.03 | 0.03 | [0.0273, | 0.00 | 2.000 | 218000.000 | 99.999 |
| _3 |  | > grey | .000 |  | 9 | 8 | 0.0545] | 1 |  |  |  |
|  | q02 | any other colour - | 94523 | 0.033 | 0.03 | 0.00 | [0, | 14.8 | 32293.000 | 218000.000 | 85.187 |
|  |  | > slate | .000 |  | 7 | 0 | 0.0498] | 13 |  |  |  |
|  | q03 | any other colour - | 39017 | 0.000 | 0.00 | 0.00 | [0, 0] | 99.4 | 216863.000 | 218000.000 | 0.522 |
|  |  | > blue | .000 |  | 0 | 0 |  | 78 |  |  |  |
|  | q10 | grey -> any other | 57687 | 0.404 | 0.03 | 0.03 | [0.0273, | 0.00 | 2.000 | 218000.000 | 99.999 |
|  |  | colour | .000 |  | 9 | 8 | 0.0545] | 1 |  |  |  |
|  | q12 | grey -> slate | 61338 | 0.040 | 0.03 | 0.03 | [0.0274, | 0.00 | 1.000 | 218000.000 | 100.000 |
|  |  |  | .000 |  | 9 | 8 | 0.0545] | 0 |  |  |  |
|  | q13 | grey -> blue | 45878 | 0.000 | 0.00 | 0.00 | [0, 0] | 99.8 | 217684.000 | 218000.000 | 0.145 |
|  |  |  | .000 |  | 0 | 0 |  | 55 |  |  |  |
|  | q20 | slate -> any other | 11390 | 0.032 | 0.03 | 0.00 | [0, | 17.6 | 38483.000 | 218000.000 | 82.347 |

|  |  | colour | 0.000 |  | 7 | 0 | 0.0504] | 53 |  |  |  |
| --- | --- | --- | --- | --- | --- | --- | --- | --- | --- | --- | --- |
|  | q21 | slate -> grey | 61674  .000 | 0.040 | 0.03  9 | 0.03  8 | [0.0273,  0.0545] | 0.00  2 | 5.000 | 218000.000 | 99.998 |
|  | q23 | slate -> blue | 89305  .000 | 0.040 | 0.03  9 | 0.00  0 | [0.0276,  0.0539] | 0.02  4 | 53.000 | 218000.000 | 99.976 |
|  | q30 | blue -> any other  colour | 93404  .000 | 0.040 | 0.03  9 | 0.03  8 | [0.0276,  0.0539] | 0.00  0 | 0.000 | 218000.000 | 100.000 |
|  | q31 | blue -> grey | 77073  .000 | 0.000 | 0.00  0 | 0.00  0 | [0, 0] | 99.9  60 | 217913.000 | 218000.000 | 0.040 |
|  | q32 | blue -> slate | 25317  .000 | 0.000 | 0.00  0 | 0.00  0 | [0, 0] | 99.5  73 | 217069.000 | 218000.000 | 0.427 |

**Table S7.** Reverse Jump analysis results for female; analysis done with the table not containing double scores species

|  | transition  rate | colour  transitions | ESS | Mea  n | Med  ian | Mod  e | 95% HPD | %  Zero | number of  zero | total  number | % not zero |
| --- | --- | --- | --- | --- | --- | --- | --- | --- | --- | --- | --- |
| fem | q01 | any other | 1716 | 0.05 | 0.05 | 0.05 | [0.0383, | 0.00 | 0.000 | 218000.00 | 100.000 |
| ale_ |  | colour -> grey | 3.000 | 8 | 7 | 1 | 0.078] | 0 |  | 0 |  |
| 1 | q02 | any other | 1456 | 0.00 | 0.00 | 0.00 | [0, | 98.3 | 214362.00 | 218000.00 | 1.669 |
|  |  | colour -> slate | 00.00 | 1 | 0 | 0 | 0.0889] | 31 | 0 | 0 |  |
|  |  |  | 0 |  |  |  |  |  |  |  |  |
|  | q03 | any other | 8061 | 0.00 | 0.00 | 0.00 | [0, 0] | 99.9 | 217861.00 | 218000.00 | 0.064 |
|  |  | colour -> blue | 3.000 | 0 | 0 | 0 |  | 36 | 0 | 0 |  |
|  | q10 | grey -> any | 1741 | 0.05 | 0.05 | 0.05 | [0.0383, | 0.00 | 0.000 | 218000.00 | 100.000 |
|  |  | other colour | 1.000 | 8 | 7 | 1 | 0.078] | 0 |  | 0 |  |
|  | q12 | grey -> slate | 1466 | 0.05 | 0.05 | 0.05 | [0.0383, | 0.00 | 6.000 | 218000.00 | 99.997 |
|  |  |  | 9.000 | 8 | 7 | 1 | 0.078] | 3 |  | 0 |  |
|  | q13 | grey -> blue | 4568 | 0.00 | 0.00 | 0.00 | [0, 0] | 99.8 | 217688.00 | 218000.00 | 0.143 |
|  |  |  | 5.000 | 0 | 0 | 0 |  | 57 | 0 | 0 |  |
|  | q20 | slate -> any | 671.0 | 0.05 | 0.05 | 0.00 | [0, | 6.45 | 14073.000 | 218000.00 | 93.544 |
|  |  | other colour | 00 | 5 | 7 | 0 | 0.0754] | 6 |  | 0 |  |
|  | q21 | slate -> grey | 4044. | 0.05 | 0.05 | 0.00 | [0.0369, | 1.29 | 2819.000 | 218000.00 | 98.707 |
|  |  |  | 000 | 7 | 7 | 0 | 0.0795] | 3 |  | 0 |  |
|  | q23 | slate -> blue | 801.0 | 0.05 | 0.05 | 0.00 | [0, | 5.42 | 11827.000 | 218000.00 | 94.575 |
|  |  |  | 00 | 5 | 7 | 0 | 0.0754] | 5 |  | 0 |  |
|  | q30 | blue -> any | 1392 | 0.05 | 0.05 | 0.05 | [0.0382, | 0.00 | 20.000 | 218000.00 | 99.991 |
|  |  | other colour | 9.000 | 8 | 7 | 1 | 0.078] | 9 |  | 0 |  |
|  | q31 | blue -> grey | 505.0 | 0.00 | 0.00 | 0.00 | [0, | 94.0 | 205101.00 | 218000.00 | 5.917 |
|  |  |  | 00 | 3 | 0 | 0 | 0.0412] | 83 | 0 | 0 |  |
|  | q32 | blue -> slate | 1493. | 0.00 | 0.00 | 0.00 | [0, 0] | 98.7 | 215246.00 | 218000.00 | 1.263 |
|  |  |  | 000 | 1 | 0 | 0 |  | 37 | 0 | 0 |  |
| fem | q01 | any other | 1048 | 0.05 | 0.05 | 0.05 | [0.0382, | 0.00 | 0.000 | 218000.00 | 100.000 |
| ale_ |  | colour -> grey | 5.000 | 8 | 7 | 4 | 0.0778] | 0 |  | 0 |  |
| 2 | q02 | any other | 6572 | 0.00 | 0.00 | 0.00 | [0, 0] | 98.3 | 214480.00 | 218000.00 | 1.615 |
|  |  | colour -> slate | 4.000 | 1 | 0 | 0 |  | 85 | 0 | 0 |  |
|  | q03 | any other | 3950 | 0.00 | 0.00 | 0.00 | [0, 0] | 99.8 | 217774.00 | 218000.00 | 0.104 |
|  |  | colour -> blue | 2.000 | 0 | 0 | 0 |  | 96 | 0 | 0 |  |
|  | q10 | grey -> any | 1059 | 0.05 | 0.05 | 0.05 | [0.0382, | 0.00 | 0.000 | 218000.00 | 100.000 |
|  |  | other colour | 7.000 | 8 | 7 | 4 | 0.0778] | 0 |  | 0 |  |
|  | q12 | grey -> slate | 9841. | 0.05 | 0.05 | 0.00 | [0.0382, | 0.02 | 44.000 | 218000.00 | 99.980 |
|  |  |  | 000 | 8 | 7 | 0 | 0.0778] | 0 |  | 0 |  |
|  | q13 | grey -> blue | 2142 | 0.00 | 0.00 | 0.00 | [0, | 99.8 | 217646.00 | 218000.00 | 0.162 |
|  |  |  | 5.000 | 0 | 0 | 0 | 0.0791] | 38 | 0 | 0 |  |
|  | q20 | slate -> any | 307.0 | 0.05 | 0.05 | 0.00 | [0, 0.075] | 8.02 | 17484.000 | 218000.00 | 91.980 |
|  |  | other colour | 00 | 3 | 6 | 0 |  | 0 |  | 0 |  |
|  | q21 | slate -> grey | 1735. | 0.05 | 0.05 | 0.00 | [0.0365, | 1.58 | 3453.000 | 218000.00 | 98.416 |
|  |  |  | 000 | 7 | 7 | 0 | 0.0809] | 4 |  | 0 |  |
|  | q23 | slate -> blue | 362.0 | 0.05 | 0.05 | 0.00 | [0, 0.075] | 6.79 | 14802.000 | 218000.00 | 93.210 |
|  |  |  | 00 | 4 | 6 | 0 |  | 0 |  | 0 |  |
|  | q30 | blue -> any | 8623. | 0.05 | 0.05 | 0.05 | [0.0382, | 0.00 | 0.000 | 218000.00 | 100.000 |
|  |  | other colour | 000 | 8 | 7 | 4 | 0.0779] | 0 |  | 0 |  |
|  | q31 | blue -> grey | 250.0 | 0.00 | 0.00 | 0.00 | [0, | 92.6 | 201972.00 | 218000.00 | 7.352 |
|  |  |  | 00 | 4 | 0 | 0 | 0.0494] | 48 | 0 | 0 |  |
|  | q32 | blue -> slate | 1639. | 0.00 | 0.00 | 0.00 | [0, 0] | 98.4 | 214668.00 | 218000.00 | 1.528 |

|  |  |  | 000 | 1 | 0 | 0 |  | 72 | 0 | 0 |  |
| --- | --- | --- | --- | --- | --- | --- | --- | --- | --- | --- | --- |
| fem | q01 | any other | 1593 | 0.05 | 0.05 | 0.05 | [0.038, | 0.00 | 0.000 | 218000.00 | 100.000 |
| ale_ |  | colour -> grey | 0.000 | 8 | 7 | 2 | 0.0776] | 0 |  | 0 |  |
| 3 | q02 | any other | 1388 | 0.00 | 0.00 | 0.00 | [0, 0] | 98.3 | 214401.00 | 218000.00 | 1.651 |
|  |  | colour -> slate | 00.00 | 1 | 0 | 0 |  | 49 | 0 | 0 |  |
|  |  |  | 0 |  |  |  |  |  |  |  |  |
|  | q03 | any other | 6261 | 0.00 | 0.00 | 0.00 | [0, 0] | 99.9 | 217792.00 | 218000.00 | 0.095 |
|  |  | colour -> blue | 1.000 | 0 | 0 | 0 |  | 05 | 0 | 0 |  |
|  | q10 | grey -> any | 1615 | 0.05 | 0.05 | 0.05 | [0.0381, | 0.00 | 0.000 | 218000.00 | 100.000 |
|  |  | other colour | 8.000 | 8 | 7 | 2 | 0.0776] | 0 |  | 0 |  |
|  | q12 | grey -> slate | 1154 | 0.05 | 0.05 | 0.05 | [0.038, | 0.00 | 1.000 | 218000.00 | 100.000 |
|  |  |  | 9.000 | 7 | 8 | 2 | 0.0776] | 0 |  | 0 |  |
|  | q13 | grey -> blue | 3869 | 0.00 | 0.00 | 0.00 | [0, 0] | 99.8 | 217596.00 | 218000.00 | 0.185 |
|  |  |  | 4.000 | 0 | 0 | 0 |  | 15 | 0 | 0 |  |
|  | q20 | slate -> any | 453.0 | 0.05 | 0.05 | 0.00 | [0, | 7.93 | 17300.000 | 218000.00 | 92.064 |
|  |  | other colour | 00 | 4 | 7 | 0 | 0.0751] | 6 |  | 0 |  |
|  | q21 | slate -> grey | 2673. | 0.05 | 0.05 | 0.00 | [0.0371, | 1.67 | 3659.000 | 218000.00 | 98.322 |
|  |  |  | 000 | 7 | 7 | 0 | 0.0804] | 8 |  | 0 |  |
|  | q23 | slate -> blue | 508.0 | 0.05 | 0.05 | 0.00 | [0, | 6.94 | 15149.000 | 218000.00 | 93.051 |
|  |  |  | 00 | 4 | 7 | 0 | 0.0751] | 9 |  | 0 |  |
|  | q30 | blue -> any | 1042 | 0.05 | 0.05 | 0.00 | [0.0379, | 0.03 | 78.000 | 218000.00 | 99.964 |
|  |  | other colour | 8.000 | 8 | 7 | 0 | 0.0776] | 6 |  | 0 |  |
|  | q31 | blue -> grey | 336.0 | 0.00 | 0.00 | 0.00 | [0, | 92.4 | 201494.00 | 218000.00 | 7.572 |
|  |  |  | 00 | 4 | 0 | 0 | 0.0472] | 28 | 0 | 0 |  |
|  | q32 | blue -> slate | 2066. | 0.00 | 0.00 | 0.00 | [0, 0] | 98.6 | 215112.00 | 218000.00 | 1.325 |
|  |  |  | 000 | 1 | 0 | 0 |  | 75 | 0 | 0 |  |

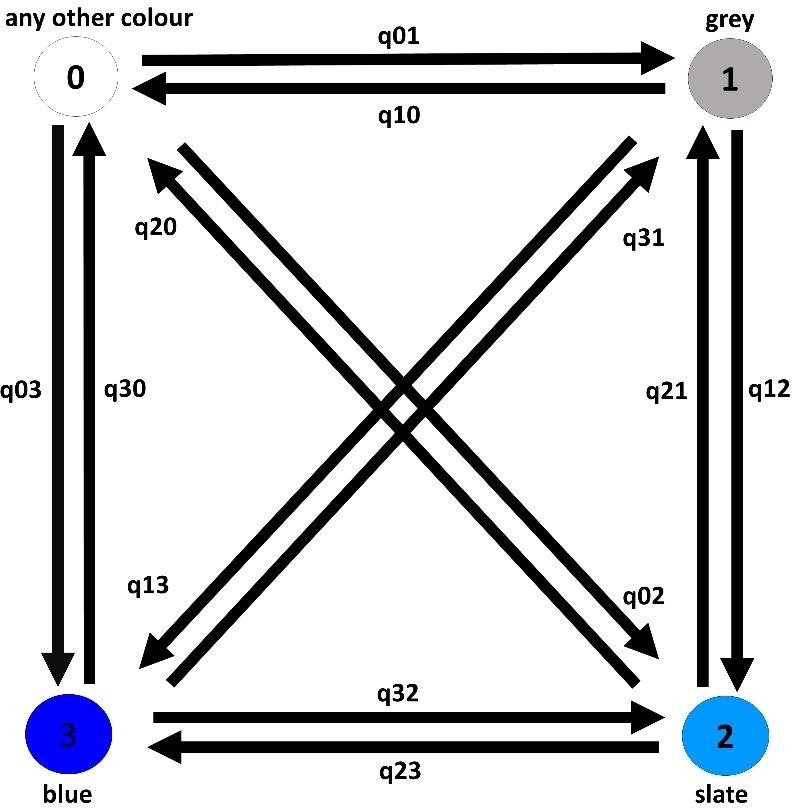

### Figure S1.

(q01: any other colour -> grey, q02: any other colour -> slate, q03: any other colour -> blue, q10: grey -> any other colour, q12: grey -> slate, q13: grey -> blue, q20: slate -> any other colour, q21: slate -> grey, q23: slate -> blue, q30: blue -> any other colour, q31: blue -> grey, q32: blue -> slate)
